# EMBER: Contextualized gene representations enable compositional prediction across cellular perturbations

**DOI:** 10.64898/2026.10.28.762655

**Authors:** Nicholas Ho, Eric P. Xing, Jian Ma

**Affiliations:** Ray and Stephanie Lane Computational Biology Department, Kenneth C. Griffin School of Computer Science, Carnegie Mellon University, Pittsburgh, PA 15213, USA; Machine Learning Department, Kenneth C. Griffin School of Computer Science, Carnegie Mellon University, Pittsburgh, PA 15213, USA

## Abstract

Predicting the effects of individual and combined perturbations is central to identifying context-specific vulnerabilities and designing targeted interventions. Yet learning gene representations that integrate functional relationships with cellular state and transfer across perturbation tasks remains challenging. Here, we present EMBER, a gene-centric foundation model that learns context-dependent functional representations of genes. EMBER combines interaction-guided contrastive learning with masked expression reconstruction during pretraining on a single-cell corpus curated for diversity and balanced sampling across cell types and experimental batches. EMBER decouples gene representation learning from task-specific prediction through a composable embedding framework: the pre-trained encoder remains frozen, while lightweight probes combine contextualized gene embeddings to predict individual and combined perturbation effects. The same pretrained representations support single-gene perturbation-response prediction, digenic interaction prediction, and multi-target drug-response prediction. EMBER improves cross-context perturbation-response prediction and genetic-interaction ranking relative to evaluated baselines. EMBER also prioritized *SCFD1*, a mediator of autophagosome–lysosome fusion, as a candidate dependency during *mTOR* inhibition. Together, these results establish contextualized gene embeddings as reusable building blocks for modeling genetic and pharmacological perturbations, supporting the prioritization of genetic interactions and context-specific interventions for experimental testing.

## Introduction

Gene functions are multifaceted, pleiotropic, and shaped by cellular state [1, 2]. Despite sharing essentially the same genome, cells within an individual exhibit distinct gene expression patterns, regulatory programs, and pathway activities across cellular states [3, 4]. As a result, gene identity alone is insufficient to determine the consequences of a perturbation. This context dependence is especially important in combinatorial genetic settings, where accurate prediction requires accounting for both the individual effects of perturbed genes and their context-dependent interactions [5, 6]. Predicting these effects at scale calls for gene representations that encode context-specific functional relationships and serve as reusable building blocks for modeling individual and combined perturbations.

Existing approaches only partially address this challenge. Representations derived from protein-protein interaction (PPI) networks [7] encode molecular relationships but are often static across contexts and limited in coverage, restricting their ability to represent context-specific gene function. Contextualized PPI methods such as PINNACLE [8] introduce cell-type-specific graph structure, but their representations are defined over the proteins and annotated cell types included in the constructed networks. Gene representations derived from text-embedding large language models [9, 10] have performed well on prior benchmarks [11, 12], but are typically static and limited by the incompleteness of the underlying knowledge sources. Single-cell foundation models (scFMs) [13–19] offer a complementary, data-driven route to learning representations from diverse cellular states. Although models such as Geneformer [13] and scGPT [14] learn gene-level features, many prominent applications emphasize cell-level tasks, including cell-type annotation, batch integration, and cell-state representation [20, 21]. However, strong performance on cell-level tasks does not necessarily translate into gene representations that generalize across gene-level prediction tasks. Evaluations have reported limitations in the downstream performance of representations extracted from scFMs [12, 20–24], and many applications rely on encoder fine-tuning [14, 17, 19, 25] or substantial task-specific adaptation [14, 15, 17]. These limitations motivate reusable gene representations that jointly capture functional relationships and cellular state.

Dedicated perturbation prediction models have made progress, but limited experimental coverage makes generalization to unseen perturbations and cellular contexts challenging, particularly for multi-gene effects. STATE [26] and STACK [27] address cross-context prediction using perturbation responses observed in other contexts, while GEARS [28] predicts unseen perturbations and their combinations within a given cellular context. More recently, PIE [29] has demonstrated joint generalization to unseen perturbations and cellular contexts for transcriptional-response prediction. Prior work has shown that simple predictors, including linear models using gene embeddings, can outperform dedicated perturbation models [12, 22]. These findings motivate a new focus on context-dependent gene representations that can be reused across tasks and composed to predict individual and combined perturbation effects across distinct experimental readouts without encoder retraining.

Here we present EMBER, a gene-centric foundation model that combines interaction-guided contrastive learning with masked expression reconstruction to learn context-dependent functional representations of genes. EMBER makes the gene, conditioned on cellular state, the primary unit of representation and downstream reuse. Its composable embedding framework separates representation learning from task-specific prediction: the pretrained encoder remains frozen, while lightweight downstream predictors combine gene embeddings to model perturbation responses and interactions involving individual genes and gene sets. We evaluate this shared interface across three tasks using distinct combinations of gene embeddings: a perturbed gene paired with a readout gene for perturbation-response prediction, two co-perturbed genes for digenic interaction prediction, and an unordered set of target genes combined with a readout gene for multi-target drug-response prediction. EMBER improves cross-context perturbation-response prediction and genetic-interaction ranking relative to evaluated gene-representation baselines. We further illustrate its utility for biological hypothesis generation by prioritizing *SCFD1* as a candidate dependency during *mTOR* inhibition. Together, these results establish contextualized gene embeddings as reusable building blocks for modeling genetic and pharmacological perturbations, supporting the prioritization of genetic interactions and context-specific perturbations for experimental testing.

## Results

### Overview of EMBER

EMBER is a gene-centric foundation model that provides reusable gene representations from single-cell transcriptomic data (**Fig. 1a**). Here, cellular context refers to the cellular state captured by single-cell transcriptomic measurements. By conditioning each gene representation on this state, EMBER is designed to capture context-specific functional relationships for reuse across downstream tasks.

**Figure 1:**
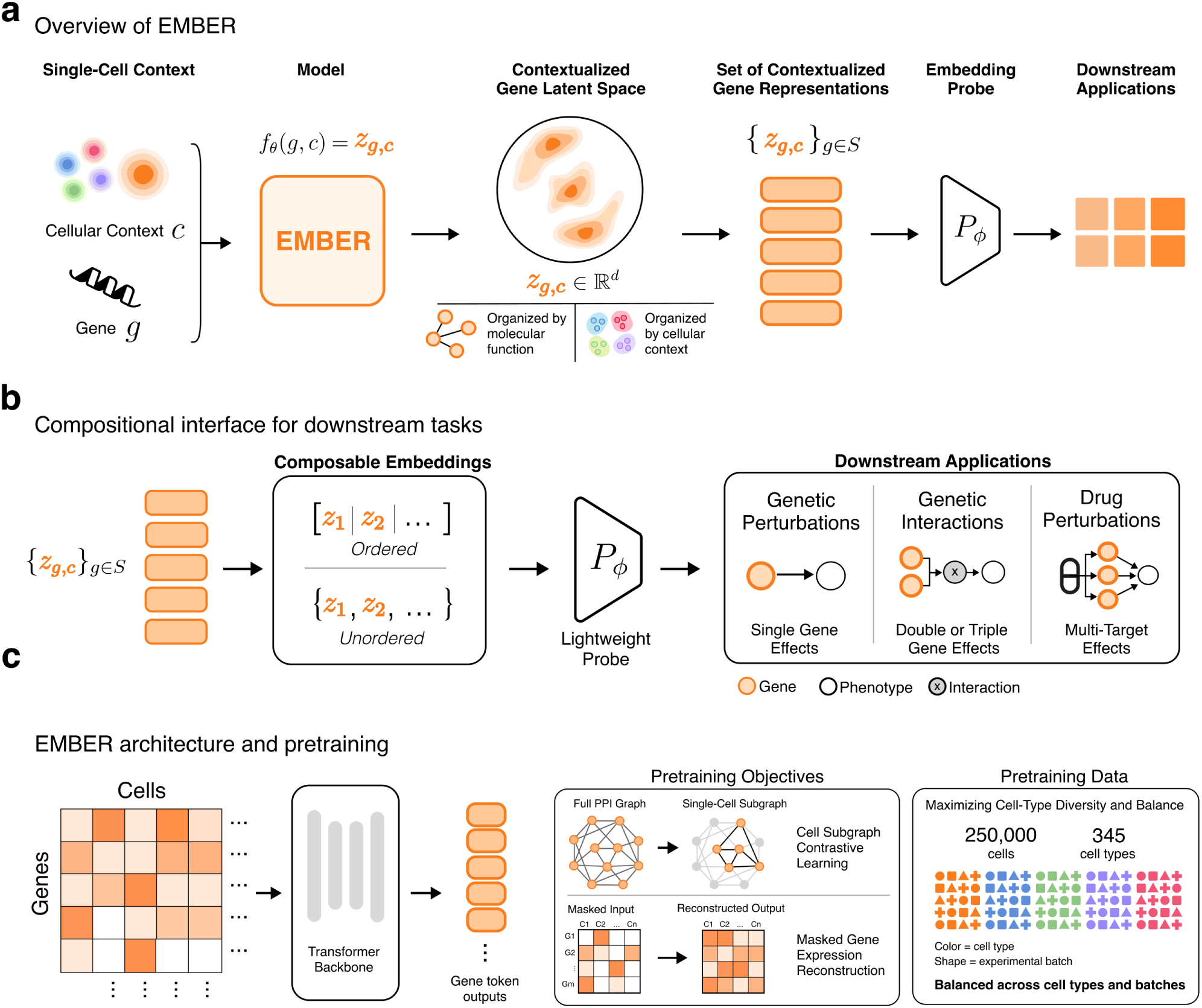
Overview of E_MBER_. a, High-level overview of E_MBER_. Given a gene and its single-cell context, E_MBER_ produces a context-specific gene embedding. E_MBER_ learns a contextualized gene latent space that jointly organizes genes by functional relationships and cellular context. These representations can be composed and passed to lightweight probes for downstream prediction. **b,** Composable interface for downstream applications. Depending on the task, embeddings can be combined as ordered inputs or permutation-invariant sets to support single-gene perturbation-response, genetic-interaction, and drug-response prediction. **c,** E_MBER_ architecture and pretraining. Gene-token representations are learned using two complementary pretraining objectives, cell subgraph contrastive learning (CSCL) and masked gene-expression reconstruction, on a pre-training corpus curated for cell-type diversity and approximately balanced sampling across cell types.

To capture diverse cellular contexts, we curated a pretraining corpus from CellxGene [30], prioritizing diversity across cell types and experimental batches while maintaining approximately balanced sampling across cell types. This design was motivated by evidence that pretraining data composition, rather than cell count alone, influences downstream performance [31–33]. The resulting CellxGene corpus contains 250,283 cells balanced across 345 cell types, with each primary cell type contributing between 255 and 1,048 cells (**Fig. 1c**). An additional 2,000 control cells from each of K562, Jurkat, HepG2, and RPE1 were also included in pretraining of the main model (**Methods**).

EMBER is pretrained on this corpus using two complementary objectives. Cell subgraph contrastive learning (CSCL) uses an annotated protein-protein interaction (PPI) network [7] to define contrastive training pairs within each cell. Positive pairs are annotated PPI edges between expressed genes in the cell’s tokenized context, contrasted against sampled pairs without annotated interactions (**Fig. 1c**). Because annotated PPIs are incomplete [34], this information is used as a contrastive training objective [35] rather than a fixed architectural constraint. In parallel, masked gene-expression reconstruction encourages the model to capture contextual expression relationships by reconstructing masked expression values from the remaining genes in the same cellular state. Joint training integrates an expression-conditioned interaction prior with transcriptional information in a shared representation space. Details of model pretraining are described in **Methods**.

After pretraining, the encoder is frozen. For downstream prediction, each gene’s embeddings are averaged across control cells within the relevant context. Each downstream task trains only a lightweight probe that composes frozen gene embeddings: a perturbed gene with a readout gene (EMBER-PERT), two co-perturbed genes (EMBER-GI), or a drug’s target-gene set with a readout gene (EMBER-DRUG) (**Fig. 1b**; Methods). In each task, we compare matched-context embeddings against static gene-identity embeddings and mismatched-context variants to assess the predictive utility of the encoder outputs and their alignment with the target context.

Together, this design makes context-dependent gene representations reusable across tasks, allowing lightweight predictors to model individual and combined perturbation effects without retraining the encoder.

### EMBER learns a unified latent space of functional relationships and cellular context

We first asked whether the resulting embedding space was organized along both functional and cellular axes. To examine the functional axis, we visualized a filtered set of Jurkat-contextualized gene embeddings using UMAP [36] (**Fig. 2c**). EMBER embedding clusters identified before projection were enriched for several Gene Ontology (GO) biological process, cellular component, and molecular function terms (**Fig. 2c**; **Supplementary Information**). We further calculated the adjusted Rand index (ARI) and normalized mutual information (NMI) to quantify recovery of GO annotations across four cell lines (**Fig. 2b** and **Fig. S1**). The context-agnostic STRING embeddings [12], trained using a graph neural network, achieved the highest clustering scores across GO categories (mean ARI 0.24, NMI 0.33). EMBER embeddings, while also encoding cellular context, recovered GO structure substantially better than the evaluated scFM baselines (ARI 0.20, NMI 0.27). Removing CSCL (EMBER (Recon-Only)) markedly reduced GO clustering (ARI 0.05, NMI 0.12), whereas the CSCL-Only ablation retained functional organization (ARI 0.20, NMI 0.28), identifying CSCL as the principal driver of functional organization.

**Figure 2:**
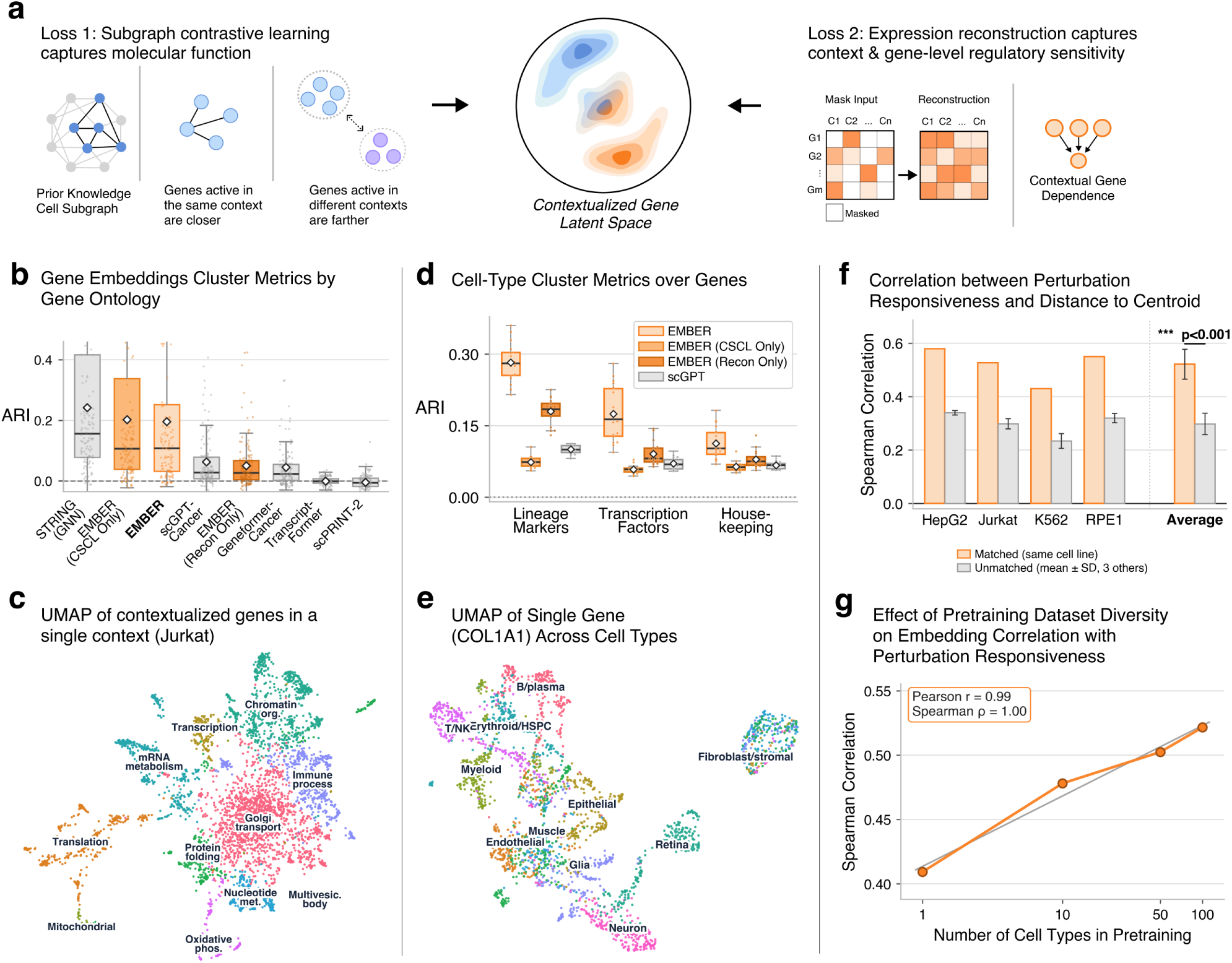
E_MBER_ pretraining learns a unified latent space of functional relationships and cellular context. a, Schematic of the two pretraining objectives. Cell subgraph contrastive learning introduces a context-conditioned PPI prior, while masked expression reconstruction learns transcriptional context. **b,** Adjusted Rand index (ARI) measuring recovery of functional annotations within 33 balanced Gene Ontology (GO) parent terms across Biological Process, Cellular Component, and Molecular Function categories. NMI and results stratified by GO category are shown in **Fig. S1**. **c,** UMAP of Jurkat-contextualized gene embeddings, colored by clusters identified before projection and labeled according to enriched GO terms. **d,** ARI measuring lineage recovery from contextualized embeddings of lineage markers, transcription factors, and housekeeping genes. Significance of comparisons with E_MBER_ was assessed with a two-sided paired Wilcoxon test; * *p <* 0.05, ** *p <* 0.01, and *** *p <* 0.001. **e,** UMAP of contextualized COL1A1 embeddings across cells from 11 lineages. Embeddings separate by lineage, with the fibroblast lineage most distinct. **f,** Spearman correlation between embedding distance from the gene centroid and experimentally measured *perturbation responsiveness*, comparing matched and mismatched cellular contexts using reconstruction-only embeddings. Significance was assessed with a two-sided paired *t*-test across n=4 cell lines; *** *p <* 0.001. **g,** Effect of cell-type diversity during pretraining, at a fixed total cell count, on the correlation between embedding centroid distance and *perturbation responsiveness* across four nested diversity tiers using reconstruction-only models.

Because STRING and GO draw partly on overlapping literature, GO recovery is not fully independent of the pretraining prior. To test whether EMBER can generalize beyond the PPI edges used during training, we withheld individual PPI edges or edges connecting genes within selected pathways during pretraining and evaluated recovery of the withheld edges in the contrastive projection space. Randomly held-out edges retained 88% of the mean reciprocal rank (MRR) and 90% of Hits@10 observed for training edges. The corresponding values for within-pathway edge holdouts were 61% and 71%, respectively, with both evaluations performing well above chance (**Fig.** S1c).

To examine the cellular-context axis, we visualized per-cell COL1A1 embeddings across 11 lineages (**Fig. 2e**). The UMAP showed embeddings separated by lineage, with a distinct fibroblast cluster consistent with COL1A1’s established role as a fibroblast marker. To quantify this pattern further, we analyzed 54 genes across three categories, 18 lineage markers, 18 transcription factors, and 18 housekeeping genes (**Fig. 2d**, **Methods**). Lineage recovery was strongest for marker genes (ARI 0.28, NMI 0.44), followed by transcription factors (ARI 0.17, NMI 0.28) and then housekeeping genes (ARI 0.11, NMI 0.20). EMBER outperformed scGPT across these groups (**Fig. 2d**), with additional baseline comparisons in **Fig. S1b**. Both single-objective ablations significantly reduced lineage recovery for marker genes, to ARI 0.18 and NMI 0.31 for EMBER (Recon-Only) and ARI 0.07 and NMI 0.16 for EMBER (CSCL-Only), supporting complementary contributions of CSCL and reconstruction to cellular contextualization.

We next asked whether masked reconstruction alone captures information associated with *perturbation responsiveness*. We defined a gene’s *perturbation responsiveness* as the fraction of assayed genetic perturbations that induced its differential expression in a given context using Perturb-seq data [37] (**Fig. S1d** and **Supplementary Information**). For this analysis, we used a reconstruction-only checkpoint pretrained on 100 cell types excluding lineages related to the four Perturb-seq cell lines (**Supplementary Information**). We found that the L2 distance between a gene’s embedding and the centroid of gene embeddings in a given cell line correlated positively with *perturbation responsiveness*, even after adjusting for expression magnitude, expression variance, and tokenization sampling frequency (**Fig. 2f** and **Supplementary Information**). Correlations were higher for matched than mismatched cell-line contexts (Spearman *ρ* = 0.522 vs. 0.298; *p* = 1.7*×*10*^−^*^4^, two-sided paired t-test across four cell lines).

We next examined whether this association depended on the diversity of the pretraining data. Holding the number of training cells fixed at 176,000, we pretrained reconstruction-only models on four nested subsets comprising 1, 10, 50, and 100 cell types (**Supplementary Information**). Across these four tiers, the mean correlation between embedding centroid distance and experimentally measured *perturbation responsiveness*, adjusted for the same covariates, increased monotonically with cell-type diversity (Spearman *ρ* = 1.00 across tiers) (**Fig. 2g**). Together, these analyses distinguish complementary contributions of the pretraining objectives: CSCL promotes functional neighborhood organization, whereas reconstruction captures transcriptional information associated with *perturbation responsiveness*. Joint training integrates these complementary signals into context-dependent gene representations that we next evaluate directly in perturbation prediction.

### EMBER predicts context-specific perturbation responses

Predicting transcriptional responses to genetic perturbations is central to understanding cell-state-specific gene regulation [38, 39]. We used the frozen EMBER model to embed the perturbed and readout genes in the relevant control-cell context, composed the ordered pair with a lightweight probe, and predicted whether the readout gene was differentially expressed (**Methods**).

To reduce predictive shortcuts based on average transcriptional response [40, 41], we used a response-aware negative-sampling strategy that matches positives and negatives by readout gene across perturbations (**Methods**). We evaluated three generalization regimes involving transfer to held-out perturbations (**Fig. 3c** and **Fig. S2c**), transfer to held-out cellular contexts (**Fig. 3b** and **Fig. S2b**), and transfer to held-out perturbations and contexts (**Fig. 3a** and **Fig. S2a**). Here, a held-out cellular context refers to a cell line excluded from probe training; its control cells were available for pretraining and embedding extraction, and a separate subset of its perturbation data was used for validation (**Methods**). We compared matched-context EMBER embeddings with static gene-identity embeddings and embeddings from a mismatched cell-type context. Additional baselines included PPI- and text-derived representations, scFM embeddings including scGPT [14] and Geneformer [13], sequence-derived embeddings including ESM2 [42], and perturbation models GEARS [28] and STATE [26]. All embedding-based methods were evaluated using the same prediction framework as EMBER (**Methods**).

**Figure 3:**
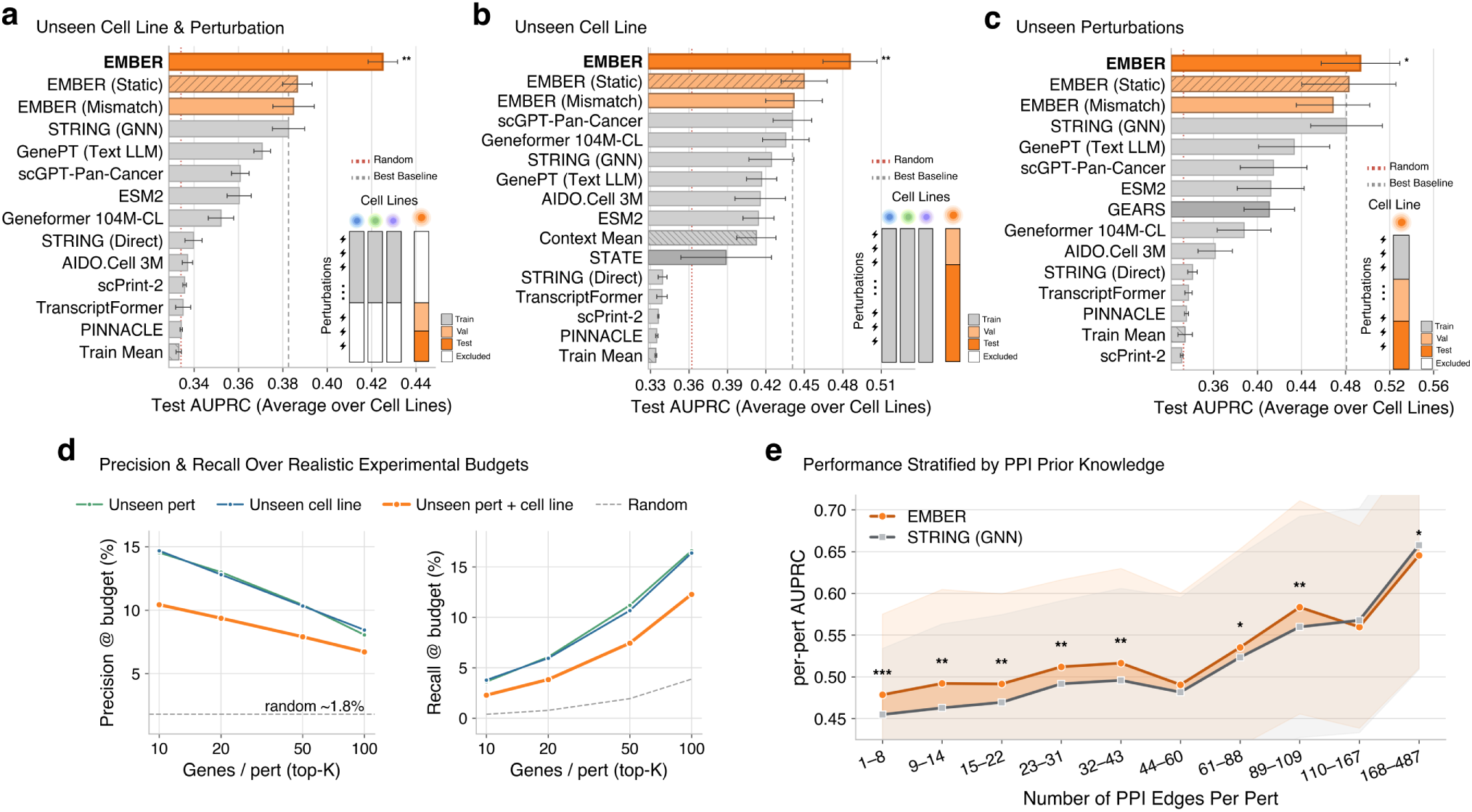
E_MBER_ predicts context-specific perturbation responses. a, Unseen cell line with unseen perturbations, the hardest generalization regime evaluated. Test AUPRC averaged over four cell lines for E_MBER_, its static and mismatched-context ablations, and external baselines, with the held-out cell line and a subset of its perturbations excluded from probe training. Bars show mean *±* s.d. across n=10 random splits, the dashed line marks the best external baseline, and the red dotted line marks chance. Significance compares E_MBER_ with the best external baseline by a two-sided paired t-test; * *p <* 0.05, ** *p <* 0.01, and *** *p <* 0.001. Results for individual cell lines are shown in **Fig. S2**. **b,** Unseen cell line with seen perturbations. Test AUPRC, shown as in panel **a**. **c,** Unseen perturbations within a single cell line. Test AUPRC, shown as in **a**. **d,** Precision and recall across top-K experimental budgets of 10, 20, 50, and 100 readout genes per perturbation, shown for three generalization regimes and averaged across cell lines. The dashed line indicates random prediction based on positive-label prevalence. **e,** Per-perturbation AUPRC stratified by the number of STRING PPI edges of the perturbed gene, comparing E_MBER_ and STRING (GNN). Shading shows variability across perturbations within each degree bin, and stars indicate per-bin significance of E_MBER_ relative to STRING (GNN); * *p <* 0.05, ** *p <* 0.01, and *** *p <* 0.001.

EMBER achieved the highest mean AUPRC and AUROC across both cross-context regimes, reaching an average AUPRC of 0.425 for the joint cell-line and perturbation holdout and 0.486 for held-out cell lines with seen perturbations across all four cell lines (**Fig. 3a,b** and **Fig. S2**). In the joint holdout, matched-context embeddings outperformed both the static (AUPRC 0.387) and mismatched-context (AUPRC 0.385) variants (**Fig. 3a**), supporting the predictive value of target-context information. STRING (GNN) embeddings were among the strongest external baselines in these regimes (AUPRC 0.383 for the joint holdout and 0.424 for cell-line-only holdout), consistent with earlier benchmarking [12]. By contrast, STRING (Direct) and STRING (Hop) performed near random (**Fig. S2**). When only the cellular context was held out, scGPT-Pan-Cancer [14] was the strongest scFM baseline (AUPRC 0.441), and the Context Mean predictor [26] was competitive with several scFM embeddings (AUPRC 0.413) (**Fig. 3b**). STATE [26] performed worse than EMBER (AUPRC 0.389), particularly in the Jurkat cell line (**Fig. S2**). These comparisons assess recovery of differentially expressed genes under the specified splits rather than overall transcriptomic prediction accuracy. For held-out perturbations within the same cell line, EMBER’s advantage was smaller (AUPRC 0.494 versus 0.481 for STRING (GNN)). EMBER remained competitive with STRING (GNN) across all four cell lines (**Fig. S2**). The strength of network-derived embeddings in this setting agrees with previous evaluations [12, 43].

We next evaluated performance under experimental prioritization budgets by using the full set of eligible negative readout genes rather than the 1:2 positive-to-negative ratio used above. Across budgets of 10 to 100 predicted genes per perturbation, EMBER maintained precision above the random baseline in all three regions. Recall increased with larger budgets (**Fig. 3d**). Holding out both context and perturbation reduced precision and recall relative to either holdout alone. We also assessed dependence on prior knowledge by stratifying performance by STRING connectivity. Performance generally increased with the number of annotated PPI edges. For held-out perturbations within the same context, EMBER and STRING (GNN) performed comparably for genes with many PPI edges, whereas EMBER showed larger gains for genes with fewer PPI edges (**Fig. 3e**). These results indicate that EMBER’s advantage extends to sparsely annotated genes.

Finally, we examined the individual contributions of the two pretraining objectives. Joint CSCL and reconstruction training outperformed either single-objective ablation in each regime (**Fig. S2**). The reconstruction-only model performed similarly to several scFM baselines in each regime (**Fig. S2**).

Together, these results show that matching reusable gene representations to cellular context improves perturbation-response prediction, with the clearest benefit in cross-cell-line transfer without encoder retraining.

### EMBER improves genetic interaction prediction

Genetic interactions reveal how genes act together within cellular pathways. In digenic interactions, two-gene perturbations produce effects that differ from expectations based on the corresponding single-gene perturbations [6, 44]. Such interactions can expose selective vulnerabilities associated with disease-related genetic alterations [45, 46]. We next applied the same frozen EMBER model to predict genetic interactions, combining the context-specific embeddings of the co-perturbed genes with a lightweight probe to predict the signed interaction score (**Fig. S3a**; **Methods**). We trained and evaluated the probe on the HAP1 genetic-interaction map [47], which measures interactions between 17,724 library genes and 222 query genes, holding out library genes while sharing query genes across training, validation and test sets. EMBER-GI achieved the highest mean AUPRC for detecting significant interactions among evaluated methods (0.181 against a chance level of 0.089), and STRING (GNN) was the strongest external baseline with an AUPRC of 0.156 (**Fig. 4a**). The AUPRC advantage held for both interaction types, with AUPRC values of 0.099 for negative and 0.112 for positive interactions and corresponding AUROC values of 0.545 and 0.568 (**Fig. S4**). Matched-context EMBER embeddings outperformed the static (AUPRC 0.161) and mismatched-context (AUPRC 0.174) variants.

**Figure 4:**
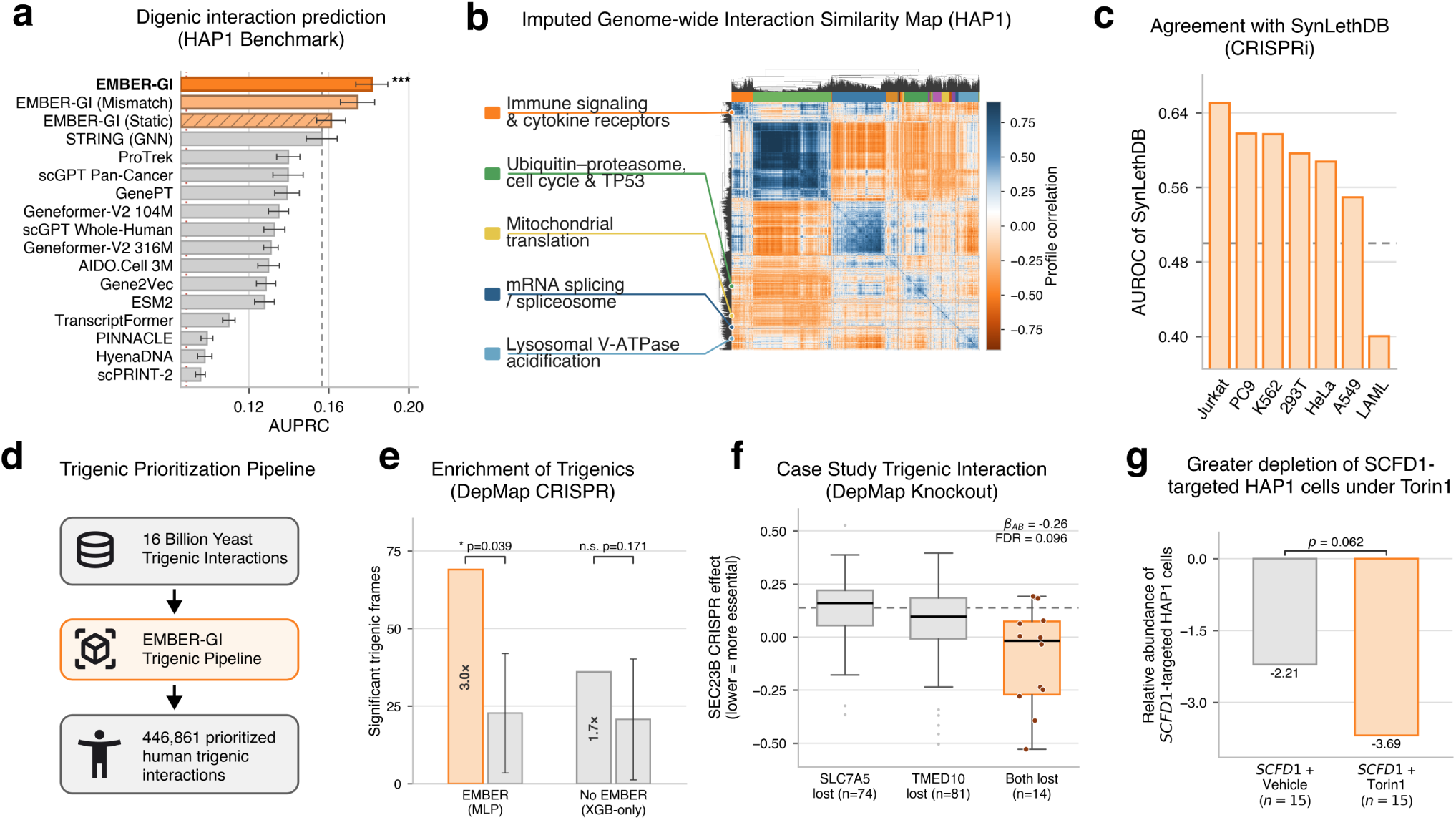
E_MBER_ improves genetic interaction prediction. a, Digenic interaction prediction on the HAP1 map from [47] for unseen library genes. Test AUPRC for E_MBER_-GI, its static and mismatched-context ablations, and evaluated baselines. Error bars show s.d. across n=10 random splits; significance was assessed with a two-sided paired t-test against the best external baseline; *** *p <* 0.001. **b,** Imputed genome-wide interaction similarity map over 16,408 genes, hierarchically clustered by interaction-profile similarity, with functional modules annotated. **c,** Agreement between the imputed map and CRISPRi synthetic-lethality screens in SynLethDB [48], stratified by cell line and reported as AUROC, with the dashed line indicating chance. **d,** Human trigenic candidate prioritization from 16 billion predicted yeast trigenic interactions, incorporating orthology, synthetic-lethality filtering, and E_MBER_-GI pairwise scores to yield 446,861 prioritized human candidate triplets. **e,** Enrichment of prioritized triplets in DepMap CRISPR dependencies relative to a 128-permutation null, comparing the E_MBER_ GI-residual pipeline with a size-matched E_MBER_-free XGBoost-only pipeline. The E_MBER_ pipeline yields a 3.0-fold enrichment of co-loss/readout configurations passing FDR *<* 0.20 relative to the null (*P* = 0.039), whereas the E_MBER_-free pipeline does not show significant enrichment (*p* = 0.171). **f,** Example prioritized triplet involving *SLC7A5*, *TMED10*, and *SEC23B*, showing increased SEC23B dependency associated with SLC7A5–TMED10 co-loss in DepMap cell lines. **g,** Greater depletion of *SCFD1*-targeted HAP1 cells under Torin1, although the difference did not reach statistical significance (*P* = 0.062).

We next used EMBER-GI to construct a genome-wide imputed genetic interaction map across 16,408 genes (**Fig. 4b**). Clustering the resulting interaction profiles revealed modules associated with signaling, DNA replication, and cell-cycle regulation. We compared interaction scores adjusted for gene-wise mean predictions with known synthetic lethal pairs from SynLethDB [48] using degree-matched negative pairs (**Fig. 4c** and **Fig. S3b**). Agreement was highest for CRISPRi screens and lowest for RNAi screens, with AUROC values of 0.60 and 0.45, respectively (**Fig. S3b**). Among CRISPRi datasets, concordance was highest for Jurkat, PC9, and K562 (AUROC 0.65, 0.62, and 0.62) and lowest for the acute myeloid leukemia dataset (AUROC 0.40) (**Fig. 4c**). The imputed map ranked several known synthetic lethal pairs within the top 0.013% of all predictions, including *KRAS*–*NDUFA12* [49], *MRPS5*–*TBCB* [50, 51], and *FAM50A*–*TRA2B* [51].

We next asked whether EMBER-GI could nominate candidate negative interaction partners of *MTOR*, a central regulator of cell growth [52]. Excluding pairs reported in SynLethDB, *SCFD1* ranked 14th among the remaining predicted *MTOR* interaction partners (**Supplementary Information**). Neither gene was a query in the HAP1 training map. *SCFD1* is a Sec1/Munc18-like protein that, upon mTORC1 inhibition, is recruited by dephosphorylated VAMP8 to promote STX17–SNAP29–VAMP8 SNARE assembly and autophagosome–lysosome fusion [53]. This mechanism suggested that mTOR inhibition might increase cellular dependence on *SCFD1*. Retrospective examination of two published CRISPR screens showed that *SCFD1*-targeted cells were more depleted under Torin1 than under vehicle, both in human HAP1 cells at day 18 (log2 fold change *−*3.69 versus *−*2.21, *P* = 0.062) [54] and in mouse EPP2 pancreatic cancer cells (*−*1.89 versus *−*1.47, *p* = 0.0046) [52] (**Fig. 4g**, **Fig. S3g-i** and **Supplementary Information**). Although the HAP1 comparison did not reach statistical significance, the direction of the effect was consistent with that observed in EPP2 cells. These observations support prioritizing *SCFD1* as a candidate dependency during *mTOR* inhibition, consistent with its role in autophagosome–lysosome fusion. EMBER ranks *SCFD1* and mTOR much higher (top 0.81%) compared to STRING (GNN) (top 1.90%) and its static counterpart (top 12.2%).

We then asked whether EMBER-GI could help prioritize candidate trigenic interactions in human cells (**Fig. 4d**). Trigenic interactions capture higher-order effects beyond the corresponding single-gene and pairwise perturbations [55, 56]. Although EMBER-GI scores pairwise interactions, trigenic effects frequently modify existing digenic interactions [55], and summed digenic scores correlated with trigenic scores in yeast (Pearson *r* = 0.507; Spearman *ρ* = 0.459; **Fig. S3c**). These observations motivated using pairwise interaction structure to prioritize trigenic candidates rather than directly predict higher-order effects.

Building on a cross-species interaction-mapping strategy [45], we developed a multi-stage pipeline that began with 16 billion predicted yeast trigenic interactions [56], mapped high-scoring candidates to human genes through orthology, and filtered triplets using an auxiliary synthetic-lethality classifier (**Methods**). We then used EMBER-GI to score the three pairwise interactions within each filtered triplet and ranked triplets by their summed pairwise residual scores, resulting in 446,861 prioritized candidate human triplets. These candidates were enriched for shared CORUM complex membership [57] and shared Reactome [58] and KEGG [59] pathway membership relative to the orthology-derived candidate pool, including SNARE-mediated membrane fusion [60] and capped pre-mRNA processing [61] (**Fig. S3d–f**).

We further assessed the prioritized triplets retrospectively using the Broad Institute Dependency Map (DepMap) [62], asking whether cancer cell lines with copy-number loss of two genes showed increased sensitivity to a CRISPR knockout of a third gene (**Methods**). This exploratory analysis yielded 69 co-loss/readout configurations with negative interaction coefficients at FDR*<* 0.20, a 3.0-fold enrichment over a 128-permutation null (*P* = 0.039, **Fig. 4e** and **Supplementary Information**). The size-matched pipeline without EMBER did not show significant enrichment relative to its null (*P* = 0.171). For example, combined copy-number loss of *SLC7A5* and *TMED10* was associated with increased *SEC23B* dependency (co-loss group, n=14; *β_AB_* = *−*0.258, FDR= 0.096; **Fig. 4f**). We hypothesize that reduced SLC7A5-mediated essential amino-acid uptake [63] and impaired export of TMED10-dependent cargoes [64] jointly sensitize cells to disruption of ER-to-Golgi transport through loss of the COPII subunit SEC23B [65].

Together, these analyses extend frozen gene representations from single-gene perturbation prediction to digenic interaction ranking and hypothesis-driven prioritization of higher-order candidates. The proposed genetic interactions and mechanisms require direct experimental testing.

### EMBER enhances drug-specific transcriptional response prediction

Many drugs engage multiple protein targets, whose effects vary across cellular contexts [66–68]. We therefore asked whether EMBER embeddings could represent drugs as sets of contextualized target genes to predict drug-induced transcriptional responses. For each drug, we obtained known target genes from DrugBank [69] and computed matched EMBER embeddings in the relevant cellular context. EMBER-DRUG pools these embeddings and target-specific mechanism-of-action (MoA) annotations using an order-invariant probe and combines the resulting drug representation with a readout-gene embedding to predict its response (**Fig. S5** and **Methods**).

We evaluated generalization to unseen drugs on LINCS L1000 [70] and Tahoe-100M using drug-level holdouts within each cell line, keeping drugs with identical annotated target sets in the same split and comparing against alternative gene and drug-structure representations and the drug-response transformer model XPert [71]. A baseline assigning every held-out drug the mean response across training drugs outperformed all learned models under control-reference Pearson correlation (**Supplementary Information** & **Table S7**), consistent with reports that this metric can reward shared responses [40, 41]. To assess drug-specific variation, we used perturbed-reference Pearson correlation as the primary metric, removing each gene’s mean response across training drugs from both predicted and observed profiles before computing correlation [40] (**Methods**).

EMBER-DRUG achieved the best average rank across the evaluated cell lines in both LINCS L1000 and Tahoe-100M (**Fig. 5a,b** and **Fig. S5c,d**). Average perturbed-reference Pearson correlations were approximately 0.15 and 0.05, respectively (**Table S7**). Contextualized EMBER embeddings also outperformed the context-free static gene-identity variant, supporting the predictive utility of the contextualized encoder outputs for this task. Matched-context embeddings also ranked above the mismatched-context variant on average in both datasets (**Fig. S5c,d**). Replicate agreement under the same metric was lower in Tahoe-100M (r=0.15) than in LINCS L1000 (r=0.51), providing a reference for measurement reproducibility rather than an established prediction ceiling.

**Figure 5:**
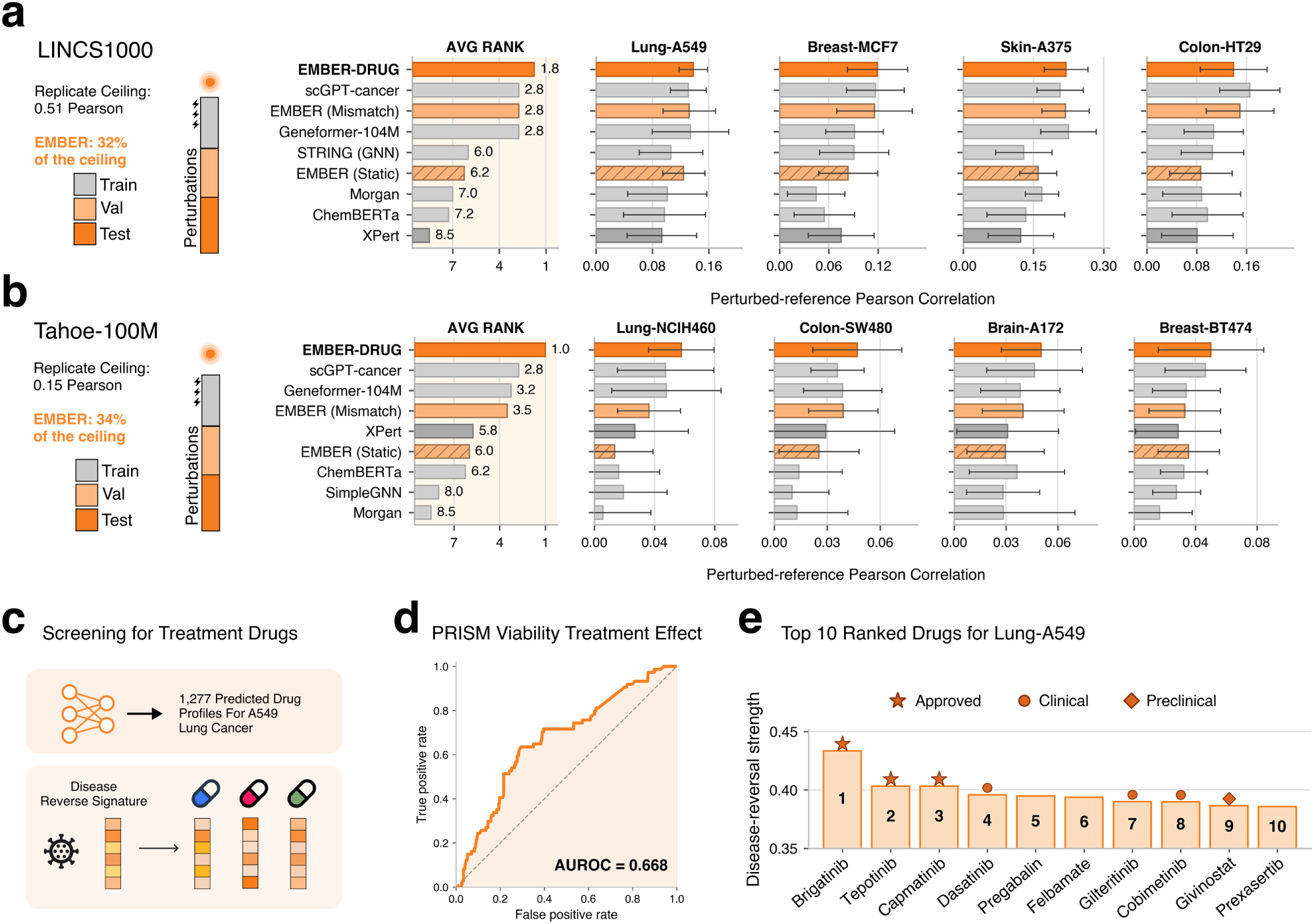
E_MBER_ enhances drug-specific transcriptional response prediction. a, Drug-response prediction on LINCS L1000 for held-out drugs, evaluated by perturbed-reference Pearson correlation. Results are shown as average method rank among displayed methods across four cell lines and prediction performance within each cell line. E_MBER_-DRUG is compared with gene embedding, drug-structure, and drug response baselines. Replicate agreement under the same metric was r=0.51. Additional baselines are shown in **Fig. S5** Error bars indicate s.d. across n=10 random splits. **b,** Corresponding benchmark on Tahoe-100M across four cell lines. Replicate agreement was r=0.15. Error bars are as in panel **a**. More baselines are shown in **Fig. S5**. **c,** Schematic of the disease-signature reversal screen. E_MBER_-DRUG predicts transcriptional responses for 1,277 drugs in A549 lung adenocarcinoma and ranks compounds by predicted reversal of a lung-cancer disease signature. **d,** Evaluation of predicted disease-signature reversal against A549 PRISM drug viability for 399 compounds (AUROC = 0.668). **e,** Top ten compounds ranked by predicted disease-signature reversal, with approved or clinically investigated lung-cancer therapies and compounds supported by prior lung-cancer studies highlighted.

We next asked whether the transcriptional responses predicted by EMBER-DRUG could support candidate prioritization by disease-signature reversal (**Fig. 5c**). We used EMBER-DRUG to predict transcriptional responses for 1,277 drugs and rank compounds by predicted reversal of the lung adenocarcinoma expression signature in A549 cells (**Fig. 5c**). Across 399 drugs with PRISM measurements, predicted reversal scores achieved an AUROC of 0.668 for identifying drugs with low A549 viability AUC (**Fig. 5d**, **Methods**). The top three were capmatinib and tepotinib, established therapies for MET exon 14-skipping NSCLC [72, 73], and brigatinib, an established therapy for ALK-positive NSCLC [74] (**Fig. 5e**). EMBER-DRUG uses A549’s single-cell expression context but does not explicitly encode the somatic alterations that define these approved indications. Other top-10 candidates included dasatinib (rank 4), which showed modest single-agent activity in a phase II NSCLC trial [75], gilteritinib (rank 7), which was associated with disease control for over 24 months in one patient with ALK-positive NSCLC after multiple ALK inhibitors [76], cobimetinib (rank 8), whose combination with atezolizumab produced responses in 5 of 28 patients with NSCLC in a phase Ib study [77], and givinostat (rank 9), which increased pemetrexed sensitivity in A549 cells and their pemetrexed-resistant derivatives in preclinical experiments [78]. Overall, the PRISM viability association and recovery of compounds previously studied in lung cancer provide complementary retrospective support for candidate prioritization, but do not establish new indications or therapeutic efficacy in the modeled cellular context.

Together, these results extend the same frozen gene-representation framework from genetic to pharmacological perturbations, showing that contextualized target-gene representations can support drug-specific transcriptional prediction and candidate prioritization without encoder retraining.

## Discussion

In this work, we developed EMBER, a contextualized gene foundation model that addresses the challenge of learning gene representations that integrate functional relationships with cellular context. Frozen EMBER embeddings composed with downstream probes improve cross-context perturbation-response prediction and genetic-interaction ranking, while also supporting drug-response modeling without task-specific encoder retraining. EMBER also nominated *SCFD1* as a candidate dependency during *mTOR* inhibition, consistent with retrospective evidence from CRISPR screens. These results support a gene-centric pretraining strategy in which functional knowledge and cellular state are integrated into representations that can be reused and composed across biological tasks. The ablations suggest complementary contributions from the interaction prior and masked reconstruction, while the fixed-cell-count comparisons highlight the importance of cellular diversity during pretraining. Together, these findings motivate attention to pretraining objectives and data composition alongside increases in corpus size.

EMBER has several limitations. First, it currently contextualizes representations from transcriptomic measurements alone. Transcription-factor activity, chromatin accessibility, protein abundance, post-translational regulation and spatial microenvironment can all shape gene function without being fully reflected in gene-expression measurements. Integrating genotype information with epigenomic, proteomic, and spatial measurements could broaden the definition of cellular context and distinguish regulatory states that are not resolved by transcriptomics alone. EMBER also depends on an incomplete, unevenly annotated interaction prior. Although its gains extend to sparsely annotated genes, this does not establish independence from prior-knowledge biases. Evaluating alternative interaction priors and transfer to underrepresented genes and cellular states will be important for assessing the robustness of the learned representations.

Second, performance gains remain task- and metric-dependent, and substantial room remains for improving predictive performance. For perturbation prediction, the main benchmark concerns essential-gene CRISPR interference in cultured cell lines and evaluates differential-expression responses rather than complete transcriptomic profiles. Target-context perturbations also inform validation, so cross-context evaluation is not entirely target-label-free. Testing primary cells, broader perturbation classes, and contexts excluded from both pretraining and model selection would better establish generalizability. For drug perturbations, the benchmarks assess held-out drugs within each cell line, rather than transfer to unseen cell types, and depend on available target and mechanism-of-action annotations. Measurement variability and limited perturbation coverage further constrain prediction. Additional replicated measurements and calibrated uncertainty estimates could help identify predictions reliable enough for experimental prioritization.

Third, compositional prediction does not itself establish higher-order genetic mechanisms. The trigenic pipeline aggregates pairwise predictions to prioritize candidates, and retrospective co-loss associations do not replace controlled three-gene perturbation experiments. Predicted disease-signature reversal and its retrospective association with PRISM viability support compound prioritization rather than establishing therapeutic efficacy. Prospective validation using data independent of candidate selection, with appropriate single- and pairwise-perturbation controls, is needed to test the proposed interactions and their context specificity. Such experiments could also provide training data for models that directly learn higher-order effects rather than relying on pairwise surrogates.

Our findings also highlight an important direction for single-cell foundation models. Many current evaluations emphasize cell-level tasks, such as cell-type classification, clustering, batch integration, and cell-state representation. Although these tasks are important, they do not directly test whether a model has learned useful representations of genes. By centering context-dependent gene embeddings as the primary learned object, EMBER provides a practical framework for gene-focused prediction without requiring encoder fine-tuning for each downstream task. Evaluating frozen representations across distinct tasks offers a complementary test of whether pretraining captures transferable functional information. More broadly, EMBER provides a gene-centric interface between observational single-cell data and perturbation experiments. Coupling reusable representations with prospective validation could help prioritize experiments that resolve context-dependent gene function and uncover genetic and pharmacological vulnerabilities.

## Methods

### Pretraining EMBER

#### Architecture and tokenization

Each cell was tokenized into 1,280 gene slots, 80% sampled uniformly at random from its expressed genes and 20% from genes with zero counts. Counts were library-size normalized, log1p-transformed and kept continuous. The input token for gene *i* summed a learned gene-identity embedding and a two-layer MLP embedding (width 128, GELU) of its scalar expression value *e_i_*,

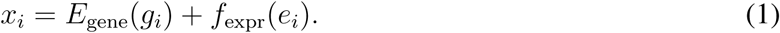

Tokens were processed by a 6-layer pre-normalization transformer encoder with embedding dimension 128, 4 attention heads, a feed-forward hidden dimension of 512, GELU activations and dropout of 0.1. The encoder output *z_i_* is the context-dependent embedding of gene *i*.

#### Pretraining objectives

For masked gene-expression reconstruction, we masked 50% of the gene tokens in each cell by replacing the expression embedding with a learned mask token while keeping the gene-identity embedding visible. A two-layer MLP decoder predicted the expression ê*_i_* of each masked gene, and the loss was the mean-squared error over the masked set 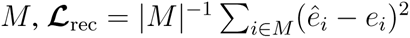

For cell subgraph contrastive learning (CSCL), each *z_i_* was passed through a two-layer projection head and L2-normalized to give *p_i_*. For every cell, we restricted the STRING v12 PPI graph [7] to edges between expressed genes in the tokenized context of that cell. The graph retained edges with a combined score above 0.8, and all edges were kept for genes with fewer than 20 edges, yielding 167,855 interactions among 18,359 genes. Each positive edge between anchor gene *a* and partner gene *b* was contrasted against *R* = 5 sampled partners ***N*** (*a*) that excluded true edges and self-loops, using cosine similarity with inverse temperature *α* = 5,

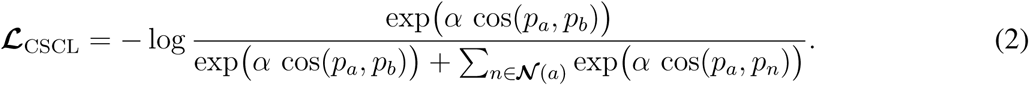

The two objectives were trained jointly as ***L*** = ***L***_rec_ + *λ **L***_CSCL_ with *λ* = 3.0.

#### Optimization

We trained with AdamW (learning rate 1 *×* 10*^−^*^3^, weight decay 0.01), a linear warmup over 1,500 steps followed by cosine decay to 1 *×* 10*^−^*^6^, batch size 64, no gradient accumulation and gradient-norm clipping at 1.0. Approximately 1% of cells were held out for validation. We trained for up to 100 epochs with early stopping. Pretraining ran on a single NVIDIA A6000 GPU for 24 h. The CSCL-Only and Recon-Only ablations followed the same recipe with one objective removed. We selected the pretraining checkpoint based on probe performance on the fold-1 validation sets of the unseen-cell-line- and-perturbation and HAP1 genetic-interaction benchmarks. This checkpoint was then fixed for every EMBER evaluation in the paper. Pretraining and probe settings are listed in **Table** S2.

#### Pretraining corpus

We assembled the corpus from human cells in the CellxGene Census (version 2025-11-08) [30], with quality filters similar to those of scTab [79]. We retained primary data from 10x assays and curated cell types against the Cell Ontology (**Supplementary Information**). We then sampled an equal quota of cells per cell type, keeping smaller cell types in full and drawing across batches within each type, which gave a base corpus of 250,283 cells from 345 cell types. Raw UMI counts were restricted to protein-coding genes, and we added 2,000 control cells from each of K562, Jurkat, HepG2 and RPE1.

### Contextual embeddings and probes

The pretrained encoder was frozen for every downstream task. To embed gene *g* in context *T*, we ran unperturbed control cells of *T* through the encoder and averaged the output *z_g_* over cells to give 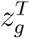 . The static variant used the learned gene-identity embedding *E*_gene_(*g*) taken before the transformer, which carries no cellular context. The mismatched-context variant replaced 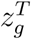 with the embedding of the same gene in a CellxGene cell type whose expression profile had a Pearson correlation of approximately 0.5 with the target cell line (**Supplementary Information** and **Table** S4). Probes composed ordered gene pairs by concatenation followed by an MLP, and unordered gene sets by a sum-pooled set encoder [80]. Only probe weights were trained (**Table** S2).

### Embedding space analyses

All analyses used frozen embeddings and are detailed in the **Supplementary Information**. Recovery of Gene Ontology (GO) structure was scored by the adjusted Rand index (ARI) and normalized mutual information (NMI) after clustering gene embeddings within 33 balanced GO parent terms in each of HepG2, Jurkat, K562 and RPE1 (**Table** S3). Lineage recovery was scored by ARI and NMI on per-cell embeddings of 54 genes (18 lineage markers, 18 transcription factors and 18 housekeeping genes) across 2,816 cells from 11 lineages.

### Perturbation-response prediction

#### Task and labels

We predicted whether perturbing gene *p* induces differential expression of readout gene *g* in a given cell line, with binary labels

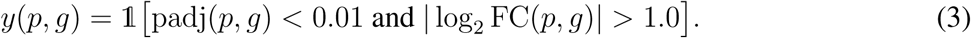

Labels came from pseudobulk DESeq2 [81] analysis of genome-scale essential-gene CRISPRi Perturb-seq in HepG2, Jurkat, K562 and RPE1, following the source study [37] and current recommendations for single-cell differential expression [82].

#### Negative sampling

Each positive was paired with two negatives that shared its readout gene but came from other perturbations with adjusted *P >* 0.1, yielding a 1:2 positive-to-negative sampling ratio (**Fig.** S2d). Readout genes too frequent to support this ratio were dropped. We kept perturbations with at least five positives and capped each at the 100 positives with the largest *|* log_2_ FC*|*. The Train Mean baseline, which predicts from the training frequency of the readout gene alone, served as the gene-frequency control (**Table** S5).

#### Generalization splits

We evaluated three regimes on the same curated pairs, each with 10 random partitions. In the unseen-perturbation regime, one third of the perturbations in a cell line were used for training and the remaining two thirds were split 30/70 into validation and test. In the unseen-cell-line regime, one cell line was held out and its perturbations were split one third for validation and two thirds for testing, while the same perturbations could appear in the training cell lines. In the unseen-cell-line- and-perturbation regime, 15% of the perturbations in the held-out cell line went to validation and 25% of the remainder to testing, and both sets were excluded from every training cell line.

#### EMBER-PERT and embedding baselines

EMBER-PERT concatenated the embeddings of the perturbed and readout genes, computed from control cells of the relevant cell line, and passed them to a two-layer MLP of width 128 that output a differential-expression probability. The probe was trained with binary cross-entropy and AdamW for up to 200 epochs with early stopping, and the configuration with the highest validation AUROC over a 24-combination grid was evaluated once on the test set (**Table** S2). Every embedding baseline used the same probe and procedure after reduction to 128 dimensions by PCA, and single-cell foundation model embeddings were extracted from control cells and mean-pooled per cell line as for EMBER (**Table** S1). Pairs missing either embedding were excluded from training and assigned the training base rate at evaluation.

#### Perturbation model baselines

GEARS [28] was evaluated in the within-cell-line regime and STATE [26] in the unseen-cell-line regime, both trained with control and perturbed cells paired within batch. We tuned the learning rate over *{*1 *×* 10*^−^*^3^, 1 *×* 10*^−^*^4^*}* and the batch size over *{*64, 128*}* and otherwise followed the notebooks provided by the authors, including the STATE notebook for this dataset. Predicted profiles were converted to differentially expressed gene calls with either DESeq2 on replicate-aware pseudobulks or the Wilcoxon test, both with FDR correction (**Supplementary Information**). The calling method was chosen for each model using validation performance, which selected DESeq2 for GEARS and the Wilcoxon test for STATE, and all scores are reported in **Table** S5.

#### Evaluation

Performance was measured by AUROC and AUPRC, reported as the mean and s.d. across the 10 partitions for each cell line. We also computed both metrics within each perturbation and averaged them (**Table** S5). Given limited sample sizes, we used 10 random partitions to reduce sensitivity to any single split; variation across these overlapping partitions reflects split variability, not independent biological replication. For the budget analysis, the sampled negatives were replaced by the full pool of eligible negatives in the first three partitions, and precision and recall were computed for the top 10, 20, 50 and 100 ranked readout genes per perturbation. Per-perturbation AUPRC was also stratified by the STRING degree of the perturbed gene (**Supplementary Information**). EMBER-PERT was compared with the best non-random external baseline by two-sided paired *t*-test across partitions.

### Genetic interaction prediction

#### Dataset and splits

We used the HAP1 CRISPR genetic-interaction map of Billmann et al. [47], which scores 222 query genes against 17,724 library genes by a quantitative genetic-interaction value (qGI) with an associated FDR. Library genes were split 80/10/10 into training, validation and test sets, each gene pair inherited the split of its library gene, and query genes were shared across splits. The split was repeated 10 times using independent random partitions of library genes.

#### EMBER-GI and baselines

EMBER-GI concatenated the library-gene and query-gene embeddings and predicted the signed qGI with a four-layer MLP of width 256 (ReLU, no dropout), trained with mean-squared error and Adam for up to 200 epochs. Embeddings were contextualized on 2,000 sampled HAP1 control cells [83]. Among four combinations of learning rate and weight decay, we selected the model with the highest validation AUPRC for significant interactions (**Table** S2). Baselines were the gene embeddings used for perturbation prediction, passed through the same probe.

#### Evaluation

Predictions were adjusted for gene-wise mean effects to obtain pair-specific residual scores,

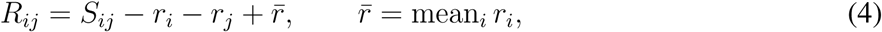

where *S_ij_* is the predicted score and *r_i_* is the mean predicted score across pairs containing gene *i*. Significant interactions were pairs with *|*qGI*| >* 0.3 and FDR *<* 0.1, divided into negative (qGI *< −*0.3) and positive (qGI *>* 0.3) interactions, and non-interacting pairs had *|*qGI*| <* 0.05 and FDR *>* 0.5. We ranked pairs by *−R_ij_*, *R_ij_* and *|R_ij_|* for negative, positive and any interaction, respectively. We report AUPRC at natural prevalence and AUROC on balanced sets of query-anchored, library-degree-matched negatives averaged over 20 draws. Pearson correlation, Spearman correlation and mean-squared error were computed on the signed qGI (**Fig.** S4 and **Table** S6).

#### Genome-wide imputation

For imputation, we trained one EMBER-GI model on signed qGI with an 85/15 split of query genes, with library genes shared, and selected it by validation Spearman correlation. We scored all ordered pairs of the 16,408 library genes to obtain a matrix *M* and symmetrized it as 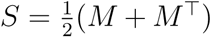, leaving the diagonal undefined. Residuals *R_ij_* were computed from *S* with *r_i_* defined as the row mean of gene *i*.

#### Synthetic lethality benchmark

We compared the residuals with synthetic-lethal pairs from Syn-LethDB [84], collapsed to unique unordered pairs with both genes in the imputed map and restricted to pairs with experimental evidence. Pairs were scored by *−R_ij_*. AUROC was computed against an equal number of degree-matched negative pairs, overall and stratified by evidence type and cell line.

#### Trigenic candidate prioritization

We mapped the top 5% of DANGO-predicted yeast triplets [56] to human genes using many-to-many InParanoid orthology [85], with a shared-domain fallback for genes lacking direct orthologs. We expanded all combinations of human orthologs, sorted genes within each triplet and retained the 46,035,705 triplets fully covered by the EMBER HAP1 embeddings. Candidates were pre-filtered with a gradient-boosted digenic synthetic-lethality classifier trained on SynLethDB [48] and Costanzo et al. [44], using GO Resnik similarities for each GO branch and dot-product similarities of ESM2 and STRING (GNN) embeddings as features (**Supplementary Information**). Retaining the top 15% by summed pair scores left 5,627,852 triplets. We ranked these by the sum of the three EMBER-GI residuals *R_ij_* and retained the top 10%, which gave 446,861 candidate human trigenic interactions. Triplets containing genes that could not be transferred were dropped, so the retained sets are smaller than the nominal fractions. As a comparator without EMBER, we omitted the EMBER-GI ranking step and selected the same number of final candidates (446,861) by the summed classifier scores alone. All other steps were identical.

#### Pathway enrichment

We tested the 446,861 prioritized triplets against the background of all 46,035,705 mapped triplets. We used Reactome [58] and KEGG [59] pathways of 3 to 500 genes and CORUM complexes [57], and counted a triplet as a hit only when all three genes belonged to the gene set. Enrichment was the observed hit count divided by its expectation under the background rate, with one-sided hypergeometric tests and Benjamini-Hochberg correction (FDR *<* 0.05). An empirical null came from 100 random foreground sets of equal size drawn from the same background.

#### DepMap co-loss analysis

We tested prioritized triplets using DepMap 24Q4 Public copy-number and CRISPR gene-effect data [62] from 1,178 cell lines. Each triplet yielded up to three frames, with one gene as the CRISPR readout and the other two as the co-lost pair. Eligible frames required genes on distinct chromosome arms, sufficient representation of all four loss groups and co-loss across at least two tissue lineages. We centered gene-effect values within tissue lineages and residualized them against readout copy number and genome-wide loss burden, then fitted the two loss indicators and their interaction to these residualized outcomes. A frame was considered significant when the interaction coefficient was negative at Benjamini-Hochberg FDR *<* 0.20 across all tested frames, and the number of significant frames was compared with 128 permutations of the loss-matrix rows (**Supplementary Information**).

### Drug-response prediction

#### Datasets

We evaluated on LINCS L1000 [70] and Tahoe-100M [86], in both cases by predicting a gene-level transcriptional response for held-out drugs within a cell line. Drug targets and the mechanism of action of each target came from high-confidence DrugBank annotations [69], and a drug was retained if it had at least one target and at least half of its targets were embedded.

For LINCS, we used level-5 moderated z-score signatures at 10 *µ*M and 24 h, averaged per drug and cell line, over 964 landmark genes and approximately 540 drugs. We used the four cell lines with the most mechanism-annotated drugs and an available single-cell control dataset, which were A375, A549, HT29 and MCF7. The level-5 moderated z-score was used directly as the prediction target.

For Tahoe, we used the pseudobulk log_2_ fold changes released with the dataset (Hugging Face repository). Because agreement between plates was low, we kept the plate with the most cells, which gave a median of 2,330 and a mean of 2,970 cells per perturbation. Four cell lines were selected for reproducible and distinct drug-specific responses across tissues, NCIH460, SW480, A172 and BT474 (**Supplementary Information**). The gene panel was 1,024 highly variable genes computed on control cells and restricted to embedded genes, and the resulting dataset contained 172 drugs.

#### Cellular contexts

Gene embeddings for each LINCS cell line were contextualized on approximately 2,000 control cells of the same line. We used vehicle-treated A549 and MCF7 cells from sci-Plex (GSE139944) [87], A375 cells from GSE164614 [88] and HT29 cells from GSE167262 [89]. Tahoe embeddings were contextualized on the DMSO-treated control cells of each Tahoe cell line.

#### EMBER-DRUG

EMBER-DRUG represents a drug *d* by its set of at most 25 DrugBank targets ***T****_d_* and predicts one transcriptional-response value per drug and readout gene. Each target token concatenated the contextual embedding *z* ^ctx^ with a learned 8-dimensional mechanism-of-action embedding *m_t_* over three classes (inhibitory, activating or other). A shared two-layer encoder *ϕ* of width 64 was applied to every token, padded positions were masked and the outputs were sum-pooled into a permutation-invariant drug vector,

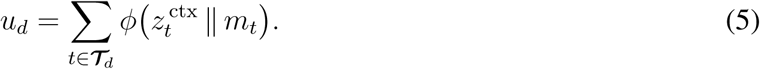

The readout-gene embedding was linearly projected, concatenated with *u_d_*and passed to a two-layer head that output the predicted response value.

#### Splits and training

Within each cell line, drugs were grouped by identical target sets, and whole groups were assigned to 70/15/15 training, validation and test partitions, repeated for 10 random folds. Models were trained with mean-squared error and AdamW for up to 100 epochs with early stopping on validation loss (patience 8). The configuration with the lowest validation mean-squared error over eight combinations of learning rate, weight decay and dropout was used to predict the test drugs (**Table** S2). Test pairs lacking a drug or gene embedding were assigned the training mean of that gene, and all embedding baselines were trained identically.

#### Evaluation

Following Systema [40], we subtracted the mean response of each gene over the training drugs of that cell line, *T̅_g_*, from both the measured profile *T̂_g_*and the predicted profile *T̂_g_*, and computed for each drug

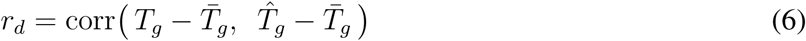

across genes before averaging over drugs. The control-reference variant omitted the subtraction. As a reference for measurement reproducibility, we computed the same metric between independent replicates of the same drug, using LINCS compounds profiled in separate campaigns and the 42 Tahoe drugs measured on replicate plates 6 and 14 (**Supplementary Information**).

#### Baselines

The gene-embedding baselines were scGPT-Pan-Cancer, Geneformer-V2 104M, Transcript-Former, scPRINT-2, Gene2Vec, GenePT and STRING (GNN), each reduced to 128 dimensions by PCA and passed through the EMBER-DRUG probe (**Table** S1). Structure baselines replaced the target set with a drug-level vector from CheMeleon [90], ChemBERTa [91] or a 2,048-dimensional Morgan fingerprint kept at full dimension, and used a learnable embedding for the readout gene. XPert [71], a transformer that predicts L1000 responses from molecular features and the control expression profile, was trained as a separate model using the code released by its authors, with the hyperparameters they used for their unseen-drug task, and was evaluated on the same folds. We excluded MiniMol [92] because its pretraining included expression prediction on LINCS L1000.

#### Disease-signature reversal in A549

Following TranSiGen [93], we built a lung adenocarcinoma signature from harmonized TCGA and GTEx expected counts [94, 95], comparing TCGA primary tumors with GTEx lung using DESeq2 [81]. Genes with *|* log_2_ FC*| >* 1 and adjusted *p <* 0.1 formed the up and down sets.

EMBER-DRUG was trained on A549 LINCS consensus profiles (10 *µ*M, 24 h) with gene embeddings contextualized on vehicle-treated A549 cells. Drugs were split 85/15 into training and validation by target-set group, which left 38 validation drugs and no test fold (**Table** S2). The selected model predicted 974 landmark responses for 1,277 DrugBank compounds with at least one target and at least half of their targets embedded, and the L1000 inference matrix [70] expanded each profile to 10,174 genes (**Supplementary Information**).

Each drug was scored against the disease gene sets with the connectivity score [96], where more negative scores indicate stronger predicted reversal. We retained drugs with at least one target detected in at least 5% of A549 control cells (417 drugs) and then removed drugs used for training, which left 303 ranked candidates.

#### Evaluation against PRISM drug viability

We evaluated the A549 disease-reversal scores using PRISM secondary-screen dose–response viability data for A549 (ACH-000681). Drug identifiers were matched by BRD core, and non-missing viability AUC values were aggregated by their median per drug. Hits were defined as drugs with median viability AUC strictly below the 20th percentile across all PRISM-measured A549 drugs, calculated before intersecting with the model predictions. The intersection contained 399 drugs, including 74 hits; drugs without viability measurements were excluded. ROC curves and AUROC were calculated using the negated predicted CMap connectivity score, so stronger predicted disease reversal corresponded to higher scores.

## Acknowledgments

The authors would like to thank Evan Trop, Yang Zhang, Jeff Winchell, Ellie Haber, Shike Wang, Ethan Wu, Aiden Zhang, Lorenzo Bini, and Caitlin Strassburg for helpful discussions and feedback on the manuscript. This work was supported, in part, by the National Institutes of Health grants UM1HG011593 (J.M.), R03OD039980 (J.M.), UH3CA268202 (J.M.), R01HG007352 (J.M.), R01HG012303 (J.M.), R21DA061481 (J.M.), and U24HG012070 (J.M.). J.M. was additionally supported by the Ray and Stephanie Lane Professorship, a Guggenheim Fellowship from the John Simon Guggenheim Memorial Foundation, and a Google Research Award. N.H. was supported by a National Science Foundation Graduate Research Fellowship DGE2140739 and DGE2631988. The funders had no role in study design, data collection and analysis, decision to publish or preparation of the manuscript.

## Code availability

The pretraining, inference, and evaluation code is available on https://github.com/ma-compbiolab/EMBER/.

## Data availability

Processed pretraining and task-specific datasets are available on Hugging Face at NickDST/EMBER-data. Source datasets are listed below.

- Pretraining corpus: CellxGene Census (version 2025-11-08) [30].
- Protein–protein interaction network: STRING v12 [7].
- K562 and RPE1 Perturb-seq data: Genome-wide Perturb-seq portal [97].
- HepG2 and Jurkat Perturb-seq data: Gene Expression Omnibus (GEO), accession GSE264667 [37].
- HAP1 genetic-interaction map: Boone laboratory repository [47].
- Yeast digenic interaction data: Boone laboratory repository [44].
- Yeast trigenic interaction data: Boone laboratory repository [55].
- Synthetic-lethal pairs: SynLethDB [48].
- DANGO imputed yeast trigenic interactions: Zenodo, record 18843411.
- HAP1 control cells: GEO, accession GSE293013 [83].
- DepMap 24Q4 Public copy-number and CRISPR gene-effect data: DepMap portal [62].
- PRISM secondary-screen viability data: Cancer Dependency Map Drug Repurposing portal [98].
- Torin1 CRISPR screen results: Supplementary Table 11 of ref. [54] and Supplementary Data of ref. [52].
- LINCS L1000 data: CLUE data dashboard [70].
- Tahoe-100M control cells and pseudobulk differential-expression data: tahoebio/Tahoe-100M on Hugging Face [86].
- Single-cell control profiles: GEO accessions GSE139944 (A549 and MCF7) [87], GSE164614 (A375) [88] and GSE167262 (HT29) [89].
- Drug targets and mechanism-of-action annotations: DrugBank [69].
- Harmonized TCGA and GTEx expression data: UCSC Xena Toil hub [94, 95, 99].

## Author Contributions

Conceptualization, N.H., J.M.; Methodology, N.H., J.M; Software, N.H.; Investigation, N.H., E.P.X., J.M; Writing, N.H., J.M; Funding Acquisition, J.M.

## References

[1] Kwon, J. J., Pan, J., Gonzalez, G., Hahn, W. C. & Zitnik, M. On knowing a gene: A distributional hypothesis of gene function. Cell Systems 15, 488–496 (2024).

[2] Ittisoponpisan, S., Alhuzimi, E., Sternberg, M. J. E. & David, A. Landscape of Pleiotropic Proteins Causing Human Disease: Structural and System Biology Insights. Human Mutation 38, 289–296 (2017).

3. The Tabula Sapiens Consortium. The Tabula Sapiens: A multiple-organ, single-cell transcriptomic atlas of humans. Science 376, eabl4896 (2022).

[4] Greene, C. S. et al. Understanding multicellular function and disease with human tissue-specific networks. Nature Genetics 47, 569–576 (2015).

[5] Shen, J. P. et al. Combinatorial CRISPR–Cas9 screens for de novo mapping of genetic interactions. Nature Methods 14, 573–576 (2017).

[6] Norman, T. M. et al. Exploring genetic interaction manifolds constructed from rich single-cell phenotypes. Science (New York, N.Y.) 365, 786–793 (2019).

[7] Szklarczyk, D. et al. The STRING database in 2025: protein networks with directionality of regulation. Nucleic Acids Research 53, D730–D737 (2025).

[8] Li, M. M. et al. Contextual AI models for single-cell protein biology. Nature Methods 21, 1546–1557 (2024).

[9] Chen, Y. & Zou, J. Simple and effective embedding model for single-cell biology built from ChatGPT. Nature Biomedical Engineering 9, 483–493 (2025).

[10] Hedley, J. G., Torr, P. H. S. & Märtens, K. GenePT Revisited: Do Better Text Embeddings Make Better Gene Embeddings? bioRxiv (2026).

[11] Zhong, J., Li, L., Dannenfelser, R. & Yao, V. Benchmarking gene embeddings from sequence, expression, network, and text models for functional prediction tasks. bioRxiv (2025).

12. Cole, E., et al. Foundation Models Improve Perturbation Response Prediction. bioRxiv (2026).

[13] Theodoris, C. V. et al. Transfer learning enables predictions in network biology. Nature 618, 616–624 (2023).

[14] Cui, H. et al. scGPT: toward building a foundation model for single-cell multi-omics using generative AI. Nature Methods 21, 1470–1480 (2024).

[15] Hao, M. et al. Large-scale foundation model on single-cell transcriptomics. Nature Methods 21, 1481–1491 (2024).

[16] Kalfon, J., Peyré, G. & Cantini, L. scPRINT-2: Towards the next-generation of cell foundation models and benchmarks. bioRxiv (2026).

17. Ho, N., et al. Scaling Dense Representations for Single Cell with Transcriptome-Scale Context. bioRxiv (2024).

[18] Pearce, J. D. et al. TranscriptFormer: A generative cell atlas across 1.5 billion years of evolution. Science 393, aec8514 (2026).

[19] Yang, F. et al. scBERT as a large-scale pretrained deep language model for cell type annotation of single-cell RNA-seq data. Nature Machine Intelligence 4, 852–866 (2022).

[20] Liu, T., Li, K., Wang, Y., Li, H. & Zhao, H. Evaluating the Utilities of Foundation Models in Single-Cell Data Analysis. Advanced Science 13, e14490 (2026).

[21] Bendidi, I., et al. Benchmarking Transcriptomics Foundation Models for Perturbation Analysis : one PCA still rules them all. *arXiv* (2024).

[22] Ahlmann-Eltze, C., Huber, W. & Anders, S. Deep-learning-based gene perturbation effect prediction does not yet outperform simple linear baselines. Nature Methods 22, 1657–1661 (2025).

[23] Boiarsky, R. et al. Deeper evaluation of a single-cell foundation model. Nature Machine Intelligence 6, 1443–1446 (2024).

[24] Kedzierska, K. Z., Crawford, L., Amini, A. P. & Lu, A. X. Zero-shot evaluation reveals limitations of single-cell foundation models. Genome Biology 26, 101 (2025).

[25] Chen, H. et al. Scaling and quantization of large-scale foundation model enables resource-efficient predictions in network biology. Nature Computational Science 6, 450–463 (2026).

[26] Adduri, A. K. et al. Predicting cellular responses to perturbation across diverse contexts with State. Cell 189, 5914–5931.e20 (2026).

[27] Dong, M., et al. Stack: In-Context Learning of Single-Cell Biology (2026). ISSN: 2692-8205 Pages: 2026.01.09.698608 Section: New Results.

[28] Roohani, Y., Huang, K. & Leskovec, J. Predicting transcriptional outcomes of novel multigene perturbations with GEARS. Nature Biotechnology 42, 927–935 (2024).

29. Verma, R., et al. PIE: Generalizing perturbation effects across unseen perturbations, contexts and datasets (2026). ISSN: 2692-8205 Pages: 2026.10.02.756297 Section: New Results.

30. CZI Cell Science Program et al. CZ CELLxGENE Discover: a single-cell data platform for scalable exploration, analysis and modeling of aggregated data. Nucleic Acids Research 53, D886–D900 (2025).

[31] DenAdel, A. et al. Evaluating the role of pretraining dataset size and diversity on single-cell foundation model performance. Nature Methods 1–11 (2026).

32. Dibaeinia, P., et al. Virtual Cells Need Context, Not Just Scale. bioRxiv (2026).

[33] Cultrera di Montesano, S., et al. Improving atlas-scale single-cell annotation models with hierarchical cross-entropy loss. Nature Computational Science 6, 243–249 (2026).

[34] Luck, K. et al. A reference map of the human binary protein interactome. Nature 580, 402–408 (2020).

35. Oord, A. v. d., Li, Y. & Vinyals, O. Representation Learning with Contrastive Predictive Coding. arXiv (2019).

[36] McInnes, L., Healy, J. & Melville, J. UMAP: Uniform Manifold Approximation and Projection for Dimension Reduction. *arXiv* (2020).

[37] Nadig, A. et al. Transcriptome-wide analysis of differential expression in perturbation atlases. Nature Genetics 57, 1228–1237 (2025).

[38] Bunne, C. et al. Learning single-cell perturbation responses using neural optimal transport. Nature Methods 20, 1759–1768 (2023).

[39] Song, L., Segal, E. & Xing, E. Toward AI-Driven Digital Organism: Multiscale Foundation Models for Predicting, Simulating and Programming Biology at All Levels. *arXiv* (2024).

[40] Viñas Torné, R., et al. Systema: a framework for evaluating genetic perturbation response prediction beyond systematic variation. Nature Biotechnology 44, 1050–1059 (2026).

[41] Mejia, G. M. et al. Diversity by Design: Addressing Mode Collapse Improves scRNA-seq Perturbation Modeling on Well-Calibrated Metrics. arXiv (2025).

[42] Lin, Z. et al. Evolutionary-scale prediction of atomic-level protein structure with a language model. Science 379, 1123–1130 (2023).

[43] Zinchenko, V. et al. Benchmarking Chemical, Genetic, and Cell Line Encodings for Cancer Perturbation Response Prediction. bioRxiv (2025).

[44] Costanzo, M. et al. A global genetic interaction network maps a wiring diagram of cellular function. Science 353, aaf1420 (2016).

[45] Srivas, R. et al. A Network of Conserved Synthetic Lethal Interactions for Exploration of Precision Cancer Therapy. Molecular Cell 63, 514–525 (2016).

[46] Nijman, S. M. Synthetic lethality: General principles, utility and detection using genetic screens in human cells. Febs Letters 585, 1–6 (2011).

[47] Billmann, M. et al. Global genetic interaction network of a human cell maps conserved principles and informs functional interpretation of gene co-essentiality profiles. Cell 189, 3736–3759.e58 (2026).

[48] Wang, J. et al. SynLethDB 2.0: a web-based knowledge graph database on synthetic lethality for novel anticancer drug discovery. Database: The Journal of Biological Databases and Curation 2022, baac030 (2022).

[49] Luo, J. et al. A genome-wide RNAi screen identifies multiple synthetic lethal interactions with the Ras oncogene. Cell 137, 835–848 (2009).

[50] Zamanighomi, M. et al. GEMINI: a variational Bayesian approach to identify genetic interactions from combinatorial CRISPR screens. Genome Biology 20, 137 (2019).

[51] Horlbeck, M. A. et al. Mapping the Genetic Landscape of Human Cells. Cell 174, 953–967.e22 (2018).

[52] Hermann, J. et al. mTORC1 cooperates with tRNA wobble modification to sustain the protein synthesis machinery. Nature Communications 16, 4201 (2025).

[53] Huang, H. et al. mTOR-mediated phosphorylation of VAMP8 and SCFD1 regulates autophagosome maturation. Nature Communications 12, 6622 (2021).

[54] Gonatopoulos-Pournatzis, T. et al. Genetic interaction mapping and exon-resolution functional genomics with a hybrid Cas9–Cas12a platform. Nature Biotechnology 38, 638–648 (2020).

[55] Kuzmin, E. et al. Systematic analysis of complex genetic interactions. Science 360, eaao1729 (2018).

[56] Zhang, R., Bafna, M., Ma, J. & Ma, J. Dango: Predicting higher-order genetic interactions. Cell Systems 17 (2026).

[57] Ruepp, A. et al. CORUM: the comprehensive resource of mammalian protein complexes—2009. Nucleic Acids Research 38, D497–D501 (2010).

[58] Milacic, M. et al. The Reactome Pathway Knowledgebase 2024. Nucleic Acids Research 52, D672–D678 (2024).

[59] Kanehisa, M. & Goto, S. KEGG: Kyoto Encyclopedia of Genes and Genomes. Nucleic Acids Research 28, 27–30 (2000).

[60] Jahn, R., Cafiso, D. C. & Tamm, L. K. Mechanisms of SNARE proteins in membrane fusion. Nature Reviews Molecular Cell Biology 25, 101–118 (2024).

[61] Jurado, A. R., Tan, D., Jiao, X., Kiledjian, M. & Tong, L. Structure and Function of Pre-mRNA 5*^′^* -End Capping Quality Control and 3*^′^* -End Processing. Biochemistry 53, 1882–1898 (2014).

[62] Tsherniak, A. et al. Defining a Cancer Dependency Map. Cell 170, 564–576.e16 (2017).

[63] Nicklin, P. et al. Bidirectional Transport of Amino Acids Regulates mTOR and Autophagy. Cell 136, 521–534 (2009).

[64] Li, T. et al. TMED10 mediates the trafficking of insulin-like growth factor 2 along the secretory pathway for myoblast differentiation. Proceedings of the National Academy of Sciences 120, e2215285120 (2023).

[65] Khoriaty, R. et al. Functions of the COPII gene paralogs SEC23A and SEC23B are interchangeable in vivo. Proceedings of the National Academy of Sciences 115, E7748–E7757 (2018).

[66] Iorio, F. et al. A Landscape of Pharmacogenomic Interactions in Cancer. Cell 166, 740–754 (2016).

[67] Barretina, J. et al. The Cancer Cell Line Encyclopedia enables predictive modelling of anticancer drug sensitivity. Nature 483, 603–607 (2012).

[68] Garnett, M. J. et al. Systematic identification of genomic markers of drug sensitivity in cancer cells. Nature 483, 570–575 (2012).

[69] Knox, C. et al. DrugBank 6.0: the DrugBank Knowledgebase for 2024. Nucleic Acids Research 52, D1265–D1275 (2024).

[70] Subramanian, A. et al. A Next Generation Connectivity Map: L1000 Platform and the First 1,000,000 Profiles. Cell 171, 1437–1452.e17 (2017).

[71] Guo, Y. et al. Modelling drug-induced cellular perturbation responses with a biologically informed dual-branch transformer. Nature Machine Intelligence 8, 96–112 (2026).

[72] Wolf, J. et al. Capmatinib in MET Exon 14–Mutated or MET-Amplified Non–Small-Cell Lung Cancer. New England Journal of Medicine 383, 944–957 (2020).

[73] Paik, P. K. et al. Tepotinib in Non–Small-Cell Lung Cancer with MET Exon 14 Skipping Mutations. New England Journal of Medicine 383, 931–943 (2020).

[74] Camidge, D. R. et al. Brigatinib versus Crizotinib in ALK-Positive Non–Small-Cell Lung Cancer. New England Journal of Medicine 379, 2027–2039 (2018).

[75] Johnson, F. M. et al. Phase II Study of Dasatinib in Patients With Advanced Non–Small-Cell Lung Cancer. Journal of Clinical Oncology 28, 4609–4615 (2010).

[76] Ou, S.-H. I., Park, C. J., Arter, Z. L. & Nagasaka, M. Successful and On-going Long-Term Disease Control (>24 Months) with Gilteritinib in an ALK+ NSCLC Patient with Brain Metastasis Who Has Progressed on Multiple ALK TKIs. A Case Report and Review of Literature on Gilteritnib. Lung Cancer 16, 139–145 (2025).

[77] Hellmann, M. D. et al. Phase Ib study of atezolizumab combined with cobimetinib in patients with solid tumors. Annals of Oncology 30, 1134–1142 (2019).

[78] Cui, J. et al. HDAC inhibitor ITF2357 reduces resistance of mutant-KRAS non-small cell lung cancer to pemetrexed through a HDAC2/miR-130a-3p-dependent mechanism. Journal of Translational Medicine 21, 125 (2023).

[79] Fischer, F. et al. scTab: Scaling cross-tissue single-cell annotation models. Nature Communications 15, 6611 (2024).

80. Zaheer, M., et al. Deep Sets. *arXiv* (2018).

[81] Love, M. I., Huber, W. & Anders, S. Moderated estimation of fold change and dispersion for RNA-seq data with DESeq2. Genome Biology 15, 550 (2014).

[82] Squair, J. W. et al. Confronting false discoveries in single-cell differential expression. Nature Communications 12, 5692 (2021).

[83] Kumari, B. et al. Single-cell exon deletion profiling reveals splicing events that shape gene expression and cell state dynamics. Nature Communications 17, 1218 (2026).

[84] Guo, J., Liu, H. & Zheng, J. SynLethDB: synthetic lethality database toward discovery of selective and sensitive anticancer drug targets. Nucleic Acids Research 44, D1011–1017 (2016).

[85] O’Brien, K. P., Remm, M. & Sonnhammer, E. L. L. Inparanoid: a comprehensive database of eukaryotic orthologs. Nucleic Acids Research 33, D476–480 (2005).

[86] Zhang, J. et al. Tahoe-100M: Mapping drug-induced molecular phenotypes at single-cell resolution. Cell 189, 5945–5961.e8 (2026).

[87] Srivatsan, S. R. et al. Massively multiplex chemical transcriptomics at single-cell resolution. Science 367, 45–51 (2020).

[88] Yang, C., Tian, C., Hoffman, T. E., Jacobsen, N. K. & Spencer, S. L. Melanoma subpopulations that rapidly escape MAPK pathway inhibition incur DNA damage and rely on stress signalling. Nature Communications 12, 1747 (2021).

[89] Miller, S. A. et al. LSD1 and Aberrant DNA Methylation Mediate Persistence of Enteroendocrine Progenitors That Support BRAF-Mutant Colorectal Cancer. Cancer Research 81, 3791–3805 (2021).

[90] Burns, J. W. et al. Deep Learning Foundation Models for Low-Data Regimes from Classical Molecular Descriptors. Journal of Chemical Information and Modeling 66, 10692–10701 (2026).

[91] Chithrananda, S., Grand, G. & Ramsundar, B. ChemBERTa: Large-Scale Self-Supervised Pretraining for Molecular Property Prediction. *arXiv* (2020).

[92] Kläser, K., et al. MiniMol: A Parameter-Efficient Foundation Model for Molecular Learning. *arXiv* (2024).

[93] Tong, X. et al. Deep representation learning of chemical-induced transcriptional profile for phenotype-based drug discovery. Nature Communications 15, 5378 (2024).

[94] Goldman, M. J. et al. Visualizing and interpreting cancer genomics data via the Xena platform. Nature Biotechnology 38, 675–678 (2020).

[95] Weinstein, J. N. et al. The Cancer Genome Atlas Pan-Cancer Analysis Project. Nature genetics 45, 1113–1120 (2013).

[96] Lamb, J. et al. The Connectivity Map: Using Gene-Expression Signatures to Connect Small Molecules, Genes, and Disease. Science 313, 1929–1935 (2006).

[97] Replogle, J. M. et al. Mapping information-rich genotype-phenotype landscapes with genome-scale Perturb-seq. Cell 185, 2559–2575.e28 (2022).

[98] Corsello, S. M. et al. Discovering the anti-cancer potential of non-oncology drugs by systematic viability profiling. Nature cancer 1, 235–248 (2020).

[99] Vivian, J. et al. Toil enables reproducible, open source, big biomedical data analyses. Nature Biotechnology 35, 314–316 (2017).

