## Supplemental Information for "EMBER: Contextualized gene representations enable compositional prediction across cellular perturbations"

#### Supplementary Notes

##### Pretraining corpus curation

Cells were drawn from the CellxGene Census (version 2025-11-08) and filtered in the following order, similar to scTab [1].

1. We kept *Homo sapiens* cells flagged as primary data, so that cells from redeposited datasets were counted once.
2. We kept 10x assays only (10x 5′ v1 and v2, 10x 3′ v1, v2 and v3, and the corresponding 10x transcription-profiling protocols).
3. We curated cell types against the Cell Ontology (version 2023-05-22). A cell type was removed if it was absent from the ontology, was not a descendant of *native cell*, contained fewer than 1,000 cells, was covered by 30 or fewer technical samples (one technical sample is one combination of dataset and donor), or had 7 or fewer ontology ancestors.
4. We balanced the corpus to approximately 250,000 cells with a quota of $250{,}000/n_{\text{cell-types}}$ cells per cell type. Cell types smaller than the quota were kept in full. Within each cell type, cells were drawn across batches, where a batch is one combination of dataset, donor, assay and general tissue.
5. We pulled raw UMI counts for all genes and restricted the gene panel to protein-coding genes.

We then added 2,000 control cells from each of K562, Jurkat, HepG2 and RPE1. Approximately 1% of cells were held out as a validation set.

##### Embedding space analyses

All analyses used frozen embeddings without task-specific training.

**GO recovery.** We measured GO recovery with ARI and NMI in HepG2, Jurkat, K562 and RPE1, evaluating each cell line separately. For each cell line, we used the top 25% most highly expressed genes shared across models and GO annotations propagated by the true-path rule. A clustering task was a GO parent with at least three children, each containing at least ten genes in every cell line. Genes assigned to exactly one child were clustered with $k$ equal to the number of children. We required at least fifty single-label genes per parent and no child larger than five times the mean size of the others, which yielded 13 Biological Process, 6 Molecular Function and 14 Cellular Component tasks (**Table** S3).

**Jurkat gene embedding map.** We removed unannotated genes, genes with only trivial annotations and the 40% of genes closest to the centroid of the Jurkat gene embeddings, which left 4,601 genes. These were clustered by Leiden community detection [2] at the resolution that maximized Newman modularity (0.7), which gave 22 clusters. Each cluster was tested for GO enrichment with a hypergeometric test against the plotted genes as background, at Benjamini-Hochberg FDR $< 0.05$ and a fold enrichment of at least two. Embeddings were visualized with UMAP [3] (25 neighbors, minimum distance 0.15).

**Lineage Clustering.** We embedded a gene once per cell, so that its tokenized context differed between cells. We drew 256 cells per lineage, spread evenly across the cell types of each lineage, over 11 lineages (2,816 cells). *COL1A1*, an established fibroblast marker, was used for visualization. For each gene in a panel of 18 lineage markers, 18 broadly expressed transcription factors and 18 housekeeping genes (**Table** S3), the per-cell embeddings were clustered by $k$-means with $k = 11$ and scored against the lineage labels with ARI and NMI.

**Perturbation responsiveness.** For each context $T$ (HepG2, Jurkat, K562 or RPE1), responsiveness was the fraction of assayed perturbations $\boldsymbol{\mathcal{P}}_T$ that induced differential expression of gene $g$, and embedding distance was the Euclidean ($L_2$) distance from the mean gene embedding $\bar{z}^T$,

$$\rho_T(g) = \frac{1}{|\boldsymbol{\mathcal{P}}_T|} \sum_{p \in \boldsymbol{\mathcal{P}}_T} \mathbb{1}\left[\mathrm{padj}_T(p, g) < 0.05\right], \qquad d_T(g) = \left\| z_g^T - \bar{z}^T \right\|_2. \tag{7}$$

Adjusted $p$-values came from the DESeq2 analysis of each cell line [4]. We regressed expression magnitude, expression variance and tokenization sampling frequency out of both quantities and computed the Spearman correlation between the residuals of $d_T$ and $\rho_T$ across shared genes, for matched and mismatched pairs of embedding context and Perturb-seq cell line. Embeddings came from the reconstruction-only model pretrained on 100 cell types described below.

**Cell-type diversity at fixed cell count.** We selected 100 ontology-curated cell types by census abundance, excluding cell-type families that overlap K562, RPE1, Jurkat or HepG2. We built four nested subsets of 1, 10, 50, and 100 cell types, each with approximately 176,000 cells distributed evenly across types. One model was trained per subset with masked reconstruction only, with architecture and optimization held fixed (AdamW, cosine schedule, batch size 64, 100 epochs).

#### Edge memorization test

To assess whether CSCL memorizes training STRING edges, we repeated pretraining under two edge-holdout schemes with the recipe and epoch budget unchanged. We evaluated held-out edges in held-out cells, testing generalization to both unseen edges and unseen cells.

**Random edge holdout.** We split undirected STRING pairs into training, validation, and test sets (80/10/10), keeping both orientations of each pair in the same split. Only training edges contributed to the contrastive loss.

**Pathway holdout.** We removed all STRING edges between genes within each held-out pathway, retaining the genes and their edges to genes outside that pathway. We selected pathways from Enrichr's Reactome 2022 and KEGG 2021 Human libraries [5], requiring 8–50 genes and at least 20 internal STRING edges. The resulting 65 pathways contained 4,412 unique held-out edges, leaving approximately 97% of STRING edges available for training. We reserved a random 2% of the remaining edges for validation and epoch selection.

**Scoring.** For each held-out cell, we ran the model unmasked, projected contextual gene embeddings through the trained graph head, and L2-normalized the outputs. We scored gene pairs by cosine similarity. Each held-out edge with both endpoints present generated two directed queries. For each query, we ranked the target against non-partners among the approximately 1,024 expressed genes in the cell, excluding the anchor and all other known STRING partners. We computed mean reciprocal rank (MRR) and Hits@1, Hits@10, and Hits@50 over directed queries within each cell, then averaged across cells with equal weight. Cells were eligible for a split only if their context contained at least three edges from that split; at most 512 cells were scored. For the trained-edge reference, positives were subsampled to at most 200 per cell. We report results from the epoch with the highest validation MRR.

#### Mismatched-context embeddings

For each target cell line, we ranked the CellxGene cell types of the pretraining corpus by the Pearson correlation between their pseudobulk expression and the expression of the target cell line. The mismatched context was the cell type with a correlation closest to 0.5 (**Table** S4). In the tasks that transfer across cell lines, each cell line received a distinct cell type.

#### Differential expression calls for GEARS and STATE

Both models predict expression profiles, whereas the benchmark scores differentially expressed gene calls. For DESeq2, which operates on replicate-aware pseudobulks, we inverse-transformed the predicted profiles using the library sizes of the input control cells and summed counts within each replicate. For the Wilcoxon test, we used the Scanpy implementation on predicted and control cells. Both procedures used FDR correction.

#### Prediction quality stratified by PPI prior knowledge

We compared predictions based on EMBER and STRING (GNN) embeddings on the unseen-perturbation generalization split to assess performance on unseen perturbations within the same cellular context. For each perturbed gene, we calculated AUPRC across samples in which that gene was perturbed. We then grouped genes by their number of interactions in STRING v12.5 with confidence scores $> 0.8$ and plotted the mean per-perturbation AUPRC for each group.

#### Macro-Pert AUROC and AUPRC

Perturbations with more retained readout genes contribute more observations to pooled metrics. Curation limits each perturbation to 5–100 differentially expressed genes ($M_{\min} = 5$, $M_{\max} = 100$), retaining the top 100 by $|\log_2 \mathrm{FC}|$ when necessary. To assess sensitivity to unequal representation, we computed AUROC and AUPRC separately for each perturbation and averaged each metric with equal weight across perturbations (**Table** S5). EMBER remained top-ranked on both metrics. Because estimates for perturbations with few readout genes can be unstable, we retain pooled metrics as the primary benchmarks and macro-per-perturbation metrics as complementary robustness checks.

#### MTOR–SCFD1 nomination and Torin1 screens

**Candidate nomination.** We scored every *MTOR* partner in the symmetrized imputed map by its residual score, excluded pairs already in SynLethDB, and ranked the rest from most negative. The ranking used only the imputed map and no screen data. Because we examined only *SCFD1* in the published screens below, we report the published nominal $P$ values. *SCFD1* ranked 14th among the 16,408 genes in the map (99.92nd percentile). Neither gene was a query gene in the HAP1 map. After the nomination, literature review identified a published mechanism [6]. Active mTORC1 phosphorylates VAMP8, which blocks SCFD1 recruitment and assembly of the STX17–SNAP29–VAMP8 SNARE complex that mediates autophagosome–lysosome fusion. mTOR inhibition removes this block, and SCFD1 knockdown under Torin1 prevents fusion. Because mTOR inhibition increases autophagy, we hypothesized that it also increases cellular dependence on SCFD1, which the screens below provide retrospective evidence for.

**Human HAP1 Torin1 CRISPR screen.** The CHyMErA Torin1 screen in HAP1 cells [7] split one transduced population into three technical arms per condition, grown with Torin1 or vehicle from day 3 to day 18. We used its dual-targeting analysis, in which both guides of each hybrid construct target *SCFD1*. The authors compared log2 fold changes relative to T0 between conditions with a two-sided Wilcoxon rank-sum test. Effect estimates and P values were taken directly from the original study's Supplementary Table 11, without reprocessing the raw counts; n = 15 represents five constructs across three technical arms per condition. Vehicle and Torin1 log2 fold changes were $-2.01$ and $-2.69$ at day 12 ($p = 0.300$) and $-2.21$ and $-3.69$ at day 18 ($p = 0.062$) (**Fig. 4g** and **Fig. S3g**).

**Comparing *SCFD1* to other baseline essential genes** To test whether baseline essentiality explained the added depletion, we took the 465 genes in the bottom 10% of the HAP1 day-18 screen by vehicle log2 fold change ($\leq -4.36$). All of them were more depleted than *SCFD1* in vehicle ($-2.21$), yet only one had a more negative Torin1-minus-vehicle shift than *SCFD1* ($-1.48$), placing *SCFD1* at the 99.8th percentile (**Fig. S3i**).

**Mouse EPP2 Torin1 CRISPR screen.** We used published results from a genome-wide CRISPR–Cas9 proliferation screen in EPP2 mouse pancreatic cancer cells grown for 14 population doublings with 300 nM Torin1 or DMSO [8]. The authors computed gene-level log2 fold changes with MAGeCK v0.5.9, fitting DMSO versus T0, Torin1 versus T0 and Torin1 versus DMSO as separate contrasts. **Fig. S3h** shows the two T0 contrasts for *Scfd1* ($-1.47$ and $-1.89$) with the $P$ value of the direct Torin1-versus-DMSO contrast ($P = 0.0046$). The screen had one endpoint population per condition, so n=6 counts *Scfd1* sgRNAs.

#### Synthetic-lethality pre-filter classifier

The pre-filter was an XGBoost classifier of digenic synthetic lethality. Its features were the normalized Resnik similarities of the two genes in each of the three GO branches and the dot-product similarities of their ESM2 and STRING (GNN) embeddings. DepMap co-essentiality was excluded from the features, avoiding direct use of DepMap co-essentiality data in the subsequent co-loss analysis. Performance was estimated by 5-fold stratified cross-validation. Within each fold, hyperparameters were chosen by a 3-fold grid search scored by AUPRC over maximum depth $\{3, 5\}$, number of trees $\{100, 200\}$, learning rate $\{0.05, 0.1\}$ and minimum child weight $\{1, 5\}$, with classes reweighted by the ratio of negatives to positives. The classifier reached a mean AUROC of 0.697 over the five cross-validation folds. The final model was trained on all data with the most frequent best setting across folds (seed 42).

##### DepMap co-loss eligibility and statistical analysis

Relative copy number below $0.70$ defined single-copy loss and below $0.25$ defined deep loss. Eligible frames $(A, B; C)$ required loss calls for $A$ and $B$, CRISPR gene-effect data for $C$, and all three genes on distinct chromosome arms to limit shared-locus confounding. Minimum group sizes were $n_{AB} \geq 5$, $n_{A\text{-only}} \geq 3$, $n_{B\text{-only}} \geq 3$, and $n_{\text{neither}} \geq 2$, with co-loss represented in at least two tissue lineages.

We centered gene effects within tissue lineages and residualized them against readout-gene copy number and loss burden (the fraction of genes lost per cell line). We then fitted

$$C^{\text{res}}_{\text{effect}} \sim A_{\text{loss}} + B_{\text{loss}} + A_{\text{loss}} : B_{\text{loss}}.$$

For group means $m_{ij}$ and sizes $n_{ij}$, the interaction estimate and standard error were

$$\beta_{AB} = m_{11} - m_{10} - m_{01} + m_{00}, \qquad \text{se}(\beta_{AB}) = \sqrt{\frac{\text{RSS}}{n-4} \sum_{i,j \in \{0,1\}} n_{ij}^{-1}}.$$

Given sparse co-loss groups, we used this two-stage procedure as an exploratory screen; its interaction coefficients describe contrasts in residualized outcomes and are not equivalent to jointly covariate-adjusted effects. We used two-sided $t$-tests with degrees of freedom reduced for residualization covariates and Benjamini–Hochberg correction across frames. For each of 128 permutations, we jointly shuffled loss-matrix rows relative to fixed outcomes and covariates, preserving the joint distribution of loss calls, and reran the pipeline. Given observed and permuted significant-frame counts $H_{\text{obs}}$ and $H_b$, we computed

$$p_{\text{emp}} = \frac{1 + \sum_{b=1}^{128} \mathbb{1}[H_b \geq H_{\text{obs}}]}{129}, \qquad \text{enrichment} = \frac{H_{\text{obs}}}{128^{-1} \sum_{b=1}^{128} H_b}.$$

##### Tahoe cell line selection

Within a cell line $c$, we subtracted the mean log fold change $\bar{x}_c$ from the profile of each drug. For a drug $d$ measured on plates $A$ and $B$, and for two different drugs $d_1 \neq d_2$, we computed

$$\rho_d^{\text{repro}} = \text{corr}\big(x_{d,A} - \bar{x}_c,\ x_{d,B} - \bar{x}_c\big), \qquad \rho_{d_1,d_2}^{\text{dist}} = \text{corr}\big(x_{d_1} - \bar{x}_c,\ x_{d_2} - \bar{x}_c\big), \tag{8}$$

over pairwise-complete genes, using the 42 drugs with two plate replicates in every cell line. Cell lines were ranked by high mean $\rho^{\text{repro}}$ and low mean $\rho^{\text{dist}}$, subject to spanning different tissues. Using these criteria, we selected the top candidates: A172 (glioblastoma), BT474 (breast), NCIH460 (lung) and SW480 (colon).

#### Replicate reference for the drug-response metric

The replicate reference is the perturbed-reference Pearson correlation between independent measurements of the same drug response. A LINCS level-5 signature already pools the wells of one run, so we treated compounds rerun in a separate campaign, identified by a distinct project code, plate family or bead batch, as independent replicates. We divided the campaign replicates of each compound into two groups, averaged within each group, and computed the perturbed-reference correlation between the two averages over pairwise-complete genes, requiring at least five shared genes. Correlations were averaged over drugs within each cell line and fold. For Tahoe, plates 6 and 14 are exact replicates, and we computed the same correlation between the two plates for the 42 drugs measured on both.

#### Disease signature and L1000 expansion

Harmonized TCGA and GTEx values were converted back to counts, genes with low counts were removed, and differential expression between lung adenocarcinoma primary tumors and GTEx lung was tested with DESeq2. Genes were mapped to symbols, duplicates were resolved by keeping the most significant record, and genes without an adjusted $p$-value were excluded from the signature. Of the 978 L1000 landmark genes, 974 mapped to embedded gene symbols. The four unmapped landmark genes were excluded from training and assigned a zero differential at prediction. Because the model predicts differential responses, each landmark profile was expanded as $\Delta_{\text{inferred}} = \Delta_{\text{landmark}} W$, where $W$ is the To obtain an inference weight matrix W matched to LINCS Level 5 compound signatures, we fit ridge regression models that predict inferred-gene z-scores from landmark-gene z-scores. All signatures from A549, MCF7, A375, and HT29 were excluded from fitting W. Concatenating the 978 landmark values with the 9,196 inferred genes gave 10,174 genes. The connectivity score of a drug was the enrichment of the up set minus the enrichment of the down set within its ranked predicted profile.

#### References


[1] Fischer, F. *et al.* scTab: Scaling cross-tissue single-cell annotation models. *Nature Communications* **15**, 6611 (2024).

[2] Traag, V., Waltman, L. & Eck, N. J. v. From Louvain to Leiden: guaranteeing well-connected communities. *Scientific Reports* **9**, 5233 (2019).

[3] McInnes, L., Healy, J. & Melville, J. UMAP: Uniform Manifold Approximation and Projection for Dimension Reduction. *arXiv* (2020).

[4] Nadig, A. *et al.* Transcriptome-wide analysis of differential expression in perturbation atlases. *Nature Genetics* **57**, 1228–1237 (2025).

[5] Kuleshov, M. V. *et al.* Enrichr: a comprehensive gene set enrichment analysis web server 2016 update. *Nucleic Acids Research* **44**, W90–97 (2016).

[6] Huang, H. *et al.* mTOR-mediated phosphorylation of VAMP8 and SCFD1 regulates autophagosome maturation. *Nature Communications* **12**, 6622 (2021).

[7] Gonatopoulos-Pournatzis, T. *et al.* Genetic interaction mapping and exon-resolution functional genomics with a hybrid Cas9–Cas12a platform. *Nature Biotechnology* **38**, 638–648 (2020).

[8] Hermann, J. *et al.* mTORC1 cooperates with tRNA wobble modification to sustain the protein synthesis machinery. *Nature Communications* **16**, 4201 (2025).

[9] Guo, J., Liu, H. & Zheng, J. SynLethDB: synthetic lethality database toward discovery of selective and sensitive anticancer drug targets. *Nucleic Acids Research* **44**, D1011–1017 (2016).

[10] Cole, E. *et al.* Foundation Models Improve Perturbation Response Prediction. *bioRxiv* (2026).

[11] Chen, Y. & Zou, J. Simple and effective embedding model for single-cell biology built from ChatGPT. *Nature Biomedical Engineering* **9**, 483–493 (2025).

[12] Neelakantan, A. *et al.* Text and Code Embeddings by Contrastive Pre-Training. *arXiv* (2022).

[13] Istrate, A.-M., Li, D. & Karaletsos, T. scGenePT: Is language all you need for modeling single-cell perturbations? *bioRxiv* (2024).

[14] Li, M. M. *et al.* Contextual AI models for single-cell protein biology. *Nature Methods* **21**, 1546–1557 (2024).

[15] Cui, H. *et al.* scGPT: toward building a foundation model for single-cell multi-omics using generative AI. *Nature Methods* **21**, 1470–1480 (2024).

[16] Theodoris, C. V. *et al.* Transfer learning enables predictions in network biology. *Nature* **618**, 616–624 (2023).

[17] Chen, H. *et al.* Scaling and quantization of large-scale foundation model enables resource-efficient predictions in network biology. *Nature Computational Science* **6**, 450–463 (2026).

[18] Pearce, J. D. *et al.* TranscriptFormer: A generative cell atlas across 1.5 billion years of evolution. *Science* **393**, aec8514 (2026).

[19] Kalfon, J., Peyré, G. & Cantini, L. scPRINT-2: Towards the next-generation of cell foundation models and benchmarks. *bioRxiv* (2026).

[20] Du, J. *et al.* Gene2vec: distributed representation of genes based on co-expression. *BMC Genomics* **20**, 82 (2019).

[21] Su, J. *et al.* A trimodal protein language model enables advanced protein searches. *Nature Biotechnology* **44**, 1310–1316 (2026).

[22] Lin, Z. *et al.* Evolutionary-scale prediction of atomic-level protein structure with a language model.

*Science* **379**, 1123–1130 (2023).

[23] Nguyen, E. *et al.* HyenaDNA: Long-Range Genomic Sequence Modeling at Single Nucleotide Resolution. *arXiv* (2023).

[24] Roohani, Y., Huang, K. & Leskovec, J. Predicting transcriptional outcomes of novel multigene perturbations with GEARS. *Nature Biotechnology* **42**, 927–935 (2024).

[25] Adduri, A. K. *et al.* Predicting cellular responses to perturbation across diverse contexts with State. *Cell* **189**, 5914–5931.e20 (2026).

### Supplementary Figures

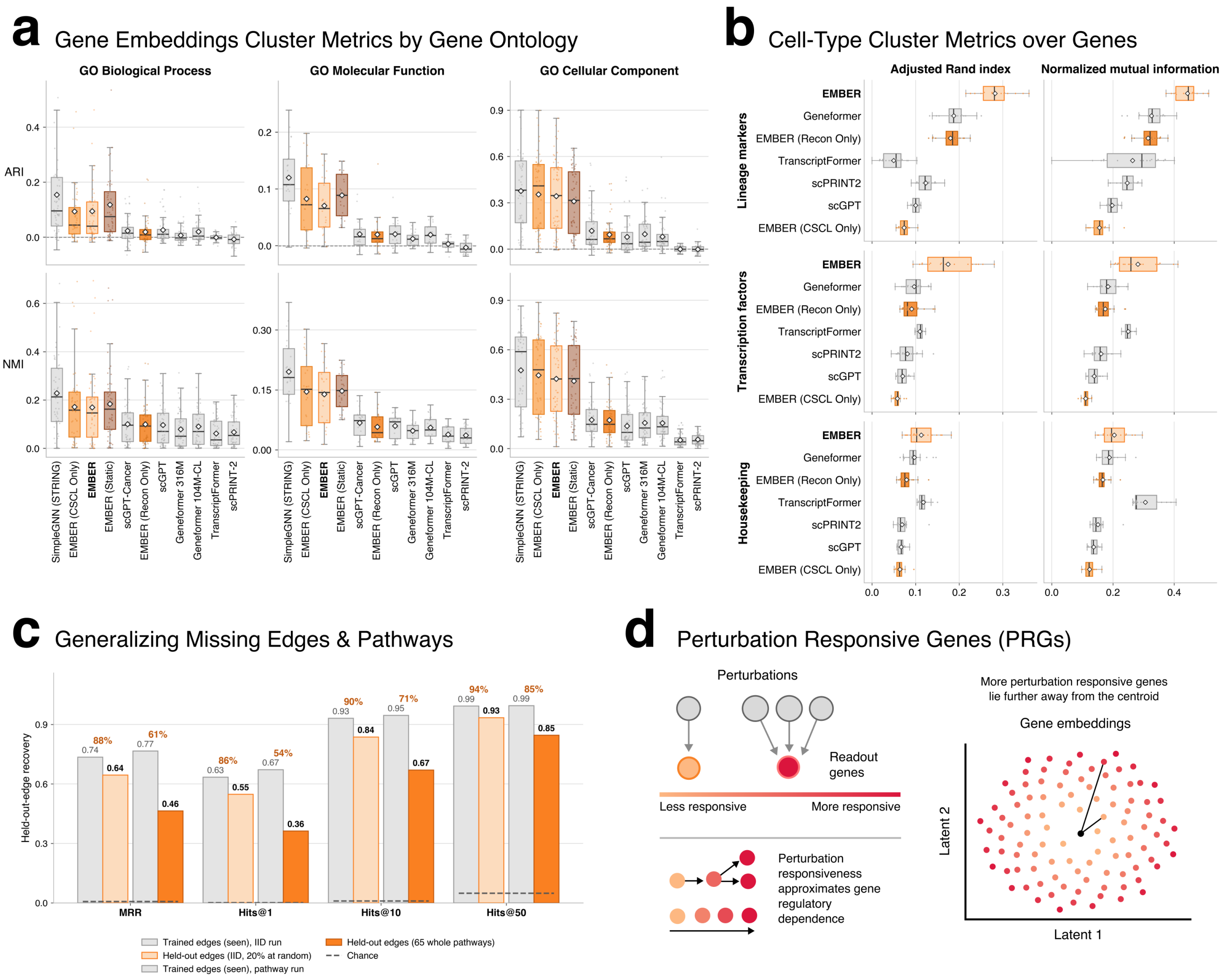


**Figure S1: Extended pretraining and embedding-organization analyses. a.** Per-category GO clustering metrics. Adjusted Rand index (top row) and normalized mutual information (bottom row) for EMBER and all compared embeddings, stratified by GO namespace into Biological Process, Molecular Function, and Cellular Component (**Methods**). Each point is one balanced GO parent term scored in one of the four Perturb-seq cell lines, giving 13, 6, and 14 parent terms per cell line for the three namespaces. Boxes show the median and interquartile range, and diamonds mark the mean. EMBER trails only the context-agnostic STRING (GNN) embedding and separates clearly from the single-cell foundation model baselines, whereas removing the CSCL objective (EMBER (Recon-Only)) reduces performance to the level of those baselines. Significance was assessed with a two-sided paired $t$-test across the $n = 4$ cell lines, where ** $p < 0.01$ and *** $p < 0.001$. **b.** Cell-type clustering metrics computed over genes, showing ARI and normalized mutual information for lineage markers, transcription factors, and housekeeping genes. EMBER separates cells by type most strongly for lineage markers and least for housekeeping genes, and outperforms scGPT and its own graph-only and no-graph ablations. Stars mark the significance of EMBER against scGPT, with n.s. denoting a non-significant difference. **c.** Generalizing to missing edges and pathways. Held-out-edge recovery measured by MRR and Hits@1, Hits@10, and Hits@50 for EMBER trained with 20% of edges held out at random and with edges connecting genes within 65 pathways withheld, each compared with the corresponding trained-edge performance and with chance. EMBER recovers a substantial proportion of held-out edges even when within-pathway edges are withheld from training (**Supplementary Information**). **d.** Schematic of perturbation responsiveness.

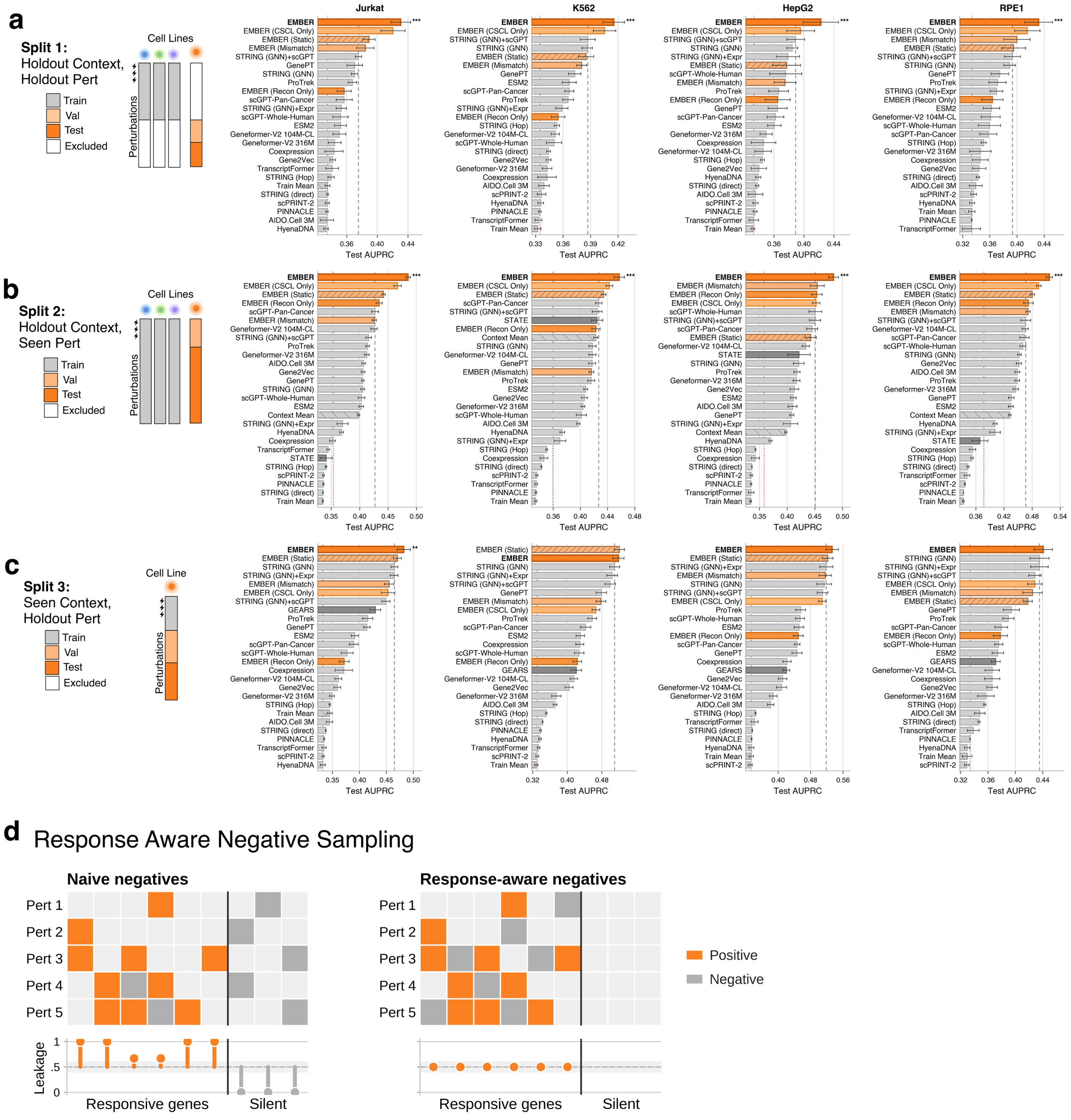


**Figure S2: Extended perturbation-prediction benchmark. a,** Unseen cell line with unseen perturbations. Test AUPRC for EMBER-PERT, its ablations and all baselines in each of the four essential-gene Perturb-seq cell lines, extending **Fig.** 3a. The schematic shows the assignment of perturbations to training, validation, test, and excluded sets. Bars show the mean and error bars the s.d. across $n = 10$ random partitions. The dashed line marks the best external baseline and the red dotted line marks chance. Stars compare EMBER with the best external baseline by a two-sided paired $t$-test, where ** $p < 0.01$ and *** $p < 0.001$. **b,** Unseen cell line with seen perturbations, extending **Fig.** 3b. Format and significance are the same as in panel **a**. **c,** Unseen perturbations within a single cell line, extending **Fig.** 3c. Format and significance are the same as in panel **a**. **d,** Negative sampling. With naive negatives, the label of a pair can be predicted from the identity of the readout gene alone (leakage, bottom). Response-aware negatives are drawn for the same readout genes under other perturbations, which reduces this leakage (**Methods**).

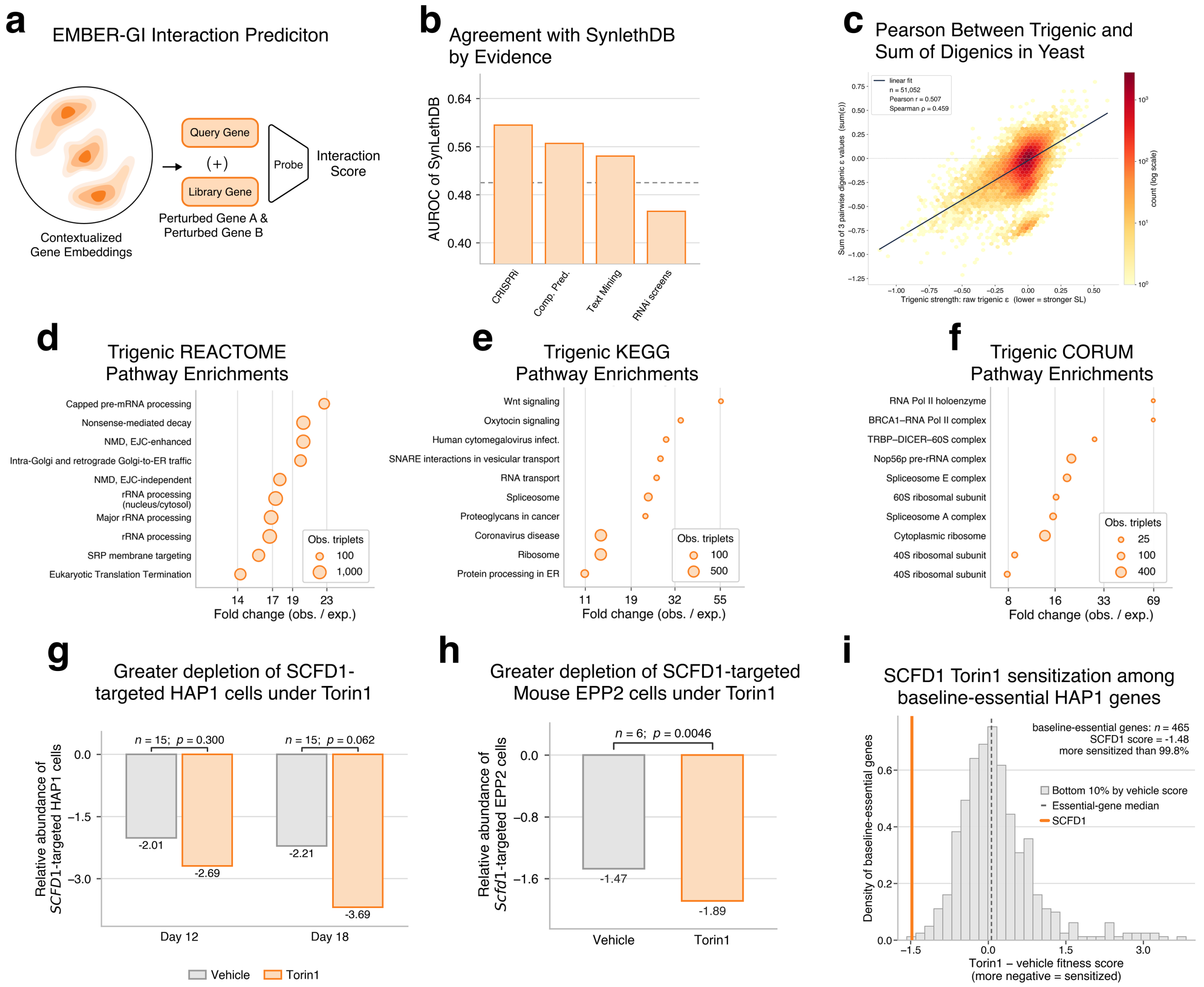


**Figure S3: Extended genetic-interaction results.** **a.** Schematic of EMBER-GI. Contextualized embeddings of a query gene and a library gene are concatenated and passed through a probe that predicts their interaction score. **b.** Agreement between the imputed genome-wide map and SynLethDB [9], stratified by evidence type and reported as AUROC. Concordance is highest for CRISPRi and lowest RNAi screens. **c.** Relationship between trigenic interaction strength and the sum of the three component pairwise digenic scores across 51,052 yeast triplets, colored by count on a log scale, where lower trigenic $\varepsilon$ indicates stronger synthetic lethality. The dark line is a linear fit, and the two measures show a Pearson correlation of $0.507$ and a Spearman correlation of $0.459$, motivating ranking candidate human triplets by the sum of their three predicted pairwise EMBER-GI scores (**Methods**). **d.** Reactome pathway enrichment of the prioritized trigenic triplets, ranked by fold enrichment of observed over expected, with point size scaled by the number of observed triplets. **e.** KEGG pathway enrichment of the prioritized trigenic triplets, in the same format as panel D. **f.** CORUM complex enrichment of the prioritized trigenic triplets, in the same format as panel D. **g.** Greater depletion of *SCFD1*-targeted HAP1 cells under Torin1 at days 12 and 18 [7]. **h.** Greater depletion of *Scfd1*-targeted mouse EPP2 cells under Torin1 [8]. **i.** Torin1 sensitization of *SCFD1* relative to baseline-essential HAP1 genes (**Supplementary Information**).

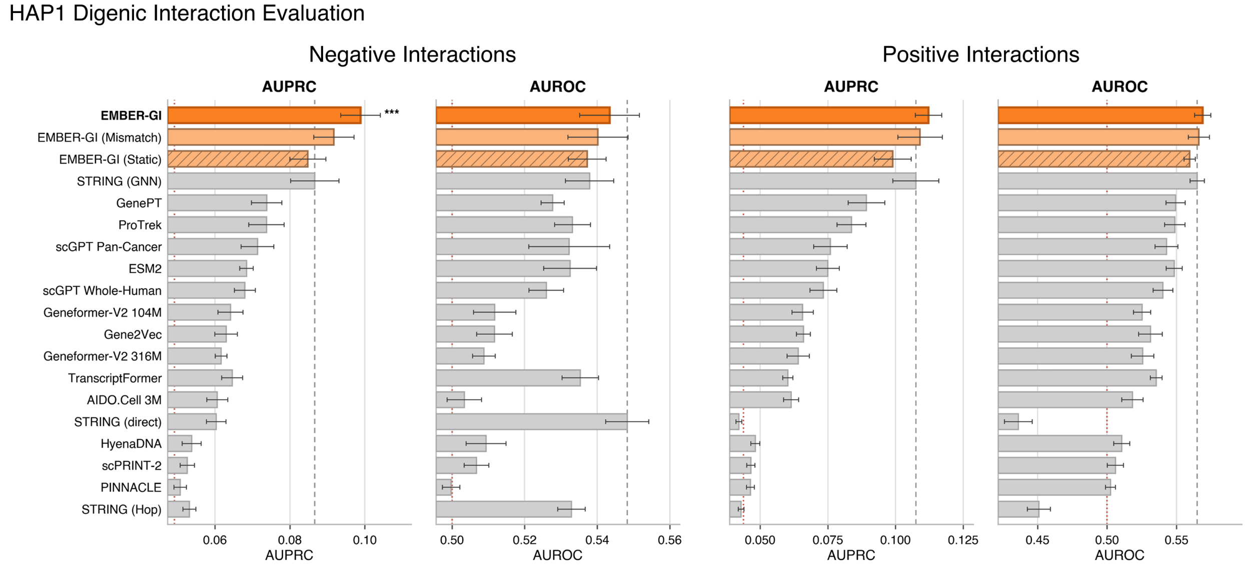


**Figure S4: Full HAP1 digenic interaction evaluation.** AUPRC and AUROC for negative-interaction and positive-interaction ranking under the unseen-library split, across EMBER-GI, its mismatched-context and static ablations, and all baselines, extending **Fig. 4**a. Bars show the mean and error bars the s.d. across $n = 10$ random splits. The dashed line marks the best external baseline and the red dotted line marks chance. Stars mark the significance of EMBER-GI against the best external baseline by a two-sided paired $t$-test, where * $p < 0.05$ and *** $p < 0.001$ (**Methods**).

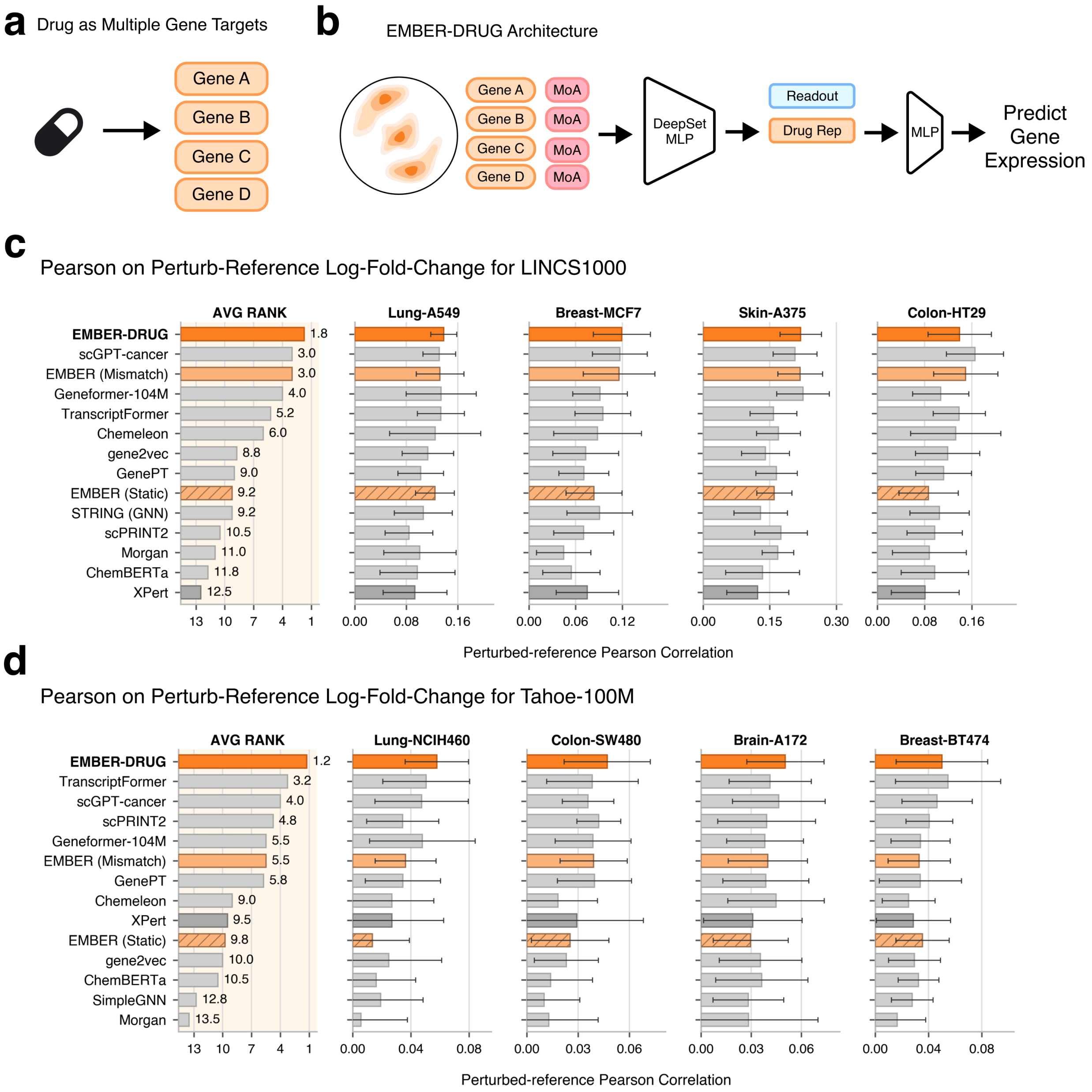


**Figure S5: Additional drug-response results. a.** A drug is represented as the set of its known target genes, each embedded in cellular context by EMBER. **b.** EMBER-DRUG architecture. Contextualized embeddings of a drug's target genes, together with their mechanism-of-action tags, are pooled by a DeepSets MLP into a drug representation, which a second MLP uses with a readout gene embedding to predict its expression change. **c.** Perturbed-reference Pearson correlation on LINCS L1000 for EMBER-DRUG and all baselines, shown per cell line and as the mean over the four cell lines. **d.** The same perturbed-reference evaluation on Tahoe-100M, in the same format as panel c.

#### Supplementary Tables

**Table S1: Perturbation-prediction baselines.** Baselines are grouped by the kind of signal they encode, with a short description, citation, and code link for each. Each embedding baseline (the Prior Knowledge, Cell, and Sequence groups, together with the Random noise floor under Trivial) is reduced to a per-gene embedding and scored with the same downstream MLP, so only the representation differs. Contextual baselines provide one vector per gene per cell line. The perturbation models predict perturbation responses directly and are described in the Perturbation model baselines section of the Methods.

| Baseline | Group | Representation |
|---|---|---|
| STRING (GNN) [10] | Prior Knowledge | Graph-convolutional-network embedding of the STRING PPI network, from Cole et al. link |
| STRING Direct | Prior Knowledge | Binary indicator of a direct STRING edge between the two genes, with no learned vector |
| STRING MultiHop | Prior Knowledge | Shortest-path hop distance between the two genes on the STRING graph, with no learned vector |
| STRING (GNN)+Expr | Prior Knowledge | STRING (GNN) embedding concatenated with the PCA of expression |
| STRING (GNN)+scGPT | Prior Knowledge | STRING (GNN) embedding concatenated with contextualized scGPT-Pan-Cancer embeddings |
| GenePT [11] | Prior Knowledge | Text-derived gene-description embedding from an OpenAI text-embedding model [12], taken from the scGenePT release [13] link |
| PINNACLE [14] | Prior Knowledge | Contextual cell-type-specific protein embedding from a hierarchical PPI network, with the four cell lines mapped to their best-matched ontology cell types link |
| scGPT, scGPT-Cancer [15] | Cell | Contextual gene embedding over control cells, whole-human and Pan-Cancer variants link |
| Geneformer-104M, Geneformer-316M [16, 17] | Cell | Rank-value transcriptome transformer embedding (V2), the 104M cancer continual-learning model and the 316M model link |
| TranscriptFormer [18] | Cell | Contextual gene embedding mean-pooled over control cells, the TF-Sapiens human variant link |
| scPRINT2 [19] | Cell | Contextual gene embedding mean-pooled over control cells |
| AIDO.Cell 3M | Cell | Pretrained gene-expression model, 3M-parameter variant |
| Coexpression-PCA | Cell | Cell-line-specific co-expression PCA embedding |
| Gene2Vec [20] | Cell | Static co-expression embedding computed across 984 GEO datasets |
| ProTrek [21] | Sequence | Tri-modal protein language model over structure, text, and sequence, averaged over each gene's isoforms |
| ESM2 [22] | Sequence | Protein language model embedding, averaged over amino acids and then isoforms |
| HyenaDNA [23] | Sequence | DNA language model embedding with 1-Mbp context, the last-token embedding of each gene |

*continued on next page*

*Table S1 (continued)*

| Baseline | Group | Representation |
|---|---|---|
| Train Mean | Trivial | Training positive frequency of the readout gene, with no perturbation information |
| Context Mean | Trivial | Mean differential-expression label of the same pair in the training cell lines, defined only for the unseen-cell-line regime |
| Random | Trivial | Random vectors, used as a noise floor |
| GEARS [24] | Perturbation model | Perturbation-response graph neural network that predicts effects directly |
| STATE [25] | Perturbation model | Perturbation-response model that predicts effects directly |

**Table S2: Model architectures and training hyperparameters.** Braces list the values searched, and the selected configuration was the best on the validation criterion of that column. Probes were trained on frozen EMBER embeddings with early stopping (tolerance $1 \times 10^{-4}$) and a fixed random seed of 42. All EMBER-DRUG models used an 8-dimensional mechanism-of-action embedding and at most 25 targets per drug. Pretraining used a linear warmup over 1,500 steps and cosine decay to $1 \times 10^{-6}$.

| | **Pretraining** | **EMBER-PERT** | **EMBER-GI** benchmark | **EMBER-GI** imputation | **EMBER-DRUG** LINCS L1000 | **EMBER-DRUG** Tahoe-100M | **EMBER-DRUG** A549 screen |
|---|---|---|---|---|---|---|---|
| Model | Transformer encoder | MLP, concatenated pair | MLP, concatenated pair | MLP, concatenated pair | Target-set DeepSet | Target-set DeepSet | Target-set DeepSet |
| Layers / width | 6 / 128 (FFN 512) | 2 / 128 | 4 / 256 | 4 / 256 | 2 / 64 | 2 / 64 | 2 / 64 |
| Loss | $\mathcal{L}_{\text{rec}} + 3.0\,\mathcal{L}_{\text{CSCL}}$ | Binary cross-entropy | MSE | MSE | MSE | MSE | MSE |
| Optimizer | AdamW | AdamW | Adam | Adam | AdamW | AdamW | AdamW |
| Max epochs / patience | 100 / 25 (epoch 24 used) | 200 / 10 | 200 / 10 | 200 / 10 | 100 / 8 | 100 / 8 | 100 / 8 |
| Learning rate | $10^{-3}$ | $\{10^{-2}, 10^{-3}, 10^{-4}\}$ | $\{10^{-3}, 10^{-4}\}$ | $\{10^{-3}, 10^{-4}\}$ | $\{10^{-3}, 10^{-4}\}$ | $\{10^{-3}, 10^{-4}\}$ | $\{10^{-3}, 3{\times}10^{-4}, 10^{-4}\}$ |
| Batch size | 64 | $\{256, 512\}$ | 256 | 256 | 512 | 512 | $\{512, 1024\}$ |
| Weight decay | 0.01 | $\{0, 0.01\}$ | $\{0, 0.01\}$ | $\{0, 0.01\}$ | $\{0, 0.01\}$ | $\{0, 0.01\}$ | 0 |
| Dropout | 0.1 | $\{0, 0.1\}$ | 0 | 0 | $\{0, 0.1\}$ | $\{0, 0.1\}$ | $\{0, 0.1\}$ |
| Validation criterion | Loss | AUROC | AUPRC, any interaction, residual scores | Spearman | MSE | MSE | MSE |
| Splits | Training / validation 99/1 | 10 partitions per regime and cell line | 10 subsamples, library genes 80/10/10 | 1 split, query genes 85/15 | 10 folds per cell line, target-set groups 70/15/15 | 10 folds per cell line, target-set groups 70/15/15 | 1 split, target-set groups 85/15, no test fold |

**Table S3: Gene and Gene Ontology panels used in the embedding space analyses.** Lineage recovery used the three gene groups and the 11 lineages. GO recovery used the 33 parent terms that passed the size and balance filters in all four cell lines. Entries are separated by a centered dot.

| Panel | Members |
|---|---|
| Lineage markers (18) | *PTPRC* · *CD3E* · *CD8A* · *NKG7* · *MS4A1* · *CD79A* · *LYZ* · *CD14* · *TYROBP* · *EPCAM* · *KRT8* · *COL1A1* · *PECAM1* · *VWF* · *HBB* · *GFAP* · *SNAP25* · *MOG* |
| Transcription factors (18) | *SP1* · *YY1* · *MAX* · *CREB1* · *USF1* · *ELK1* · *E2F1* · *ATF4* · *JUN* · *FOS* · *EGR1* · *STAT1* · *STAT3* · *NFKB1* · *CTCF* · *REST* · *KLF4* · *MYC* |
| Housekeeping genes (18) | *ACTB* · *GAPDH* · *B2M* · *TUBB* · *RPL13A* · *HSP90AA1* · *PPIA* · *PGK1* · *YWHAZ* · *SDHA* · *HPRT1* · *UBC* · *RPLP0* · *EEF1A1* · *RPS18* · *GUSB* · *PGAM1* · *VCP* |
| Lineages (11) | Fibroblast and stromal · Epithelial · Retina · Neuron · Endothelial · Myeloid · Erythroid and HSPC · Muscle · T and NK · Glia · B and plasma |
| GO Biological Process (13) | anatomical structure development · animal gross anatomical part developmental process · cell differentiation · developmental process · establishment of localization · intracellular protein localization · localization · localization within membrane · metabolic process · mitotic cell cycle process · regulation of biological quality · response to chemical · transport |
| GO Cellular Component (14) | bounding membrane of organelle · catalytic complex · cytoplasmic vesicle membrane · intracellular membraneless organelle · membrane protein complex · membrane-bounded organelle · nuclear protein-containing complex · plasma membrane bounded cell projection · protein-containing complex · respiratory chain complex · ribonucleoprotein complex · secretory granule · transferase complex · transmembrane transporter complex |
| GO Molecular Function (6) | catalytic activity · catalytic activity, acting on a protein · enzyme binding · hydrolase activity · metal ion binding · transferase activity |

**Table S4: Cell types used as mismatched contexts.** For each target cell line, the CellxGene cell type whose pseudobulk expression had a Pearson correlation closest to 0.5 with the target.

| Task | Cell line | Mismatched cell type | Pearson |
|---|---|---|---|
| Perturbation response | HepG2 | endothelial cell | 0.500 |
| | Jurkat | kidney loop of Henle thin descending limb epithelial cell | 0.512 |
| | K562 | alveolar capillary type 1 endothelial cell | 0.499 |
| | RPE1 | trabecular meshwork cell | 0.491 |
| Genetic interaction | HAP1 | fast muscle cell | 0.498 |
| Drug response, LINCS L1000 | A375 | neutrophil | 0.499 |
| | A549 | kidney interstitial alternatively activated macrophage | 0.503 |
| | HT29 | tracheobronchial goblet cell | 0.500 |
| | MCF7 | kidney interstitial alternatively activated macrophage | 0.498 |
| Drug response, Tahoe-100M | A172 | centrilobular region hepatocyte | 0.501 |
| | BT-474 | kidney interstitial alternatively activated macrophage | 0.500 |
| | NCI-H460 | kidney interstitial alternatively activated macrophage | 0.496 |
| | SW480 | blood vessel endothelial cell | 0.501 |

**Table S5: Extended perturbation-prediction benchmark, per cell line.** Test AUPRC and AUROC (pooled) and their macro-per-perturbation counterparts for EMBER-PERT and all baselines, across the three generalization regimes and the four essential-gene Perturb-seq cell lines. Values are mean$_{\pm \text{s.d.}}$ across the ten random splits. For each cell line, the single best value in each column is shown in **bold**. Higher is better for every metric.

| Group | Model | AUPRC | Macro AUPRC | AUROC | Macro AUROC |
|---|---|---|---|---|---|
| **Regime: Unseen cell line, seen perturbations** | | | | | |
| *Cell line: Jurkat* | | | | | |
| EMBER | EMBER | $\mathbf{0.485}_{\pm 0.004}$ | $\mathbf{0.525}_{\pm 0.005}$ | $\mathbf{0.651}_{\pm 0.004}$ | $\mathbf{0.641}_{\pm 0.005}$ |
| Ablations | EMBER (CSCL-Only) | $0.467_{\pm 0.007}$ | $0.509_{\pm 0.006}$ | $0.637_{\pm 0.006}$ | $0.621_{\pm 0.007}$ |
| | EMBER (Static) | $0.442_{\pm 0.003}$ | $0.482_{\pm 0.006}$ | $0.615_{\pm 0.003}$ | $0.613_{\pm 0.004}$ |
| | EMBER (Recon-Only) | $0.434_{\pm 0.005}$ | $0.485_{\pm 0.005}$ | $0.614_{\pm 0.004}$ | $0.608_{\pm 0.005}$ |
| Prior Knowledge | STRING (GNN)+scGPT | $0.416_{\pm 0.005}$ | $0.459_{\pm 0.005}$ | $0.595_{\pm 0.004}$ | $0.596_{\pm 0.005}$ |
| | GenePT | $0.406_{\pm 0.002}$ | $0.438_{\pm 0.002}$ | $0.576_{\pm 0.003}$ | $0.576_{\pm 0.003}$ |
| | STRING (GNN) | $0.406_{\pm 0.003}$ | $0.447_{\pm 0.004}$ | $0.577_{\pm 0.002}$ | $0.583_{\pm 0.002}$ |
| | STRING (GNN)+Expr | $0.370_{\pm 0.010}$ | $0.409_{\pm 0.011}$ | $0.546_{\pm 0.007}$ | $0.547_{\pm 0.010}$ |
| | STRING (Hop) | $0.341_{\pm 0.001}$ | $0.370_{\pm 0.001}$ | $0.500_{\pm 0.003}$ | $0.517_{\pm 0.003}$ |
| | PINNACLE | $0.336_{\pm 0.001}$ | $0.339_{\pm 0.002}$ | $0.503_{\pm 0.002}$ | $0.503_{\pm 0.002}$ |
| | STRING (Direct) | $0.336_{\pm 0.000}$ | $0.359_{\pm 0.001}$ | $0.504_{\pm 0.001}$ | $0.518_{\pm 0.001}$ |
| Cell | scGPT-Pan-Cancer | $0.427_{\pm 0.006}$ | $0.465_{\pm 0.006}$ | $0.601_{\pm 0.003}$ | $0.596_{\pm 0.005}$ |
| | Geneformer-V2 104M | $0.425_{\pm 0.006}$ | $0.471_{\pm 0.004}$ | $0.605_{\pm 0.005}$ | $0.605_{\pm 0.004}$ |
| | Geneformer-V2 316M | $0.413_{\pm 0.004}$ | $0.446_{\pm 0.006}$ | $0.589_{\pm 0.002}$ | $0.584_{\pm 0.005}$ |
| | AIDO.Cell 3M | $0.407_{\pm 0.003}$ | $0.449_{\pm 0.003}$ | $0.586_{\pm 0.002}$ | $0.584_{\pm 0.003}$ |
| | Gene2Vec | $0.406_{\pm 0.003}$ | $0.442_{\pm 0.005}$ | $0.582_{\pm 0.003}$ | $0.580_{\pm 0.005}$ |
| | scGPT-Whole-Human | $0.403_{\pm 0.006}$ | $0.434_{\pm 0.006}$ | $0.577_{\pm 0.005}$ | $0.561_{\pm 0.008}$ |
| | Coexpression | $0.352_{\pm 0.005}$ | $0.406_{\pm 0.007}$ | $0.524_{\pm 0.007}$ | $0.518_{\pm 0.008}$ |
| | TranscriptFormer | $0.345_{\pm 0.002}$ | $0.366_{\pm 0.008}$ | $0.516_{\pm 0.003}$ | $0.497_{\pm 0.007}$ |
| | scPRINT-2 | $0.336_{\pm 0.001}$ | $0.371_{\pm 0.003}$ | $0.503_{\pm 0.002}$ | $0.507_{\pm 0.003}$ |
| Sequence | ProTrek | $0.414_{\pm 0.003}$ | $0.454_{\pm 0.005}$ | $0.583_{\pm 0.002}$ | $0.588_{\pm 0.005}$ |
| | ESM2 | $0.403_{\pm 0.004}$ | $0.445_{\pm 0.003}$ | $0.573_{\pm 0.003}$ | $0.579_{\pm 0.003}$ |
| | HyenaDNA | $0.368_{\pm 0.003}$ | $0.408_{\pm 0.004}$ | $0.544_{\pm 0.003}$ | $0.549_{\pm 0.004}$ |
| Perturbation model | STATE | $0.341_{\pm 0.010}$ | $0.457_{\pm 0.006}$ | $0.505_{\pm 0.008}$ | $0.521_{\pm 0.008}$ |
| Trivial / Random | Context Mean | $0.398_{\pm 0.002}$ | $0.428_{\pm 0.003}$ | $0.564_{\pm 0.001}$ | $0.575_{\pm 0.002}$ |

*continued on next page*

*Table S5 (continued)*

| Group | Model | AUPRC | Macro AUPRC | AUROC | Macro AUROC |
|---|---|---|---|---|---|
| | Random (Noise Floor) | 0.354$_{\pm 0.004}$ | 0.394$_{\pm 0.006}$ | 0.528$_{\pm 0.004}$ | 0.533$_{\pm 0.006}$ |
| | Train Mean | 0.335$_{\pm 0.001}$ | 0.378$_{\pm 0.002}$ | 0.503$_{\pm 0.001}$ | 0.511$_{\pm 0.002}$ |
| *Cell line: K562* | | | | | |
| EMBER | EMBER | **0.457**$_{\pm 0.008}$ | **0.495**$_{\pm 0.007}$ | **0.624**$_{\pm 0.004}$ | **0.617**$_{\pm 0.004}$ |
| Ablations | EMBER (CSCL-Only) | 0.443$_{\pm 0.005}$ | 0.484$_{\pm 0.005}$ | 0.607$_{\pm 0.005}$ | 0.605$_{\pm 0.004}$ |
| | EMBER (Static) | 0.434$_{\pm 0.003}$ | 0.475$_{\pm 0.005}$ | 0.601$_{\pm 0.003}$ | 0.605$_{\pm 0.003}$ |
| | EMBER (Recon-Only) | 0.423$_{\pm 0.006}$ | 0.461$_{\pm 0.005}$ | 0.596$_{\pm 0.004}$ | 0.591$_{\pm 0.005}$ |
| Prior Knowledge | STRING (GNN)+scGPT | 0.425$_{\pm 0.007}$ | 0.467$_{\pm 0.007}$ | 0.594$_{\pm 0.004}$ | 0.600$_{\pm 0.005}$ |
| | STRING (GNN) | 0.418$_{\pm 0.005}$ | 0.464$_{\pm 0.006}$ | 0.584$_{\pm 0.004}$ | 0.595$_{\pm 0.005}$ |
| | GenePT | 0.417$_{\pm 0.005}$ | 0.452$_{\pm 0.006}$ | 0.584$_{\pm 0.003}$ | 0.585$_{\pm 0.004}$ |
| | STRING (GNN)+Expr | 0.370$_{\pm 0.009}$ | 0.411$_{\pm 0.015}$ | 0.545$_{\pm 0.007}$ | 0.546$_{\pm 0.013}$ |
| | STRING (Hop) | 0.350$_{\pm 0.001}$ | 0.383$_{\pm 0.003}$ | 0.510$_{\pm 0.003}$ | 0.529$_{\pm 0.003}$ |
| | STRING (Direct) | 0.343$_{\pm 0.001}$ | 0.368$_{\pm 0.003}$ | 0.514$_{\pm 0.001}$ | 0.528$_{\pm 0.002}$ |
| | PINNACLE | 0.335$_{\pm 0.001}$ | 0.360$_{\pm 0.004}$ | 0.502$_{\pm 0.001}$ | 0.510$_{\pm 0.003}$ |
| Cell | scGPT-Pan-Cancer | 0.427$_{\pm 0.005}$ | 0.466$_{\pm 0.007}$ | 0.596$_{\pm 0.002}$ | 0.598$_{\pm 0.007}$ |
| | Geneformer-V2 104M | 0.418$_{\pm 0.005}$ | 0.466$_{\pm 0.007}$ | 0.592$_{\pm 0.003}$ | 0.601$_{\pm 0.005}$ |
| | Gene2Vec | 0.406$_{\pm 0.004}$ | 0.448$_{\pm 0.007}$ | 0.581$_{\pm 0.003}$ | 0.583$_{\pm 0.004}$ |
| | Geneformer-V2 316M | 0.404$_{\pm 0.003}$ | 0.450$_{\pm 0.003}$ | 0.582$_{\pm 0.003}$ | 0.586$_{\pm 0.003}$ |
| | scGPT-Whole-Human | 0.402$_{\pm 0.008}$ | 0.440$_{\pm 0.010}$ | 0.580$_{\pm 0.007}$ | 0.578$_{\pm 0.010}$ |
| | AIDO.Cell 3M | 0.397$_{\pm 0.002}$ | 0.445$_{\pm 0.005}$ | 0.571$_{\pm 0.003}$ | 0.577$_{\pm 0.005}$ |
| | Coexpression | 0.346$_{\pm 0.006}$ | 0.395$_{\pm 0.011}$ | 0.516$_{\pm 0.006}$ | 0.517$_{\pm 0.006}$ |
| | scPRINT-2 | 0.336$_{\pm 0.002}$ | 0.376$_{\pm 0.004}$ | 0.503$_{\pm 0.002}$ | 0.507$_{\pm 0.004}$ |
| | TranscriptFormer | 0.335$_{\pm 0.002}$ | 0.362$_{\pm 0.009}$ | 0.503$_{\pm 0.003}$ | 0.495$_{\pm 0.009}$ |
| Sequence | ProTrek | 0.416$_{\pm 0.005}$ | 0.461$_{\pm 0.005}$ | 0.585$_{\pm 0.002}$ | 0.592$_{\pm 0.005}$ |
| | ESM2 | 0.408$_{\pm 0.003}$ | 0.455$_{\pm 0.004}$ | 0.578$_{\pm 0.003}$ | 0.588$_{\pm 0.003}$ |
| | HyenaDNA | 0.373$_{\pm 0.004}$ | 0.420$_{\pm 0.005}$ | 0.548$_{\pm 0.003}$ | 0.557$_{\pm 0.004}$ |
| Perturbation model | STATE | 0.424$_{\pm 0.008}$ | 0.465$_{\pm 0.008}$ | 0.556$_{\pm 0.005}$ | 0.547$_{\pm 0.005}$ |
| Trivial / Random | Context Mean | 0.423$_{\pm 0.003}$ | 0.461$_{\pm 0.004}$ | 0.574$_{\pm 0.002}$ | 0.592$_{\pm 0.003}$ |
| | Random (Noise Floor) | 0.360$_{\pm 0.006}$ | 0.408$_{\pm 0.007}$ | 0.534$_{\pm 0.007}$ | 0.543$_{\pm 0.007}$ |

*continued on next page*

*Table S5 (continued)*

| Group | Model | AUPRC | Macro AUPRC | AUROC | Macro AUROC |
|---|---|---|---|---|---|
| | Train Mean | $0.334_{\pm 0.002}$ | $0.372_{\pm 0.003}$ | $0.501_{\pm 0.002}$ | $0.495_{\pm 0.003}$ |
| *Cell line: HepG2* | | | | | |
| EMBER | EMBER | $\mathbf{0.484}_{\pm 0.009}$ | $\mathbf{0.525}_{\pm 0.005}$ | $\mathbf{0.664}_{\pm 0.007}$ | $\mathbf{0.643}_{\pm 0.006}$ |
| Ablations | EMBER (Recon-Only) | $0.454_{\pm 0.010}$ | $0.489_{\pm 0.008}$ | $0.629_{\pm 0.008}$ | $0.614_{\pm 0.007}$ |
| | EMBER (CSCL-Only) | $0.452_{\pm 0.007}$ | $0.502_{\pm 0.004}$ | $0.635_{\pm 0.007}$ | $0.615_{\pm 0.005}$ |
| | EMBER (Static) | $0.443_{\pm 0.010}$ | $0.483_{\pm 0.007}$ | $0.627_{\pm 0.007}$ | $0.615_{\pm 0.006}$ |
| Prior Knowledge | STRING (GNN)+scGPT | $0.450_{\pm 0.010}$ | $0.491_{\pm 0.004}$ | $0.632_{\pm 0.005}$ | $0.622_{\pm 0.003}$ |
| | STRING (GNN) | $0.421_{\pm 0.010}$ | $0.470_{\pm 0.006}$ | $0.610_{\pm 0.007}$ | $0.603_{\pm 0.006}$ |
| | GenePT | $0.408_{\pm 0.004}$ | $0.444_{\pm 0.005}$ | $0.586_{\pm 0.003}$ | $0.579_{\pm 0.003}$ |
| | STRING (GNN)+Expr | $0.406_{\pm 0.013}$ | $0.444_{\pm 0.013}$ | $0.585_{\pm 0.007}$ | $0.585_{\pm 0.007}$ |
| | STRING (Hop) | $0.343_{\pm 0.001}$ | $0.363_{\pm 0.002}$ | $0.504_{\pm 0.002}$ | $0.509_{\pm 0.003}$ |
| | STRING (Direct) | $0.336_{\pm 0.000}$ | $0.355_{\pm 0.002}$ | $0.506_{\pm 0.001}$ | $0.511_{\pm 0.001}$ |
| | PINNACLE | $0.335_{\pm 0.001}$ | $0.345_{\pm 0.004}$ | $0.503_{\pm 0.002}$ | $0.505_{\pm 0.004}$ |
| Cell | scGPT-Whole-Human | $0.451_{\pm 0.012}$ | $0.473_{\pm 0.009}$ | $0.625_{\pm 0.006}$ | $0.610_{\pm 0.007}$ |
| | scGPT-Pan-Cancer | $0.445_{\pm 0.010}$ | $0.480_{\pm 0.007}$ | $0.624_{\pm 0.007}$ | $0.611_{\pm 0.006}$ |
| | Geneformer-V2 104M | $0.434_{\pm 0.006}$ | $0.480_{\pm 0.005}$ | $0.618_{\pm 0.005}$ | $0.609_{\pm 0.005}$ |
| | Geneformer-V2 316M | $0.417_{\pm 0.006}$ | $0.466_{\pm 0.005}$ | $0.601_{\pm 0.005}$ | $0.596_{\pm 0.005}$ |
| | Gene2Vec | $0.413_{\pm 0.008}$ | $0.461_{\pm 0.004}$ | $0.600_{\pm 0.005}$ | $0.592_{\pm 0.003}$ |
| | AIDO.Cell 3M | $0.410_{\pm 0.008}$ | $0.454_{\pm 0.008}$ | $0.592_{\pm 0.008}$ | $0.583_{\pm 0.007}$ |
| | Coexpression | $0.343_{\pm 0.007}$ | $0.403_{\pm 0.012}$ | $0.511_{\pm 0.011}$ | $0.513_{\pm 0.011}$ |
| | scPRINT-2 | $0.335_{\pm 0.002}$ | $0.367_{\pm 0.003}$ | $0.502_{\pm 0.002}$ | $0.503_{\pm 0.005}$ |
| | TranscriptFormer | $0.335_{\pm 0.005}$ | $0.368_{\pm 0.008}$ | $0.502_{\pm 0.006}$ | $0.503_{\pm 0.007}$ |
| Sequence | ProTrek | $0.418_{\pm 0.005}$ | $0.459_{\pm 0.004}$ | $0.599_{\pm 0.005}$ | $0.593_{\pm 0.005}$ |
| | ESM2 | $0.411_{\pm 0.005}$ | $0.453_{\pm 0.006}$ | $0.587_{\pm 0.006}$ | $0.588_{\pm 0.006}$ |
| | HyenaDNA | $0.370_{\pm 0.003}$ | $0.405_{\pm 0.004}$ | $0.541_{\pm 0.002}$ | $0.545_{\pm 0.004}$ |
| Perturbation model | STATE | $0.421_{\pm 0.020}$ | $0.466_{\pm 0.017}$ | $0.572_{\pm 0.014}$ | $0.545_{\pm 0.011}$ |
| Trivial / Random | Context Mean | $0.397_{\pm 0.002}$ | $0.429_{\pm 0.002}$ | $0.555_{\pm 0.004}$ | $0.568_{\pm 0.003}$ |
| | Random (Noise Floor) | $0.359_{\pm 0.004}$ | $0.396_{\pm 0.005}$ | $0.531_{\pm 0.005}$ | $0.535_{\pm 0.005}$ |
| | Train Mean | $0.334_{\pm 0.002}$ | $0.367_{\pm 0.002}$ | $0.501_{\pm 0.002}$ | $0.496_{\pm 0.002}$ |

*continued on next page*

*Table S5 (continued)*

| Group | Model | AUPRC | Macro AUPRC | AUROC | Macro AUROC |
|---|---|---|---|---|---|
| *Cell line: RPE1* | | | | | |
| EMBER | EMBER | $\mathbf{0.517}_{\pm 0.007}$ | $\mathbf{0.540}_{\pm 0.007}$ | $\mathbf{0.673}_{\pm 0.005}$ | $\mathbf{0.644}_{\pm 0.007}$ |
| Ablations | EMBER (CSCL-Only) | $0.494_{\pm 0.005}$ | $0.529_{\pm 0.004}$ | $0.654_{\pm 0.005}$ | $0.622_{\pm 0.006}$ |
| | EMBER (Static) | $0.480_{\pm 0.004}$ | $0.510_{\pm 0.004}$ | $0.641_{\pm 0.003}$ | $0.623_{\pm 0.006}$ |
| | EMBER (Recon-Only) | $0.472_{\pm 0.010}$ | $0.502_{\pm 0.007}$ | $0.638_{\pm 0.008}$ | $0.609_{\pm 0.008}$ |
| Prior Knowledge | STRING (GNN)+scGPT | $0.466_{\pm 0.011}$ | $0.490_{\pm 0.009}$ | $0.626_{\pm 0.011}$ | $0.605_{\pm 0.011}$ |
| | STRING (GNN) | $0.453_{\pm 0.004}$ | $0.491_{\pm 0.005}$ | $0.613_{\pm 0.004}$ | $0.603_{\pm 0.004}$ |
| | GenePT | $0.436_{\pm 0.006}$ | $0.469_{\pm 0.006}$ | $0.602_{\pm 0.005}$ | $0.588_{\pm 0.006}$ |
| | STRING (GNN)+Expr | $0.401_{\pm 0.011}$ | $0.452_{\pm 0.015}$ | $0.569_{\pm 0.009}$ | $0.565_{\pm 0.014}$ |
| | STRING (Hop) | $0.352_{\pm 0.003}$ | $0.382_{\pm 0.003}$ | $0.516_{\pm 0.004}$ | $0.535_{\pm 0.004}$ |
| | STRING (Direct) | $0.343_{\pm 0.002}$ | $0.371_{\pm 0.003}$ | $0.516_{\pm 0.003}$ | $0.529_{\pm 0.003}$ |
| | PINNACLE | $0.334_{\pm 0.000}$ | $0.335_{\pm 0.001}$ | $0.500_{\pm 0.001}$ | $0.500_{\pm 0.000}$ |
| Cell | Geneformer-V2 104M | $0.465_{\pm 0.007}$ | $0.496_{\pm 0.006}$ | $0.629_{\pm 0.007}$ | $0.606_{\pm 0.007}$ |
| | scGPT-Pan-Cancer | $0.463_{\pm 0.010}$ | $0.495_{\pm 0.010}$ | $0.629_{\pm 0.009}$ | $0.611_{\pm 0.011}$ |
| | scGPT-Whole-Human | $0.460_{\pm 0.006}$ | $0.501_{\pm 0.007}$ | $0.634_{\pm 0.006}$ | $0.617_{\pm 0.008}$ |
| | Gene2Vec | $0.452_{\pm 0.005}$ | $0.485_{\pm 0.004}$ | $0.619_{\pm 0.006}$ | $0.596_{\pm 0.005}$ |
| | AIDO.Cell 3M | $0.449_{\pm 0.005}$ | $0.481_{\pm 0.006}$ | $0.615_{\pm 0.006}$ | $0.593_{\pm 0.006}$ |
| | Geneformer-V2 316M | $0.445_{\pm 0.006}$ | $0.477_{\pm 0.004}$ | $0.614_{\pm 0.006}$ | $0.588_{\pm 0.008}$ |
| | Coexpression | $0.353_{\pm 0.007}$ | $0.407_{\pm 0.011}$ | $0.528_{\pm 0.007}$ | $0.505_{\pm 0.014}$ |
| | TranscriptFormer | $0.341_{\pm 0.006}$ | $0.369_{\pm 0.023}$ | $0.508_{\pm 0.009}$ | $0.481_{\pm 0.023}$ |
| | scPRINT-2 | $0.337_{\pm 0.001}$ | $0.371_{\pm 0.002}$ | $0.502_{\pm 0.001}$ | $0.502_{\pm 0.003}$ |
| Sequence | ProTrek | $0.448_{\pm 0.005}$ | $0.484_{\pm 0.003}$ | $0.609_{\pm 0.003}$ | $0.595_{\pm 0.003}$ |
| | ESM2 | $0.435_{\pm 0.005}$ | $0.471_{\pm 0.004}$ | $0.593_{\pm 0.003}$ | $0.583_{\pm 0.003}$ |
| | HyenaDNA | $0.401_{\pm 0.004}$ | $0.441_{\pm 0.004}$ | $0.571_{\pm 0.005}$ | $0.567_{\pm 0.005}$ |
| Perturbation model | STATE | $0.369_{\pm 0.017}$ | $0.473_{\pm 0.007}$ | $0.527_{\pm 0.010}$ | $0.549_{\pm 0.006}$ |
| Trivial / Random | Context Mean | $0.432_{\pm 0.003}$ | $0.447_{\pm 0.002}$ | $0.585_{\pm 0.003}$ | $0.574_{\pm 0.002}$ |
| | Random (Noise Floor) | $0.377_{\pm 0.005}$ | $0.418_{\pm 0.006}$ | $0.545_{\pm 0.005}$ | $0.548_{\pm 0.006}$ |
| | Train Mean | $0.333_{\pm 0.002}$ | $0.388_{\pm 0.003}$ | $0.500_{\pm 0.002}$ | $0.513_{\pm 0.003}$ |
| **Regime: Unseen cell line, unseen perturbations** | | | | | |

*continued on next page*

*Table S5 (continued)*

| Group | Model | AUPRC | Macro AUPRC | AUROC | Macro AUROC |
|---|---|---|---|---|---|
| *Cell line: Jurkat* | | | | | |
| EMBER | EMBER | **0.430**$_{\pm0.013}$ | **0.481**$_{\pm0.010}$ | **0.609**$_{\pm0.014}$ | **0.601**$_{\pm0.009}$ |
| Ablations | EMBER (CSCL-Only) | 0.421$_{\pm0.016}$ | 0.471$_{\pm0.012}$ | 0.599$_{\pm0.011}$ | 0.586$_{\pm0.013}$ |
| | EMBER (Static) | 0.389$_{\pm0.008}$ | 0.428$_{\pm0.006}$ | 0.569$_{\pm0.008}$ | 0.563$_{\pm0.007}$ |
| | EMBER (Recon-Only) | 0.357$_{\pm0.009}$ | 0.420$_{\pm0.011}$ | 0.533$_{\pm0.011}$ | 0.537$_{\pm0.011}$ |
| Prior Knowledge | STRING (GNN)+scGPT | 0.375$_{\pm0.004}$ | 0.413$_{\pm0.005}$ | 0.556$_{\pm0.004}$ | 0.552$_{\pm0.006}$ |
| | GenePT | 0.370$_{\pm0.010}$ | 0.397$_{\pm0.009}$ | 0.542$_{\pm0.011}$ | 0.536$_{\pm0.010}$ |
| | STRING (GNN) | 0.370$_{\pm0.004}$ | 0.401$_{\pm0.007}$ | 0.547$_{\pm0.004}$ | 0.546$_{\pm0.006}$ |
| | STRING (GNN)+Expr | 0.353$_{\pm0.007}$ | 0.384$_{\pm0.011}$ | 0.529$_{\pm0.009}$ | 0.527$_{\pm0.008}$ |
| | STRING (Hop) | 0.340$_{\pm0.004}$ | 0.365$_{\pm0.007}$ | 0.499$_{\pm0.009}$ | 0.511$_{\pm0.009}$ |
| | STRING (Direct) | 0.335$_{\pm0.001}$ | 0.355$_{\pm0.006}$ | 0.503$_{\pm0.002}$ | 0.514$_{\pm0.004}$ |
| | PINNACLE | 0.334$_{\pm0.002}$ | 0.337$_{\pm0.003}$ | 0.500$_{\pm0.003}$ | 0.500$_{\pm0.003}$ |
| Cell | scGPT-Pan-Cancer | 0.357$_{\pm0.010}$ | 0.397$_{\pm0.015}$ | 0.533$_{\pm0.011}$ | 0.527$_{\pm0.012}$ |
| | scGPT-Whole-Human | 0.352$_{\pm0.010}$ | 0.384$_{\pm0.014}$ | 0.525$_{\pm0.013}$ | 0.507$_{\pm0.010}$ |
| | Geneformer-V2 104M | 0.350$_{\pm0.008}$ | 0.388$_{\pm0.014}$ | 0.524$_{\pm0.008}$ | 0.519$_{\pm0.013}$ |
| | Geneformer-V2 316M | 0.344$_{\pm0.008}$ | 0.379$_{\pm0.007}$ | 0.517$_{\pm0.010}$ | 0.512$_{\pm0.010}$ |
| | Coexpression | 0.343$_{\pm0.012}$ | 0.398$_{\pm0.018}$ | 0.512$_{\pm0.013}$ | 0.515$_{\pm0.014}$ |
| | Gene2Vec | 0.342$_{\pm0.003}$ | 0.374$_{\pm0.009}$ | 0.513$_{\pm0.004}$ | 0.513$_{\pm0.007}$ |
| | TranscriptFormer | 0.341$_{\pm0.008}$ | 0.365$_{\pm0.009}$ | 0.510$_{\pm0.012}$ | 0.495$_{\pm0.011}$ |
| | scPRINT-2 | 0.335$_{\pm0.003}$ | 0.367$_{\pm0.005}$ | 0.501$_{\pm0.004}$ | 0.503$_{\pm0.006}$ |
| | AIDO.Cell 3M | 0.334$_{\pm0.009}$ | 0.373$_{\pm0.006}$ | 0.501$_{\pm0.011}$ | 0.501$_{\pm0.011}$ |
| Sequence | ProTrek | 0.367$_{\pm0.006}$ | 0.396$_{\pm0.010}$ | 0.540$_{\pm0.005}$ | 0.535$_{\pm0.007}$ |
| | ESM2 | 0.352$_{\pm0.008}$ | 0.383$_{\pm0.015}$ | 0.525$_{\pm0.009}$ | 0.525$_{\pm0.012}$ |
| | HyenaDNA | 0.333$_{\pm0.003}$ | 0.364$_{\pm0.007}$ | 0.501$_{\pm0.004}$ | 0.503$_{\pm0.007}$ |
| Trivial / Random | Random (Noise Floor) | 0.335$_{\pm0.006}$ | 0.361$_{\pm0.007}$ | 0.502$_{\pm0.008}$ | 0.500$_{\pm0.007}$ |
| | Train Mean | 0.335$_{\pm0.003}$ | 0.375$_{\pm0.006}$ | 0.501$_{\pm0.005}$ | 0.507$_{\pm0.006}$ |
| *Cell line: K562* | | | | | |
| EMBER | EMBER | **0.415**$_{\pm0.012}$ | **0.449**$_{\pm0.009}$ | **0.587**$_{\pm0.012}$ | **0.573**$_{\pm0.009}$ |
| Ablations | EMBER (CSCL-Only) | 0.405$_{\pm0.012}$ | 0.435$_{\pm0.014}$ | 0.572$_{\pm0.010}$ | 0.556$_{\pm0.013}$ |

*continued on next page*

*Table S5 (continued)*

| Group | Model | AUPRC | Macro AUPRC | AUROC | Macro AUROC |
|---|---|---|---|---|---|
| | EMBER (Static) | $0.385_{\pm 0.008}$ | $0.412_{\pm 0.011}$ | $0.558_{\pm 0.009}$ | $0.545_{\pm 0.010}$ |
| | EMBER (Recon-Only) | $0.355_{\pm 0.007}$ | $0.397_{\pm 0.010}$ | $0.526_{\pm 0.008}$ | $0.521_{\pm 0.014}$ |
| Prior Knowledge | STRING (GNN)+scGPT | $0.387_{\pm 0.008}$ | $0.417_{\pm 0.013}$ | $0.560_{\pm 0.006}$ | $0.553_{\pm 0.007}$ |
| | STRING (GNN) | $0.386_{\pm 0.006}$ | $0.419_{\pm 0.013}$ | $0.553_{\pm 0.006}$ | $0.552_{\pm 0.008}$ |
| | GenePT | $0.373_{\pm 0.007}$ | $0.403_{\pm 0.007}$ | $0.540_{\pm 0.004}$ | $0.536_{\pm 0.006}$ |
| | STRING (GNN)+Expr | $0.359_{\pm 0.007}$ | $0.393_{\pm 0.011}$ | $0.531_{\pm 0.007}$ | $0.528_{\pm 0.007}$ |
| | STRING (Hop) | $0.353_{\pm 0.003}$ | $0.385_{\pm 0.007}$ | $0.516_{\pm 0.005}$ | $0.532_{\pm 0.006}$ |
| | STRING (Direct) | $0.344_{\pm 0.002}$ | $0.371_{\pm 0.007}$ | $0.516_{\pm 0.002}$ | $0.531_{\pm 0.005}$ |
| | PINNACLE | $0.335_{\pm 0.001}$ | $0.354_{\pm 0.009}$ | $0.502_{\pm 0.002}$ | $0.506_{\pm 0.008}$ |
| Cell | scGPT-Pan-Cancer | $0.367_{\pm 0.005}$ | $0.398_{\pm 0.008}$ | $0.538_{\pm 0.006}$ | $0.528_{\pm 0.011}$ |
| | Geneformer-V2 104M | $0.351_{\pm 0.005}$ | $0.387_{\pm 0.006}$ | $0.523_{\pm 0.006}$ | $0.520_{\pm 0.007}$ |
| | scGPT-Whole-Human | $0.350_{\pm 0.008}$ | $0.381_{\pm 0.010}$ | $0.521_{\pm 0.008}$ | $0.512_{\pm 0.008}$ |
| | Gene2Vec | $0.344_{\pm 0.003}$ | $0.376_{\pm 0.005}$ | $0.515_{\pm 0.005}$ | $0.508_{\pm 0.007}$ |
| | Geneformer-V2 316M | $0.343_{\pm 0.005}$ | $0.371_{\pm 0.011}$ | $0.513_{\pm 0.008}$ | $0.509_{\pm 0.011}$ |
| | Coexpression | $0.342_{\pm 0.010}$ | $0.391_{\pm 0.014}$ | $0.512_{\pm 0.014}$ | $0.516_{\pm 0.012}$ |
| | AIDO.Cell 3M | $0.339_{\pm 0.006}$ | $0.375_{\pm 0.007}$ | $0.504_{\pm 0.007}$ | $0.503_{\pm 0.007}$ |
| | scPRINT-2 | $0.337_{\pm 0.005}$ | $0.374_{\pm 0.010}$ | $0.504_{\pm 0.006}$ | $0.506_{\pm 0.012}$ |
| | TranscriptFormer | $0.333_{\pm 0.004}$ | $0.360_{\pm 0.014}$ | $0.501_{\pm 0.005}$ | $0.491_{\pm 0.013}$ |
| Sequence | ESM2 | $0.367_{\pm 0.008}$ | $0.401_{\pm 0.008}$ | $0.536_{\pm 0.006}$ | $0.534_{\pm 0.012}$ |
| | ProTrek | $0.365_{\pm 0.006}$ | $0.398_{\pm 0.008}$ | $0.538_{\pm 0.004}$ | $0.531_{\pm 0.008}$ |
| | HyenaDNA | $0.336_{\pm 0.003}$ | $0.373_{\pm 0.007}$ | $0.501_{\pm 0.004}$ | $0.502_{\pm 0.006}$ |
| Trivial / Random | Random (Noise Floor) | $0.335_{\pm 0.003}$ | $0.367_{\pm 0.003}$ | $0.502_{\pm 0.006}$ | $0.499_{\pm 0.005}$ |
| | Train Mean | $0.332_{\pm 0.004}$ | $0.369_{\pm 0.007}$ | $0.496_{\pm 0.004}$ | $0.486_{\pm 0.008}$ |
| *Cell line: HepG2* | | | | | |
| EMBER | EMBER | $\mathbf{0.422}_{\pm 0.024}$ | $\mathbf{0.474}_{\pm 0.014}$ | $\mathbf{0.602}_{\pm 0.018}$ | $\mathbf{0.589}_{\pm 0.016}$ |
| Ablations | EMBER (CSCL-Only) | $0.396_{\pm 0.019}$ | $0.449_{\pm 0.012}$ | $0.577_{\pm 0.020}$ | $0.560_{\pm 0.019}$ |
| | EMBER (Static) | $0.377_{\pm 0.019}$ | $0.421_{\pm 0.013}$ | $0.562_{\pm 0.019}$ | $0.553_{\pm 0.017}$ |
| | EMBER (Recon-Only) | $0.365_{\pm 0.017}$ | $0.420_{\pm 0.012}$ | $0.537_{\pm 0.018}$ | $0.534_{\pm 0.015}$ |
| Prior Knowledge | STRING (GNN)+scGPT | $0.389_{\pm 0.012}$ | $0.434_{\pm 0.009}$ | $0.576_{\pm 0.009}$ | $0.563_{\pm 0.011}$ |

*continued on next page*

*Table S5 (continued)*

| Group | Model | AUPRC | Macro AUPRC | AUROC | Macro AUROC |
|---|---|---|---|---|---|
| | STRING (GNN) | 0.384$_{\pm 0.007}$ | 0.426$_{\pm 0.011}$ | 0.576$_{\pm 0.009}$ | 0.564$_{\pm 0.011}$ |
| | STRING (GNN)+Expr | 0.379$_{\pm 0.015}$ | 0.418$_{\pm 0.016}$ | 0.561$_{\pm 0.013}$ | 0.561$_{\pm 0.017}$ |
| | GenePT | 0.365$_{\pm 0.013}$ | 0.393$_{\pm 0.015}$ | 0.542$_{\pm 0.013}$ | 0.534$_{\pm 0.012}$ |
| | STRING (Hop) | 0.345$_{\pm 0.003}$ | 0.363$_{\pm 0.007}$ | 0.507$_{\pm 0.005}$ | 0.510$_{\pm 0.007}$ |
| | STRING (Direct) | 0.338$_{\pm 0.002}$ | 0.355$_{\pm 0.006}$ | 0.508$_{\pm 0.003}$ | 0.512$_{\pm 0.006}$ |
| | PINNACLE | 0.335$_{\pm 0.003}$ | 0.344$_{\pm 0.006}$ | 0.502$_{\pm 0.005}$ | 0.503$_{\pm 0.008}$ |
| Cell | scGPT-Whole-Human | 0.376$_{\pm 0.022}$ | 0.414$_{\pm 0.015}$ | 0.549$_{\pm 0.023}$ | 0.548$_{\pm 0.021}$ |
| | scGPT-Pan-Cancer | 0.362$_{\pm 0.011}$ | 0.409$_{\pm 0.011}$ | 0.543$_{\pm 0.017}$ | 0.544$_{\pm 0.013}$ |
| | Geneformer-V2 316M | 0.349$_{\pm 0.008}$ | 0.392$_{\pm 0.009}$ | 0.518$_{\pm 0.007}$ | 0.519$_{\pm 0.009}$ |
| | Coexpression | 0.346$_{\pm 0.016}$ | 0.400$_{\pm 0.015}$ | 0.516$_{\pm 0.021}$ | 0.517$_{\pm 0.019}$ |
| | Geneformer-V2 104M | 0.345$_{\pm 0.011}$ | 0.388$_{\pm 0.012}$ | 0.518$_{\pm 0.014}$ | 0.518$_{\pm 0.014}$ |
| | Gene2Vec | 0.340$_{\pm 0.010}$ | 0.390$_{\pm 0.010}$ | 0.512$_{\pm 0.013}$ | 0.516$_{\pm 0.012}$ |
| | AIDO.Cell 3M | 0.335$_{\pm 0.010}$ | 0.377$_{\pm 0.010}$ | 0.504$_{\pm 0.014}$ | 0.504$_{\pm 0.014}$ |
| | scPRINT-2 | 0.335$_{\pm 0.004}$ | 0.364$_{\pm 0.004}$ | 0.502$_{\pm 0.005}$ | 0.501$_{\pm 0.006}$ |
| | TranscriptFormer | 0.333$_{\pm 0.006}$ | 0.362$_{\pm 0.011}$ | 0.497$_{\pm 0.011}$ | 0.499$_{\pm 0.013}$ |
| Sequence | ProTrek | 0.366$_{\pm 0.013}$ | 0.405$_{\pm 0.009}$ | 0.543$_{\pm 0.011}$ | 0.540$_{\pm 0.012}$ |
| | ESM2 | 0.360$_{\pm 0.009}$ | 0.390$_{\pm 0.013}$ | 0.536$_{\pm 0.011}$ | 0.535$_{\pm 0.010}$ |
| | HyenaDNA | 0.339$_{\pm 0.004}$ | 0.368$_{\pm 0.008}$ | 0.507$_{\pm 0.005}$ | 0.509$_{\pm 0.006}$ |
| Trivial / Random | Random (Noise Floor) | 0.333$_{\pm 0.004}$ | 0.354$_{\pm 0.009}$ | 0.498$_{\pm 0.007}$ | 0.499$_{\pm 0.007}$ |
| | Train Mean | 0.332$_{\pm 0.002}$ | 0.364$_{\pm 0.003}$ | 0.497$_{\pm 0.004}$ | 0.493$_{\pm 0.005}$ |
| *Cell line: RPE1* | | | | | |
| Ember | Ember | **0.432**$_{\pm 0.020}$ | **0.479**$_{\pm 0.015}$ | **0.603**$_{\pm 0.020}$ | **0.580**$_{\pm 0.020}$ |
| Ablations | Ember (CSCL-Only) | 0.415$_{\pm 0.019}$ | 0.467$_{\pm 0.014}$ | 0.585$_{\pm 0.017}$ | 0.561$_{\pm 0.016}$ |
| | Ember (Static) | 0.395$_{\pm 0.018}$ | 0.439$_{\pm 0.015}$ | 0.566$_{\pm 0.013}$ | 0.551$_{\pm 0.015}$ |
| | Ember (Recon-Only) | 0.364$_{\pm 0.016}$ | 0.429$_{\pm 0.017}$ | 0.541$_{\pm 0.018}$ | 0.531$_{\pm 0.018}$ |
| Prior Knowledge | STRING (GNN)+scGPT | 0.393$_{\pm 0.012}$ | 0.434$_{\pm 0.007}$ | 0.564$_{\pm 0.015}$ | 0.545$_{\pm 0.012}$ |
| | STRING (GNN) | 0.389$_{\pm 0.008}$ | 0.435$_{\pm 0.011}$ | 0.560$_{\pm 0.008}$ | 0.552$_{\pm 0.010}$ |
| | GenePT | 0.375$_{\pm 0.013}$ | 0.412$_{\pm 0.017}$ | 0.547$_{\pm 0.013}$ | 0.530$_{\pm 0.020}$ |
| | STRING (GNN)+Expr | 0.370$_{\pm 0.009}$ | 0.423$_{\pm 0.016}$ | 0.546$_{\pm 0.011}$ | 0.540$_{\pm 0.015}$ |

*continued on next page*

*Table S5 (continued)*

| Group | Model | AUPRC | Macro AUPRC | AUROC | Macro AUROC |
|---|---|---|---|---|---|
| | STRING (Hop) | 0.351$_{\pm 0.004}$ | 0.382$_{\pm 0.005}$ | 0.515$_{\pm 0.005}$ | 0.535$_{\pm 0.005}$ |
| | STRING (Direct) | 0.343$_{\pm 0.002}$ | 0.370$_{\pm 0.005}$ | 0.516$_{\pm 0.004}$ | 0.528$_{\pm 0.005}$ |
| | PINNACLE | 0.333$_{\pm 0.001}$ | 0.336$_{\pm 0.001}$ | 0.500$_{\pm 0.001}$ | 0.499$_{\pm 0.001}$ |
| Cell | Geneformer-V2 104M | 0.361$_{\pm 0.014}$ | 0.400$_{\pm 0.010}$ | 0.533$_{\pm 0.008}$ | 0.516$_{\pm 0.012}$ |
| | scGPT-Whole-Human | 0.360$_{\pm 0.016}$ | 0.418$_{\pm 0.026}$ | 0.529$_{\pm 0.018}$ | 0.529$_{\pm 0.027}$ |
| | scGPT-Pan-Cancer | 0.358$_{\pm 0.013}$ | 0.398$_{\pm 0.013}$ | 0.526$_{\pm 0.014}$ | 0.507$_{\pm 0.020}$ |
| | Geneformer-V2 316M | 0.346$_{\pm 0.016}$ | 0.408$_{\pm 0.011}$ | 0.516$_{\pm 0.015}$ | 0.511$_{\pm 0.011}$ |
| | Coexpression | 0.346$_{\pm 0.012}$ | 0.404$_{\pm 0.017}$ | 0.520$_{\pm 0.016}$ | 0.505$_{\pm 0.014}$ |
| | Gene2Vec | 0.344$_{\pm 0.010}$ | 0.408$_{\pm 0.013}$ | 0.511$_{\pm 0.010}$ | 0.508$_{\pm 0.008}$ |
| | AIDO.Cell 3M | 0.339$_{\pm 0.009}$ | 0.401$_{\pm 0.020}$ | 0.509$_{\pm 0.015}$ | 0.507$_{\pm 0.022}$ |
| | scPRINT-2 | 0.336$_{\pm 0.004}$ | 0.370$_{\pm 0.006}$ | 0.503$_{\pm 0.005}$ | 0.505$_{\pm 0.004}$ |
| | TranscriptFormer | 0.333$_{\pm 0.013}$ | 0.385$_{\pm 0.042}$ | 0.500$_{\pm 0.017}$ | 0.488$_{\pm 0.038}$ |
| Sequence | ProTrek | 0.371$_{\pm 0.013}$ | 0.418$_{\pm 0.011}$ | 0.542$_{\pm 0.013}$ | 0.534$_{\pm 0.011}$ |
| | ESM2 | 0.362$_{\pm 0.012}$ | 0.403$_{\pm 0.014}$ | 0.529$_{\pm 0.010}$ | 0.522$_{\pm 0.012}$ |
| | HyenaDNA | 0.334$_{\pm 0.004}$ | 0.374$_{\pm 0.008}$ | 0.500$_{\pm 0.004}$ | 0.503$_{\pm 0.007}$ |
| Trivial / Random | Train Mean | 0.334$_{\pm 0.005}$ | 0.387$_{\pm 0.007}$ | 0.499$_{\pm 0.004}$ | 0.511$_{\pm 0.005}$ |
| | Random (Noise Floor) | 0.334$_{\pm 0.004}$ | 0.367$_{\pm 0.013}$ | 0.499$_{\pm 0.006}$ | 0.504$_{\pm 0.008}$ |
| **Regime: Seen cell line, unseen perturbations** | | | | | |
| *Cell line: Jurkat* | | | | | |
| EMBER | EMBER | **0.482**$_{\pm 0.012}$ | **0.529**$_{\pm 0.005}$ | **0.646**$_{\pm 0.010}$ | **0.625**$_{\pm 0.007}$ |
| Ablations | EMBER (Static) | 0.471$_{\pm 0.007}$ | 0.503$_{\pm 0.005}$ | 0.635$_{\pm 0.005}$ | 0.619$_{\pm 0.005}$ |
| | EMBER (CSCL-Only) | 0.453$_{\pm 0.013}$ | 0.507$_{\pm 0.006}$ | 0.622$_{\pm 0.012}$ | 0.603$_{\pm 0.008}$ |
| | EMBER (Recon-Only) | 0.372$_{\pm 0.009}$ | 0.444$_{\pm 0.007}$ | 0.550$_{\pm 0.011}$ | 0.540$_{\pm 0.007}$ |
| Prior Knowledge | STRING (GNN) | 0.465$_{\pm 0.006}$ | 0.484$_{\pm 0.010}$ | 0.632$_{\pm 0.005}$ | 0.616$_{\pm 0.006}$ |
| | STRING (GNN)+Expr | 0.464$_{\pm 0.007}$ | 0.490$_{\pm 0.005}$ | 0.626$_{\pm 0.006}$ | 0.611$_{\pm 0.007}$ |
| | STRING (GNN)+scGPT | 0.449$_{\pm 0.008}$ | 0.475$_{\pm 0.009}$ | 0.617$_{\pm 0.006}$ | 0.597$_{\pm 0.010}$ |
| | GenePT | 0.414$_{\pm 0.006}$ | 0.435$_{\pm 0.008}$ | 0.586$_{\pm 0.005}$ | 0.564$_{\pm 0.006}$ |
| | STRING (Hop) | 0.345$_{\pm 0.002}$ | 0.382$_{\pm 0.002}$ | 0.506$_{\pm 0.003}$ | 0.529$_{\pm 0.003}$ |
| | STRING (Direct) | 0.338$_{\pm 0.001}$ | 0.368$_{\pm 0.002}$ | 0.508$_{\pm 0.001}$ | 0.527$_{\pm 0.002}$ |

*continued on next page*

*Table S5 (continued)*

| Group | Model | AUPRC | Macro AUPRC | AUROC | Macro AUROC |
|---|---|---|---|---|---|
| | PINNACLE | $0.335_{\pm 0.001}$ | $0.338_{\pm 0.002}$ | $0.502_{\pm 0.002}$ | $0.502_{\pm 0.002}$ |
| Cell | scGPT-Pan-Cancer | $0.390_{\pm 0.008}$ | $0.434_{\pm 0.006}$ | $0.565_{\pm 0.008}$ | $0.543_{\pm 0.006}$ |
| | scGPT-Whole-Human | $0.377_{\pm 0.011}$ | $0.430_{\pm 0.007}$ | $0.553_{\pm 0.010}$ | $0.540_{\pm 0.009}$ |
| | Coexpression | $0.371_{\pm 0.016}$ | $0.451_{\pm 0.016}$ | $0.555_{\pm 0.015}$ | $0.548_{\pm 0.013}$ |
| | Geneformer-V2 104M | $0.361_{\pm 0.006}$ | $0.417_{\pm 0.006}$ | $0.536_{\pm 0.008}$ | $0.533_{\pm 0.005}$ |
| | Gene2Vec | $0.359_{\pm 0.006}$ | $0.405_{\pm 0.007}$ | $0.530_{\pm 0.005}$ | $0.522_{\pm 0.010}$ |
| | Geneformer-V2 316M | $0.349_{\pm 0.004}$ | $0.399_{\pm 0.005}$ | $0.521_{\pm 0.007}$ | $0.512_{\pm 0.007}$ |
| | AIDO.Cell 3M | $0.345_{\pm 0.006}$ | $0.393_{\pm 0.011}$ | $0.515_{\pm 0.007}$ | $0.503_{\pm 0.013}$ |
| | TranscriptFormer | $0.335_{\pm 0.004}$ | $0.373_{\pm 0.013}$ | $0.502_{\pm 0.007}$ | $0.491_{\pm 0.016}$ |
| | scPRINT-2 | $0.333_{\pm 0.002}$ | $0.359_{\pm 0.009}$ | $0.498_{\pm 0.004}$ | $0.490_{\pm 0.008}$ |
| Sequence | ProTrek | $0.415_{\pm 0.010}$ | $0.446_{\pm 0.005}$ | $0.587_{\pm 0.009}$ | $0.570_{\pm 0.007}$ |
| | ESM2 | $0.392_{\pm 0.006}$ | $0.424_{\pm 0.007}$ | $0.562_{\pm 0.006}$ | $0.550_{\pm 0.005}$ |
| | HyenaDNA | $0.332_{\pm 0.005}$ | $0.367_{\pm 0.006}$ | $0.497_{\pm 0.007}$ | $0.491_{\pm 0.007}$ |
| Perturbation model | GEARS | $0.430_{\pm 0.010}$ | $0.449_{\pm 0.006}$ | $0.574_{\pm 0.007}$ | $0.558_{\pm 0.008}$ |
| Trivial / Random | Train Mean | $0.345_{\pm 0.005}$ | $0.377_{\pm 0.005}$ | $0.518_{\pm 0.008}$ | $0.457_{\pm 0.008}$ |
| | Random (Noise Floor) | $0.332_{\pm 0.003}$ | $0.358_{\pm 0.013}$ | $0.499_{\pm 0.004}$ | $0.495_{\pm 0.009}$ |
| *Cell line: K562* | | | | | |
| Ember | Ember | $0.518_{\pm 0.013}$ | $\mathbf{0.526}_{\pm 0.007}$ | $0.668_{\pm 0.007}$ | $0.638_{\pm 0.005}$ |
| Ablations | Ember (Static) | $\mathbf{0.520}_{\pm 0.012}$ | $0.515_{\pm 0.010}$ | $\mathbf{0.673}_{\pm 0.009}$ | $0.640_{\pm 0.008}$ |
| | Ember (CSCL-Only) | $0.466_{\pm 0.009}$ | $0.494_{\pm 0.006}$ | $0.628_{\pm 0.006}$ | $0.608_{\pm 0.006}$ |
| | Ember (Recon-Only) | $0.423_{\pm 0.010}$ | $0.448_{\pm 0.006}$ | $0.584_{\pm 0.008}$ | $0.565_{\pm 0.006}$ |
| Prior Knowledge | STRING (GNN) | $0.510_{\pm 0.012}$ | $0.511_{\pm 0.010}$ | $0.671_{\pm 0.007}$ | $\mathbf{0.647}_{\pm 0.006}$ |
| | STRING (GNN)+Expr | $0.504_{\pm 0.012}$ | $0.510_{\pm 0.011}$ | $0.663_{\pm 0.010}$ | $0.637_{\pm 0.010}$ |
| | STRING (GNN)+scGPT | $0.499_{\pm 0.012}$ | $0.504_{\pm 0.009}$ | $0.659_{\pm 0.009}$ | $0.634_{\pm 0.006}$ |
| | GenePT | $0.479_{\pm 0.012}$ | $0.474_{\pm 0.010}$ | $0.638_{\pm 0.010}$ | $0.606_{\pm 0.008}$ |
| | STRING (Hop) | $0.352_{\pm 0.002}$ | $0.387_{\pm 0.003}$ | $0.512_{\pm 0.003}$ | $0.531_{\pm 0.003}$ |
| | STRING (Direct) | $0.343_{\pm 0.001}$ | $0.372_{\pm 0.002}$ | $0.515_{\pm 0.001}$ | $0.532_{\pm 0.002}$ |
| | PINNACLE | $0.338_{\pm 0.002}$ | $0.362_{\pm 0.007}$ | $0.506_{\pm 0.003}$ | $0.510_{\pm 0.005}$ |
| Cell | scGPT-Pan-Cancer | $0.442_{\pm 0.012}$ | $0.455_{\pm 0.006}$ | $0.604_{\pm 0.009}$ | $0.579_{\pm 0.008}$ |

*continued on next page*

*Table S5 (continued)*

| Group | Model | AUPRC | Macro AUPRC | AUROC | Macro AUROC |
|---|---|---|---|---|---|
| | Coexpression | $0.429_{\pm 0.009}$ | $0.466_{\pm 0.009}$ | $0.594_{\pm 0.007}$ | $0.584_{\pm 0.009}$ |
| | scGPT-Whole-Human | $0.428_{\pm 0.011}$ | $0.447_{\pm 0.008}$ | $0.591_{\pm 0.007}$ | $0.573_{\pm 0.007}$ |
| | Geneformer-V2 104M | $0.415_{\pm 0.009}$ | $0.431_{\pm 0.005}$ | $0.581_{\pm 0.007}$ | $0.563_{\pm 0.005}$ |
| | Gene2Vec | $0.404_{\pm 0.009}$ | $0.432_{\pm 0.008}$ | $0.576_{\pm 0.007}$ | $0.563_{\pm 0.008}$ |
| | Geneformer-V2 316M | $0.375_{\pm 0.011}$ | $0.407_{\pm 0.006}$ | $0.545_{\pm 0.010}$ | $0.535_{\pm 0.008}$ |
| | AIDO.Cell 3M | $0.372_{\pm 0.004}$ | $0.411_{\pm 0.006}$ | $0.542_{\pm 0.005}$ | $0.532_{\pm 0.006}$ |
| | TranscriptFormer | $0.334_{\pm 0.003}$ | $0.381_{\pm 0.007}$ | $0.500_{\pm 0.004}$ | $0.507_{\pm 0.008}$ |
| | scPRINT-2 | $0.331_{\pm 0.004}$ | $0.369_{\pm 0.006}$ | $0.496_{\pm 0.005}$ | $0.499_{\pm 0.004}$ |
| Sequence | ProTrek | $0.458_{\pm 0.010}$ | $0.466_{\pm 0.008}$ | $0.626_{\pm 0.008}$ | $0.604_{\pm 0.006}$ |
| | ESM2 | $0.430_{\pm 0.010}$ | $0.445_{\pm 0.013}$ | $0.598_{\pm 0.008}$ | $0.581_{\pm 0.008}$ |
| | HyenaDNA | $0.337_{\pm 0.003}$ | $0.375_{\pm 0.005}$ | $0.504_{\pm 0.004}$ | $0.500_{\pm 0.004}$ |
| Perturbation model | GEARS | $0.421_{\pm 0.012}$ | $0.432_{\pm 0.010}$ | $0.542_{\pm 0.003}$ | $0.560_{\pm 0.009}$ |
| Trivial / Random | Random (Noise Floor) | $0.331_{\pm 0.002}$ | $0.360_{\pm 0.011}$ | $0.497_{\pm 0.003}$ | $0.498_{\pm 0.005}$ |
| | Train Mean | $0.328_{\pm 0.004}$ | $0.373_{\pm 0.005}$ | $0.487_{\pm 0.006}$ | $0.468_{\pm 0.008}$ |
| *Cell line: HepG2* | | | | | |
| EMBER | EMBER | $\mathbf{0.533}_{\pm 0.015}$ | $\mathbf{0.530}_{\pm 0.008}$ | $\mathbf{0.682}_{\pm 0.009}$ | $\mathbf{0.649}_{\pm 0.010}$ |
| Ablations | EMBER (Static) | $0.523_{\pm 0.013}$ | $0.509_{\pm 0.009}$ | $0.678_{\pm 0.008}$ | $0.644_{\pm 0.009}$ |
| | EMBER (CSCL-Only) | $0.509_{\pm 0.009}$ | $0.511_{\pm 0.006}$ | $0.659_{\pm 0.007}$ | $0.628_{\pm 0.008}$ |
| | EMBER (Recon-Only) | $0.450_{\pm 0.012}$ | $0.466_{\pm 0.006}$ | $0.609_{\pm 0.009}$ | $0.583_{\pm 0.006}$ |
| Prior Knowledge | STRING (GNN)+Expr | $0.519_{\pm 0.016}$ | $0.512_{\pm 0.008}$ | $0.677_{\pm 0.010}$ | $0.644_{\pm 0.006}$ |
| | STRING (GNN) | $0.514_{\pm 0.018}$ | $0.506_{\pm 0.011}$ | $0.681_{\pm 0.010}$ | $0.646_{\pm 0.008}$ |
| | STRING (GNN)+scGPT | $0.510_{\pm 0.013}$ | $0.504_{\pm 0.006}$ | $0.671_{\pm 0.008}$ | $0.638_{\pm 0.007}$ |
| | GenePT | $0.447_{\pm 0.012}$ | $0.458_{\pm 0.012}$ | $0.619_{\pm 0.008}$ | $0.594_{\pm 0.006}$ |
| | STRING (Hop) | $0.344_{\pm 0.001}$ | $0.371_{\pm 0.003}$ | $0.504_{\pm 0.002}$ | $0.516_{\pm 0.003}$ |
| | STRING (Direct) | $0.336_{\pm 0.001}$ | $0.360_{\pm 0.002}$ | $0.505_{\pm 0.001}$ | $0.515_{\pm 0.002}$ |
| | PINNACLE | $0.335_{\pm 0.001}$ | $0.350_{\pm 0.004}$ | $0.502_{\pm 0.002}$ | $0.506_{\pm 0.004}$ |
| Cell | scGPT-Whole-Human | $0.454_{\pm 0.011}$ | $0.472_{\pm 0.007}$ | $0.620_{\pm 0.009}$ | $0.599_{\pm 0.009}$ |
| | scGPT-Pan-Cancer | $0.447_{\pm 0.007}$ | $0.467_{\pm 0.007}$ | $0.615_{\pm 0.007}$ | $0.593_{\pm 0.007}$ |
| | Coexpression | $0.422_{\pm 0.010}$ | $0.459_{\pm 0.005}$ | $0.591_{\pm 0.010}$ | $0.575_{\pm 0.011}$ |

*continued on next page*

*Table S5 (continued)*

| Group | Model | AUPRC | Macro AUPRC | AUROC | Macro AUROC |
|---|---|---|---|---|---|
| | Gene2Vec | $0.411_{\pm 0.011}$ | $0.443_{\pm 0.005}$ | $0.583_{\pm 0.009}$ | $0.571_{\pm 0.007}$ |
| | Geneformer-V2 104M | $0.409_{\pm 0.012}$ | $0.439_{\pm 0.008}$ | $0.580_{\pm 0.008}$ | $0.567_{\pm 0.009}$ |
| | Geneformer-V2 316M | $0.388_{\pm 0.009}$ | $0.425_{\pm 0.004}$ | $0.560_{\pm 0.010}$ | $0.548_{\pm 0.007}$ |
| | AIDO.Cell 3M | $0.382_{\pm 0.008}$ | $0.419_{\pm 0.007}$ | $0.550_{\pm 0.005}$ | $0.541_{\pm 0.005}$ |
| | TranscriptFormer | $0.342_{\pm 0.009}$ | $0.393_{\pm 0.009}$ | $0.507_{\pm 0.008}$ | $0.512_{\pm 0.007}$ |
| | scPRINT-2 | $0.331_{\pm 0.005}$ | $0.370_{\pm 0.008}$ | $0.496_{\pm 0.005}$ | $0.498_{\pm 0.007}$ |
| Sequence | ProTrek | $0.456_{\pm 0.012}$ | $0.465_{\pm 0.011}$ | $0.628_{\pm 0.008}$ | $0.604_{\pm 0.005}$ |
| | ESM2 | $0.451_{\pm 0.012}$ | $0.461_{\pm 0.012}$ | $0.617_{\pm 0.008}$ | $0.596_{\pm 0.009}$ |
| | HyenaDNA | $0.334_{\pm 0.006}$ | $0.372_{\pm 0.004}$ | $0.500_{\pm 0.008}$ | $0.500_{\pm 0.007}$ |
| Perturbation model | GEARS | $0.421_{\pm 0.008}$ | $0.455_{\pm 0.008}$ | $0.575_{\pm 0.008}$ | $0.576_{\pm 0.008}$ |
| Trivial / Random | Random (Noise Floor) | $0.334_{\pm 0.002}$ | $0.367_{\pm 0.011}$ | $0.502_{\pm 0.002}$ | $0.502_{\pm 0.007}$ |
| | Train Mean | $0.334_{\pm 0.005}$ | $0.384_{\pm 0.005}$ | $0.498_{\pm 0.008}$ | $0.463_{\pm 0.008}$ |
| *Cell line: RPE1* | | | | | |
| EMBER | EMBER | $\mathbf{0.441}_{\pm 0.013}$ | $\mathbf{0.517}_{\pm 0.006}$ | $\mathbf{0.611}_{\pm 0.009}$ | $\mathbf{0.617}_{\pm 0.006}$ |
| Ablations | EMBER (CSCL-Only) | $0.428_{\pm 0.011}$ | $0.508_{\pm 0.005}$ | $0.602_{\pm 0.009}$ | $0.609_{\pm 0.006}$ |
| | EMBER (Static) | $0.418_{\pm 0.006}$ | $0.499_{\pm 0.005}$ | $0.595_{\pm 0.007}$ | $0.603_{\pm 0.008}$ |
| | EMBER (Recon-Only) | $0.378_{\pm 0.010}$ | $0.458_{\pm 0.015}$ | $0.550_{\pm 0.011}$ | $0.554_{\pm 0.016}$ |
| Prior Knowledge | STRING (GNN) | $0.435_{\pm 0.014}$ | $0.510_{\pm 0.007}$ | $0.608_{\pm 0.007}$ | $\mathbf{0.617}_{\pm 0.005}$ |
| | STRING (GNN)+Expr | $0.435_{\pm 0.013}$ | $0.508_{\pm 0.008}$ | $0.605_{\pm 0.009}$ | $0.614_{\pm 0.008}$ |
| | STRING (GNN)+scGPT | $0.428_{\pm 0.009}$ | $0.505_{\pm 0.005}$ | $0.603_{\pm 0.008}$ | $0.612_{\pm 0.006}$ |
| | GenePT | $0.394_{\pm 0.012}$ | $0.470_{\pm 0.006}$ | $0.571_{\pm 0.008}$ | $0.580_{\pm 0.005}$ |
| | STRING (Hop) | $0.355_{\pm 0.002}$ | $0.390_{\pm 0.003}$ | $0.519_{\pm 0.003}$ | $0.539_{\pm 0.003}$ |
| | STRING (Direct) | $0.347_{\pm 0.002}$ | $0.379_{\pm 0.002}$ | $0.521_{\pm 0.002}$ | $0.537_{\pm 0.002}$ |
| | PINNACLE | $0.334_{\pm 0.000}$ | $0.339_{\pm 0.002}$ | $0.501_{\pm 0.001}$ | $0.502_{\pm 0.001}$ |
| Cell | scGPT-Pan-Cancer | $0.379_{\pm 0.009}$ | $0.462_{\pm 0.009}$ | $0.555_{\pm 0.009}$ | $0.567_{\pm 0.006}$ |
| | scGPT-Whole-Human | $0.376_{\pm 0.006}$ | $0.460_{\pm 0.007}$ | $0.551_{\pm 0.005}$ | $0.562_{\pm 0.007}$ |
| | Geneformer-V2 104M | $0.366_{\pm 0.010}$ | $0.447_{\pm 0.006}$ | $0.538_{\pm 0.008}$ | $0.548_{\pm 0.006}$ |
| | Coexpression | $0.366_{\pm 0.011}$ | $0.463_{\pm 0.010}$ | $0.541_{\pm 0.010}$ | $0.558_{\pm 0.007}$ |
| | Gene2Vec | $0.366_{\pm 0.008}$ | $0.442_{\pm 0.008}$ | $0.538_{\pm 0.007}$ | $0.546_{\pm 0.008}$ |

*continued on next page*

*Table S5 (continued)*

| Group | Model | AUPRC | Macro AUPRC | AUROC | Macro AUROC |
|---|---|---|---|---|---|
| | Geneformer-V2 316M | $0.357_{\pm 0.012}$ | $0.434_{\pm 0.010}$ | $0.528_{\pm 0.014}$ | $0.538_{\pm 0.010}$ |
| | AIDO.Cell 3M | $0.348_{\pm 0.008}$ | $0.424_{\pm 0.008}$ | $0.518_{\pm 0.008}$ | $0.526_{\pm 0.009}$ |
| | TranscriptFormer | $0.339_{\pm 0.008}$ | $0.414_{\pm 0.006}$ | $0.504_{\pm 0.009}$ | $0.511_{\pm 0.009}$ |
| | scPRINT-2 | $0.329_{\pm 0.003}$ | $0.375_{\pm 0.010}$ | $0.494_{\pm 0.004}$ | $0.498_{\pm 0.006}$ |
| Sequence | ProTrek | $0.390_{\pm 0.008}$ | $0.476_{\pm 0.009}$ | $0.567_{\pm 0.007}$ | $0.586_{\pm 0.008}$ |
| | ESM2 | $0.374_{\pm 0.008}$ | $0.458_{\pm 0.008}$ | $0.549_{\pm 0.008}$ | $0.568_{\pm 0.005}$ |
| | HyenaDNA | $0.330_{\pm 0.004}$ | $0.387_{\pm 0.011}$ | $0.493_{\pm 0.005}$ | $0.501_{\pm 0.005}$ |
| Perturbation model | GEARS | $0.371_{\pm 0.006}$ | $0.431_{\pm 0.005}$ | $0.545_{\pm 0.006}$ | $0.539_{\pm 0.007}$ |
| Trivial / Random | Random (Noise Floor) | $0.334_{\pm 0.003}$ | $0.365_{\pm 0.013}$ | $0.501_{\pm 0.005}$ | $0.501_{\pm 0.006}$ |
| | Train Mean | $0.330_{\pm 0.007}$ | $0.422_{\pm 0.006}$ | $0.491_{\pm 0.012}$ | $0.521_{\pm 0.011}$ |

**Table S6: Extended genetic-interaction benchmark on the HAP1 map.** Ranking (AUPRC and AUROC for any, negative, and positive interactions) and regression (Pearson and Spearman) performance for EMBER-GI, its ablations, and all baselines, under the unseen-library split. Values are mean$_{\pm\text{s.d.}}$ across the ten random splits. The single best value in each column is in **bold**. Higher is better for every metric.

| | | Any interaction | | Negative interaction | | Positive interaction | | Regression | |
|---|---|---|---|---|---|---|---|---|---|
| Group | Model | AUPRC | AUROC | AUPRC | AUROC | AUPRC | AUROC | Pearson | Spearman |
| EMBER | EMBER-GI | $\mathbf{0.181}_{\pm0.008}$ | $\mathbf{0.602}_{\pm0.010}$ | $\mathbf{0.099}_{\pm0.005}$ | $0.545_{\pm0.007}$ | $\mathbf{0.112}_{\pm0.005}$ | $\mathbf{0.568}_{\pm0.006}$ | $\mathbf{0.072}_{\pm0.007}$ | $\mathbf{0.050}_{\pm0.003}$ |
| Ablations | EMBER-GI (Mismatch) | $0.174_{\pm0.009}$ | $0.597_{\pm0.011}$ | $0.092_{\pm0.005}$ | $0.543_{\pm0.007}$ | $0.109_{\pm0.008}$ | $0.564_{\pm0.007}$ | $0.066_{\pm0.008}$ | $0.047_{\pm0.004}$ |
| | EMBER-GI (Static) | $0.161_{\pm0.007}$ | $0.582_{\pm0.006}$ | $0.085_{\pm0.005}$ | $0.540_{\pm0.006}$ | $0.099_{\pm0.007}$ | $0.558_{\pm0.005}$ | $0.059_{\pm0.007}$ | $0.041_{\pm0.003}$ |
| Baselines | STRING (GNN) | $0.156_{\pm0.008}$ | $0.566_{\pm0.008}$ | $0.087_{\pm0.006}$ | $0.542_{\pm0.007}$ | $0.107_{\pm0.008}$ | $0.563_{\pm0.005}$ | $0.071_{\pm0.008}$ | $0.039_{\pm0.003}$ |
| | ProTrek | $0.140_{\pm0.006}$ | $0.563_{\pm0.006}$ | $0.074_{\pm0.005}$ | $0.536_{\pm0.006}$ | $0.084_{\pm0.005}$ | $0.547_{\pm0.007}$ | $0.049_{\pm0.006}$ | $0.036_{\pm0.005}$ |
| | scGPT-Pan-Cancer | $0.140_{\pm0.008}$ | $0.571_{\pm0.008}$ | $0.071_{\pm0.004}$ | $0.534_{\pm0.013}$ | $0.076_{\pm0.006}$ | $0.542_{\pm0.011}$ | $0.041_{\pm0.010}$ | $0.037_{\pm0.006}$ |
| | GenePT | $0.139_{\pm0.006}$ | $0.554_{\pm0.007}$ | $0.074_{\pm0.004}$ | $0.532_{\pm0.006}$ | $0.089_{\pm0.007}$ | $0.547_{\pm0.006}$ | $0.056_{\pm0.005}$ | $0.033_{\pm0.002}$ |
| | Geneformer-V2 104M | $0.135_{\pm0.005}$ | $0.581_{\pm0.005}$ | $0.064_{\pm0.003}$ | $0.512_{\pm0.007}$ | $0.066_{\pm0.004}$ | $0.526_{\pm0.006}$ | $0.022_{\pm0.004}$ | $0.020_{\pm0.003}$ |
| | scGPT-Whole-Human | $0.133_{\pm0.005}$ | $0.560_{\pm0.008}$ | $0.068_{\pm0.003}$ | $0.528_{\pm0.006}$ | $0.073_{\pm0.005}$ | $0.539_{\pm0.006}$ | $0.041_{\pm0.007}$ | $0.034_{\pm0.005}$ |
| | Geneformer-V2 316M | $0.131_{\pm0.004}$ | $0.577_{\pm0.005}$ | $0.062_{\pm0.002}$ | $0.509_{\pm0.004}$ | $0.064_{\pm0.004}$ | $0.526_{\pm0.008}$ | $0.020_{\pm0.004}$ | $0.019_{\pm0.004}$ |
| | AIDO.Cell 3M | $0.130_{\pm0.005}$ | $0.567_{\pm0.005}$ | $0.061_{\pm0.003}$ | $0.505_{\pm0.005}$ | $0.061_{\pm0.003}$ | $0.517_{\pm0.007}$ | $0.013_{\pm0.003}$ | $0.016_{\pm0.003}$ |
| | Gene2Vec | $0.129_{\pm0.005}$ | $0.560_{\pm0.007}$ | $0.063_{\pm0.003}$ | $0.512_{\pm0.006}$ | $0.066_{\pm0.003}$ | $0.531_{\pm0.010}$ | $0.026_{\pm0.005}$ | $0.025_{\pm0.003}$ |
| | ESM2 | $0.128_{\pm0.005}$ | $0.549_{\pm0.007}$ | $0.068_{\pm0.002}$ | $0.535_{\pm0.008}$ | $0.075_{\pm0.004}$ | $0.547_{\pm0.005}$ | $0.047_{\pm0.007}$ | $0.039_{\pm0.005}$ |
| | STRING (Direct) | $0.117_{\pm0.005}$ | $0.563_{\pm0.009}$ | $0.060_{\pm0.003}$ | $\mathbf{0.546}_{\pm0.006}$ | $0.042_{\pm0.001}$ | $0.438_{\pm0.010}$ | $-0.004_{\pm0.001}$ | $-0.002_{\pm0.001}$ |
| | TranscriptFormer | $0.110_{\pm0.003}$ | $0.537_{\pm0.007}$ | $0.065_{\pm0.003}$ | $0.534_{\pm0.005}$ | $0.060_{\pm0.002}$ | $0.538_{\pm0.004}$ | $0.059_{\pm0.003}$ | $0.044_{\pm0.002}$ |
| | STRING (Hop) | $0.102_{\pm0.003}$ | $0.523_{\pm0.006}$ | $0.053_{\pm0.002}$ | $0.531_{\pm0.004}$ | $0.043_{\pm0.001}$ | $0.452_{\pm0.008}$ | $0.002_{\pm0.001}$ | $0.002_{\pm0.001}$ |
| | PINNACLE | $0.099_{\pm0.003}$ | $0.531_{\pm0.003}$ | $0.051_{\pm0.002}$ | $0.500_{\pm0.003}$ | $0.046_{\pm0.001}$ | $0.502_{\pm0.003}$ | $0.005_{\pm0.003}$ | $0.003_{\pm0.002}$ |
| | HyenaDNA | $0.098_{\pm0.004}$ | $0.514_{\pm0.008}$ | $0.054_{\pm0.003}$ | $0.506_{\pm0.006}$ | $0.048_{\pm0.002}$ | $0.515_{\pm0.005}$ | $0.015_{\pm0.007}$ | $0.016_{\pm0.005}$ |
| | scPRINT-2 | $0.096_{\pm0.002}$ | $0.510_{\pm0.004}$ | $0.053_{\pm0.002}$ | $0.506_{\pm0.003}$ | $0.047_{\pm0.002}$ | $0.508_{\pm0.006}$ | $0.013_{\pm0.003}$ | $0.011_{\pm0.002}$ |

**Table S7: Drug-response benchmark, per cell line.** Per-drug Pearson and Spearman correlation for EMBER-DRUG and all baselines on LINCS L1000 and Tahoe-100M, evaluated on held-out drugs. The perturbed-reference columns isolate drug-specific signal (the paper's main metric); the control-reference columns are shown for reference. Values are mean$_{\pm \text{s.d.}}$ across the ten folds. For each cell line, the best value in each column is in **bold** and the second best is underlined.

| Cell line | Group | Model | Perturbed-reference | | Control-reference | |
|---|---|---|---|---|---|---|
| | | | Pearson | Spearman | Pearson | Spearman |
| **Dataset: LINCS L1000** | | | | | | |
| *A375 (Skin)* | | | | | | |
| A375 | EMBER | EMBER-DRUG | <u>0.220</u>$_{\pm 0.047}$ | <u>0.246</u>$_{\pm 0.047}$ | 0.272$_{\pm 0.046}$ | 0.242$_{\pm 0.051}$ |
| | Ablations | EMBER (Static) | 0.160$_{\pm 0.040}$ | 0.186$_{\pm 0.042}$ | 0.266$_{\pm 0.040}$ | 0.234$_{\pm 0.043}$ |
| | Gene embedding | Geneformer-V2 104M | **0.225**$_{\pm 0.059}$ | **0.248**$_{\pm 0.060}$ | 0.283$_{\pm 0.043}$ | 0.252$_{\pm 0.045}$ |
| | | scGPT-Pan-Cancer | 0.207$_{\pm 0.049}$ | 0.231$_{\pm 0.049}$ | 0.281$_{\pm 0.043}$ | 0.250$_{\pm 0.043}$ |
| | | scPRINT-2 | 0.175$_{\pm 0.059}$ | 0.202$_{\pm 0.062}$ | 0.265$_{\pm 0.043}$ | 0.234$_{\pm 0.047}$ |
| | | GenePT | 0.165$_{\pm 0.047}$ | 0.192$_{\pm 0.046}$ | 0.268$_{\pm 0.040}$ | 0.236$_{\pm 0.045}$ |
| | | TranscriptFormer | 0.158$_{\pm 0.053}$ | 0.188$_{\pm 0.050}$ | 0.202$_{\pm 0.032}$ | 0.168$_{\pm 0.032}$ |
| | | Gene2Vec | 0.140$_{\pm 0.054}$ | 0.168$_{\pm 0.056}$ | 0.255$_{\pm 0.037}$ | 0.224$_{\pm 0.041}$ |
| | | STRING (GNN) | 0.129$_{\pm 0.060}$ | 0.152$_{\pm 0.063}$ | 0.266$_{\pm 0.041}$ | 0.233$_{\pm 0.043}$ |
| | Drug structure | Chemeleon | 0.169$_{\pm 0.049}$ | 0.200$_{\pm 0.043}$ | 0.295$_{\pm 0.039}$ | <u>0.265</u>$_{\pm 0.042}$ |
| | | Morgan | 0.168$_{\pm 0.036}$ | 0.204$_{\pm 0.035}$ | 0.281$_{\pm 0.037}$ | 0.251$_{\pm 0.040}$ |
| | | ChemBERTa | 0.134$_{\pm 0.083}$ | 0.163$_{\pm 0.087}$ | <u>0.303</u>$_{\pm 0.040}$ | **0.271**$_{\pm 0.042}$ |
| | Drug-response model | XPert | 0.123$_{\pm 0.070}$ | 0.152$_{\pm 0.069}$ | 0.275$_{\pm 0.034}$ | 0.241$_{\pm 0.037}$ |
| | Trivial | train-mean (no drug) | – | – | **0.304**$_{\pm 0.041}$ | – |
| *MCF7 (Breast)* | | | | | | |
| MCF7 | EMBER | EMBER-DRUG | **0.119**$_{\pm 0.037}$ | **0.133**$_{\pm 0.038}$ | 0.273$_{\pm 0.036}$ | 0.241$_{\pm 0.035}$ |
| | Ablations | EMBER (Static) | 0.084$_{\pm 0.036}$ | 0.097$_{\pm 0.038}$ | 0.272$_{\pm 0.039}$ | 0.239$_{\pm 0.036}$ |
| | Gene embedding | scGPT-Pan-Cancer | <u>0.117</u>$_{\pm 0.035}$ | <u>0.123</u>$_{\pm 0.035}$ | 0.286$_{\pm 0.043}$ | 0.254$_{\pm 0.041}$ |
| | | TranscriptFormer | 0.095$_{\pm 0.036}$ | 0.115$_{\pm 0.036}$ | 0.248$_{\pm 0.029}$ | 0.216$_{\pm 0.028}$ |
| | | Geneformer-V2 104M | 0.091$_{\pm 0.035}$ | 0.105$_{\pm 0.039}$ | 0.281$_{\pm 0.040}$ | 0.248$_{\pm 0.040}$ |
| | | STRING (GNN) | 0.091$_{\pm 0.042}$ | 0.097$_{\pm 0.046}$ | 0.280$_{\pm 0.040}$ | 0.249$_{\pm 0.039}$ |
| | | Gene2Vec | 0.073$_{\pm 0.042}$ | 0.086$_{\pm 0.041}$ | 0.276$_{\pm 0.043}$ | 0.243$_{\pm 0.041}$ |
| | | GenePT | 0.070$_{\pm 0.032}$ | 0.082$_{\pm 0.038}$ | 0.283$_{\pm 0.036}$ | 0.249$_{\pm 0.036}$ |
| | | scPRINT-2 | 0.070$_{\pm 0.039}$ | 0.080$_{\pm 0.039}$ | 0.285$_{\pm 0.034}$ | 0.252$_{\pm 0.034}$ |
| | Drug structure | Chemeleon | 0.088$_{\pm 0.056}$ | 0.103$_{\pm 0.046}$ | 0.312$_{\pm 0.042}$ | <u>0.280</u>$_{\pm 0.041}$ |
| | | ChemBERTa | 0.054$_{\pm 0.037}$ | 0.067$_{\pm 0.033}$ | <u>0.320</u>$_{\pm 0.039}$ | **0.285**$_{\pm 0.039}$ |
| | | Morgan | 0.044$_{\pm 0.035}$ | 0.062$_{\pm 0.029}$ | 0.313$_{\pm 0.044}$ | 0.279$_{\pm 0.043}$ |
| | Drug-response model | XPert | 0.075$_{\pm 0.040}$ | 0.099$_{\pm 0.042}$ | 0.289$_{\pm 0.047}$ | 0.256$_{\pm 0.043}$ |
| | Trivial | train-mean (no drug) | – | – | **0.327**$_{\pm 0.040}$ | – |
| *HT29 (Colon)* | | | | | | |
| HT29 | EMBER | EMBER-DRUG | <u>0.139</u>$_{\pm 0.054}$ | 0.160$_{\pm 0.049}$ | 0.290$_{\pm 0.057}$ | 0.256$_{\pm 0.058}$ |
| | Ablations | EMBER (Static) | 0.086$_{\pm 0.050}$ | 0.108$_{\pm 0.051}$ | 0.294$_{\pm 0.040}$ | 0.259$_{\pm 0.046}$ |
| | Gene embedding | scGPT-Pan-Cancer | **0.165**$_{\pm 0.048}$ | **0.182**$_{\pm 0.044}$ | 0.323$_{\pm 0.049}$ | 0.289$_{\pm 0.052}$ |
| | | TranscriptFormer | 0.138$_{\pm 0.044}$ | <u>0.165</u>$_{\pm 0.041}$ | 0.264$_{\pm 0.052}$ | 0.235$_{\pm 0.052}$ |

*continued on next page*

*Table S7 (continued)*

| Cell line | Group | Model | Perturbed-reference | | Control-reference | |
|---|---|---|---|---|---|---|
| | | | Pearson | Spearman | Pearson | Spearman |
| | | Gene2Vec | $0.119_{\pm0.054}$ | $0.140_{\pm0.058}$ | $0.287_{\pm0.050}$ | $0.253_{\pm0.052}$ |
| | | GenePT | $0.112_{\pm0.047}$ | $0.130_{\pm0.051}$ | $0.295_{\pm0.043}$ | $0.260_{\pm0.047}$ |
| | | Geneformer-V2 104M | $0.107_{\pm0.048}$ | $0.127_{\pm0.046}$ | $0.309_{\pm0.049}$ | $0.276_{\pm0.052}$ |
| | | STRING (GNN) | $0.105_{\pm0.050}$ | $0.126_{\pm0.059}$ | $0.292_{\pm0.045}$ | $0.257_{\pm0.049}$ |
| | | scPRINT-2 | $0.097_{\pm0.047}$ | $0.116_{\pm0.048}$ | $0.280_{\pm0.053}$ | $0.245_{\pm0.055}$ |
| | Drug structure | Chemeleon | $0.133_{\pm0.076}$ | $0.161_{\pm0.078}$ | $0.341_{\pm0.052}$ | $0.302_{\pm0.056}$ |
| | | ChemBERTa | $0.097_{\pm0.057}$ | $0.117_{\pm0.063}$ | $0.342_{\pm0.050}$ | $\mathbf{0.304}_{\pm0.052}$ |
| | | Morgan | $0.088_{\pm0.062}$ | $0.119_{\pm0.065}$ | $0.322_{\pm0.052}$ | $0.286_{\pm0.056}$ |
| | Drug-response model | XPert | $0.081_{\pm0.058}$ | $0.114_{\pm0.057}$ | $0.242_{\pm0.024}$ | $0.205_{\pm0.024}$ |
| | Trivial | train-mean (no drug) | – | – | $\mathbf{0.349}_{\pm0.049}$ | – |
| *A549 (Lung)* | | | | | | |
| A549 | Ember | Ember-DRUG | $\mathbf{0.138}_{\pm0.020}$ | $\mathbf{0.169}_{\pm0.022}$ | $0.216_{\pm0.038}$ | $0.183_{\pm0.035}$ |
| | Ablations | Ember (Static) | $0.124_{\pm0.030}$ | $0.151_{\pm0.032}$ | $0.234_{\pm0.035}$ | $0.194_{\pm0.031}$ |
| | Gene embedding | Geneformer-V2 104M | $0.134_{\pm0.054}$ | $0.154_{\pm0.053}$ | $0.248_{\pm0.019}$ | $0.209_{\pm0.021}$ |
| | | TranscriptFormer | $0.134_{\pm0.036}$ | $0.162_{\pm0.034}$ | $0.194_{\pm0.044}$ | $0.161_{\pm0.036}$ |
| | | scGPT-Pan-Cancer | $0.131_{\pm0.025}$ | $0.155_{\pm0.027}$ | $0.248_{\pm0.029}$ | $0.208_{\pm0.027}$ |
| | | Gene2Vec | $0.113_{\pm0.040}$ | $0.141_{\pm0.045}$ | $0.229_{\pm0.032}$ | $0.195_{\pm0.030}$ |
| | | STRING (GNN) | $0.106_{\pm0.045}$ | $0.129_{\pm0.058}$ | $0.236_{\pm0.029}$ | $0.197_{\pm0.028}$ |
| | | GenePT | $0.102_{\pm0.035}$ | $0.123_{\pm0.041}$ | $0.248_{\pm0.029}$ | $0.208_{\pm0.027}$ |
| | | scPRINT-2 | $0.084_{\pm0.037}$ | $0.107_{\pm0.043}$ | $0.227_{\pm0.027}$ | $0.190_{\pm0.029}$ |
| | Drug structure | Chemeleon | $0.125_{\pm0.071}$ | $0.154_{\pm0.074}$ | $0.279_{\pm0.032}$ | $\mathbf{0.240}_{\pm0.032}$ |
| | | Morgan | $0.101_{\pm0.056}$ | $0.136_{\pm0.053}$ | $0.269_{\pm0.024}$ | $0.231_{\pm0.024}$ |
| | | ChemBERTa | $0.097_{\pm0.058}$ | $0.128_{\pm0.064}$ | $0.278_{\pm0.025}$ | $0.239_{\pm0.028}$ |
| | Drug-response model | XPert | $0.093_{\pm0.050}$ | $0.120_{\pm0.050}$ | $0.270_{\pm0.019}$ | $0.228_{\pm0.017}$ |
| | Trivial | train-mean (no drug) | – | – | $\mathbf{0.287}_{\pm0.028}$ | – |
| **Dataset: Tahoe-100M** | | | | | | |
| *A172 (Brain)* | | | | | | |
| A172 | Ember | Ember-DRUG | $\mathbf{0.050}_{\pm0.023}$ | $0.058_{\pm0.031}$ | $0.192_{\pm0.035}$ | $0.183_{\pm0.038}$ |
| | Ablations | Ember (Static) | $0.030_{\pm0.022}$ | $0.036_{\pm0.026}$ | $0.235_{\pm0.022}$ | $0.229_{\pm0.024}$ |
| | Gene embedding | scGPT-Pan-Cancer | $0.046_{\pm0.028}$ | $0.059_{\pm0.025}$ | $0.204_{\pm0.025}$ | $0.208_{\pm0.027}$ |
| | | TranscriptFormer | $0.041_{\pm0.025}$ | $\mathbf{0.063}_{\pm0.027}$ | $0.161_{\pm0.016}$ | $0.152_{\pm0.017}$ |
| | | scPRINT-2 | $0.039_{\pm0.029}$ | $0.043_{\pm0.025}$ | $0.202_{\pm0.029}$ | $0.190_{\pm0.029}$ |
| | | GenePT | $0.039_{\pm0.026}$ | $0.045_{\pm0.025}$ | $0.231_{\pm0.023}$ | $0.224_{\pm0.024}$ |
| | | Geneformer-V2 104M | $0.038_{\pm0.023}$ | $0.050_{\pm0.023}$ | $0.202_{\pm0.020}$ | $0.210_{\pm0.021}$ |
| | | Gene2Vec | $0.035_{\pm0.025}$ | $0.045_{\pm0.024}$ | $0.217_{\pm0.024}$ | $0.201_{\pm0.025}$ |
| | | STRING (GNN) | $0.028_{\pm0.021}$ | $0.032_{\pm0.022}$ | $0.226_{\pm0.022}$ | $0.214_{\pm0.024}$ |
| | Drug structure | Chemeleon | $0.045_{\pm0.029}$ | $0.054_{\pm0.035}$ | $0.264_{\pm0.023}$ | $0.261_{\pm0.025}$ |
| | | ChemBERTa | $0.036_{\pm0.028}$ | $0.045_{\pm0.034}$ | $0.270_{\pm0.023}$ | $\mathbf{0.267}_{\pm0.024}$ |
| | | Morgan | $0.028_{\pm0.042}$ | $0.035_{\pm0.045}$ | $0.268_{\pm0.025}$ | $0.264_{\pm0.025}$ |
| | Drug-response model | XPert | $0.031_{\pm0.029}$ | $0.042_{\pm0.029}$ | $0.169_{\pm0.097}$ | $0.169_{\pm0.098}$ |

*continued on next page*

*Table S7 (continued)*

| Cell line | Group | Model | Perturbed-reference | | Control-reference | |
|---|---|---|---|---|---|---|
| | | | Pearson | Spearman | Pearson | Spearman |
| | Trivial | train-mean (no drug) | – | – | **0.275**$_{\pm 0.024}$ | – |
| *SW480 (Colon)* | | | | | | |
| SW480 | EMBER | EMBER-DRUG | **0.047**$_{\pm 0.025}$ | <u>0.054</u>$_{\pm 0.031}$ | 0.130$_{\pm 0.054}$ | 0.122$_{\pm 0.053}$ |
| | Ablations | EMBER (Static) | 0.025$_{\pm 0.023}$ | 0.029$_{\pm 0.025}$ | 0.204$_{\pm 0.029}$ | 0.187$_{\pm 0.034}$ |
| | Gene embedding | scPRINT-2 | <u>0.042</u>$_{\pm 0.013}$ | 0.051$_{\pm 0.014}$ | 0.178$_{\pm 0.030}$ | 0.164$_{\pm 0.034}$ |
| | | GenePT | 0.040$_{\pm 0.022}$ | 0.048$_{\pm 0.023}$ | 0.196$_{\pm 0.025}$ | 0.181$_{\pm 0.023}$ |
| | | Geneformer-V2 104M | 0.039$_{\pm 0.022}$ | 0.051$_{\pm 0.028}$ | 0.147$_{\pm 0.021}$ | 0.158$_{\pm 0.023}$ |
| | | TranscriptFormer | 0.038$_{\pm 0.027}$ | **0.059**$_{\pm 0.030}$ | 0.114$_{\pm 0.028}$ | 0.105$_{\pm 0.029}$ |
| | | scGPT-Pan-Cancer | 0.036$_{\pm 0.015}$ | 0.047$_{\pm 0.022}$ | 0.156$_{\pm 0.026}$ | 0.166$_{\pm 0.029}$ |
| | | Gene2Vec | 0.023$_{\pm 0.019}$ | 0.032$_{\pm 0.016}$ | 0.180$_{\pm 0.017}$ | 0.166$_{\pm 0.021}$ |
| | | STRING (GNN) | 0.010$_{\pm 0.021}$ | 0.017$_{\pm 0.028}$ | 0.191$_{\pm 0.027}$ | 0.173$_{\pm 0.030}$ |
| | Drug structure | Chemeleon | 0.018$_{\pm 0.023}$ | 0.024$_{\pm 0.031}$ | 0.223$_{\pm 0.025}$ | 0.211$_{\pm 0.024}$ |
| | | ChemBERTa | 0.014$_{\pm 0.024}$ | 0.019$_{\pm 0.030}$ | 0.229$_{\pm 0.031}$ | <u>0.217</u>$_{\pm 0.035}$ |
| | | Morgan | 0.013$_{\pm 0.029}$ | 0.019$_{\pm 0.026}$ | <u>0.232</u>$_{\pm 0.028}$ | **0.223**$_{\pm 0.032}$ |
| | Drug-response model | XPert | 0.029$_{\pm 0.039}$ | 0.041$_{\pm 0.050}$ | 0.083$_{\pm 0.105}$ | 0.085$_{\pm 0.104}$ |
| | Trivial | train-mean (no drug) | – | – | **0.251**$_{\pm 0.023}$ | – |
| *NCIH460 (Lung)* | | | | | | |
| NCIH460 | EMBER | EMBER-DRUG | **0.058**$_{\pm 0.022}$ | **0.080**$_{\pm 0.026}$ | 0.150$_{\pm 0.052}$ | 0.160$_{\pm 0.054}$ |
| | Ablations | EMBER (Static) | 0.013$_{\pm 0.025}$ | 0.025$_{\pm 0.033}$ | 0.201$_{\pm 0.028}$ | 0.198$_{\pm 0.037}$ |
| | Gene embedding | TranscriptFormer | <u>0.050</u>$_{\pm 0.030}$ | <u>0.079</u>$_{\pm 0.034}$ | 0.110$_{\pm 0.021}$ | 0.126$_{\pm 0.031}$ |
| | | Geneformer-V2 104M | 0.048$_{\pm 0.036}$ | 0.068$_{\pm 0.040}$ | 0.161$_{\pm 0.025}$ | 0.189$_{\pm 0.035}$ |
| | | scGPT-Pan-Cancer | 0.047$_{\pm 0.032}$ | 0.068$_{\pm 0.035}$ | 0.164$_{\pm 0.031}$ | 0.192$_{\pm 0.041}$ |
| | | GenePT | 0.034$_{\pm 0.026}$ | 0.052$_{\pm 0.031}$ | 0.216$_{\pm 0.026}$ | 0.218$_{\pm 0.038}$ |
| | | scPRINT-2 | 0.034$_{\pm 0.025}$ | 0.054$_{\pm 0.028}$ | 0.177$_{\pm 0.040}$ | 0.175$_{\pm 0.049}$ |
| | | Gene2Vec | 0.025$_{\pm 0.036}$ | 0.039$_{\pm 0.045}$ | 0.196$_{\pm 0.034}$ | 0.187$_{\pm 0.041}$ |
| | | STRING (GNN) | 0.019$_{\pm 0.029}$ | 0.037$_{\pm 0.038}$ | 0.205$_{\pm 0.029}$ | 0.204$_{\pm 0.040}$ |
| | Drug structure | Chemeleon | 0.027$_{\pm 0.029}$ | 0.049$_{\pm 0.050}$ | <u>0.260</u>$_{\pm 0.034}$ | **0.265**$_{\pm 0.043}$ |
| | | ChemBERTa | 0.016$_{\pm 0.027}$ | 0.033$_{\pm 0.031}$ | 0.254$_{\pm 0.029}$ | <u>0.255</u>$_{\pm 0.039}$ |
| | | Morgan | 0.006$_{\pm 0.032}$ | 0.016$_{\pm 0.041}$ | 0.250$_{\pm 0.028}$ | 0.253$_{\pm 0.037}$ |
| | Drug-response model | XPert | 0.027$_{\pm 0.035}$ | 0.059$_{\pm 0.051}$ | 0.175$_{\pm 0.115}$ | 0.187$_{\pm 0.127}$ |
| | Trivial | train-mean (no drug) | – | – | **0.270**$_{\pm 0.028}$ | – |
| *BT474 (Breast)* | | | | | | |
| BT474 | EMBER | EMBER-DRUG | <u>0.050</u>$_{\pm 0.034}$ | <u>0.055</u>$_{\pm 0.032}$ | 0.196$_{\pm 0.040}$ | 0.184$_{\pm 0.035}$ |
| | Ablations | EMBER (Static) | 0.035$_{\pm 0.020}$ | 0.034$_{\pm 0.021}$ | 0.248$_{\pm 0.023}$ | 0.228$_{\pm 0.022}$ |
| | Gene embedding | TranscriptFormer | **0.055**$_{\pm 0.040}$ | **0.065**$_{\pm 0.043}$ | 0.165$_{\pm 0.027}$ | 0.144$_{\pm 0.026}$ |
| | | scGPT-Pan-Cancer | 0.046$_{\pm 0.026}$ | 0.049$_{\pm 0.025}$ | 0.228$_{\pm 0.019}$ | 0.212$_{\pm 0.016}$ |
| | | scPRINT-2 | 0.041$_{\pm 0.018}$ | 0.046$_{\pm 0.015}$ | 0.209$_{\pm 0.024}$ | 0.193$_{\pm 0.023}$ |
| | | Geneformer-V2 104M | 0.034$_{\pm 0.022}$ | 0.043$_{\pm 0.025}$ | 0.210$_{\pm 0.021}$ | 0.202$_{\pm 0.020}$ |
| | | GenePT | 0.034$_{\pm 0.031}$ | 0.037$_{\pm 0.028}$ | 0.231$_{\pm 0.021}$ | 0.214$_{\pm 0.020}$ |
| | | Gene2Vec | 0.029$_{\pm 0.019}$ | 0.032$_{\pm 0.021}$ | 0.218$_{\pm 0.021}$ | 0.202$_{\pm 0.018}$ |

*continued on next page*

*Table S7 (continued)*

| Cell line | Group | Model | Perturbed-reference | | Control-reference | |
|---|---|---|---|---|---|---|
| | | | Pearson | Spearman | Pearson | Spearman |
| | | STRING (GNN) | $0.028_{\pm 0.016}$ | $0.030_{\pm 0.018}$ | $0.217_{\pm 0.024}$ | $0.199_{\pm 0.022}$ |
| | Drug structure | ChemBERTa | $0.032_{\pm 0.015}$ | $0.045_{\pm 0.015}$ | $\underline{0.273}_{\pm 0.020}$ | $\mathbf{0.260}_{\pm 0.018}$ |
| | | Chemeleon | $0.025_{\pm 0.020}$ | $0.039_{\pm 0.024}$ | $0.268_{\pm 0.027}$ | $0.256_{\pm 0.026}$ |
| | | Morgan | $0.017_{\pm 0.021}$ | $0.026_{\pm 0.025}$ | $0.272_{\pm 0.023}$ | $\underline{0.258}_{\pm 0.022}$ |
| | Drug-response model | XPert | $0.029_{\pm 0.028}$ | $0.041_{\pm 0.033}$ | $0.044_{\pm 0.058}$ | $0.041_{\pm 0.051}$ |
| | Trivial | train-mean (no drug) | – | – | $\mathbf{0.279}_{\pm 0.022}$ | – |
